# Worker-behavior and behavior-behavior interaction networks in the trap-jaw ant *Odontomachus chelifer* (Latreille, 1802) (Hymenoptera: Formicidae)

**DOI:** 10.1101/2020.05.25.115063

**Authors:** Felipe Marcel Neves, Marcelo Eduardo Borges, Marcio R. Pie

## Abstract

Division of labor is among the main factors to explain the evolutionary success of social systems, from the origins of multicellularity to complex animal societies. The remarkable ecological success of social insects seems to have been largely driven by ergonomic advantages stemming from the behavioral specialization of workers. However, little is known about how individuals and their corresponding behavioral repertoires are related to each other within a division of labor context, particularly by viewing such relationships as complex networks. Applications of network theory to the study of social insects are almost exclusively used to analyze behavioral interactions between individuals, rather than to the study of relations among individuals and behaviors. Here, we use an approach to the study of the organization of the behavioral repertoire of ant colonies that considers both individual-behavior interactions and behavior-behavior interactions, besides colony time budgets. Our study investigates the organization of division of labor in colonies of the trap-jaw ant *Odontomachus chelifer* (Latreille, 1802). All the behavioral acts (including inactivity) performed within three queenright colonies of different sizes (n = 7, 30, and 60 workers) were studied under controlled laboratory conditions. Each ant was individually marked and observed by scan sampling in 10 min intervals for 10 h each (n = 5,919 behavioral acts). We describe the network topologies in terms of centrality, specialization, modularity, and nestedness. This study shows that workers of *O. chelifer* are organized according to structured networks, composed of individuals exhibiting varying degrees of specialization. The observed centrality scores indicate that some behaviors could have a disproportionately larger impact on the network organization (especially self-grooming). The results underscore the potential of the use of complex networks (particularly measures of modularity and nestedness) in order to discover and study novel organizational patterns of social groups in animal behavior.

## Introduction

Division of labor is a property of social systems thought to have been favored by natural selection and that occurs in a variety of phenomena, from subcellular levels to complex animal societies (Maynard Smith & Szathmáry 1997). In particular, the remarkable ecological success of social insects seems to have been largely driven by the ergonomic advantages stemming from division of labor, such as individual specialization and parallel task execution (Oster & Wilson 1978, Robinson 1992, Traniello & Rosengaus 1997, Beshers & Fewell 2001, Rocha & al. 2014, Avril & al. 2016). Division of labor in social insects can be broadly defined as “any behavioral pattern that results in some individuals in a colony performing different functions from others” (Michener 1974). Uncovering general principles of division of labor requires the assessment of the contribution from individual workers toward the completion of colony tasks (Beshers & Fewell 2001). Therefore, insights about the organization of the division of labor in social insects might be obtained through the view of the relationship between workers and their corresponding behavioral performance as complex networks.

A complex network is described as a system of interacting nodes (within a social insect colony, workers or behaviors connected by links) that communicate with each other, displaying patterns of connection that are neither purely regular nor purely random (Newman 2003). In social insects, the network concept itself has been present in the literature for a long time, especially considering the interaction between individuals, i.e., social interactions (Gordon 2010). Even then, empirical data that explicitly describe interaction networks in social insect colonies have been only recently studied (e.g., Naug 2008, Bhadra & al. 2009, Naug 2009, Sendova-Franks & al. 2010, Pinter-Wollman & al. 2011, Waters & Fewell 2012, Mersch & al. 2013). While social interaction networks are an interesting approach to understand the behavioral influence of one individual on another, it does not reflect explicitly how workers interact with the behaviors performed by them. In the literature there are several theories and evidence regarding the division of labor exhibited by ant colonies, yet there is still a need for studies that reflect the empirical complexity exhibited by the interactions of workers and behavioral acts within a colony. This lack of knowledge may hinder the understanding of the organization of division of labor, and even our ability to assess the extent to which the mechanisms proposed to explain division of labor are mutually consistent. In order to fill this gap, one could use a framework considering networks in different ways. For instance, we could envision behavioral acts as nodes and individual workers as links (such as symbolic dynamics) (Fewell 2003). Charbonneau & al. (2013) presented one empirical analysis of task-task interaction networks (unipartite networks) in order to identify dynamics of task switching within a colony of the ant *Temnothorax rugatalus* (Emery, 1985). In this work, they used betweenness centrality to measure how central is the role of one task compared to others. The results showed that ants walk throughout the nest between tasks rather than directly switching among them. Recently, Pasquaretta & Jeanson (2018) proposed the use of bipartite networks to represent interactions between workers and tasks. Moreover, the authors quantified the bipartite network in the context of division of labor, using specialization and modularity measures, which consider the specialization of nodes and the strength of division of a network into groups, respectively.

Here, we use an approach that goes beyond simple social interaction networks by considering both worker-behavior interactions and behavior-behavior interactions, besides usual colony time budgets. Such an integrative approach offers complementary results, showing possibly hidden dynamics in the formation of complex behavioral interactions. Besides the network metrics used in the previously mentioned studies, we also considered theoretical concepts developed in the context of community ecology, such as nestedness, and expanded the use of modularity for behavior-behavior interactions. A nested network structure is observed when specialists mainly interact with proper subsets of the nodes of generalists. Nested networks are generally robust against random node loss (i.e., loss of workers or behaviors, depending on the type of network) (Thébault & al. 2010), while networks with a high degree of specialization are more vulnerable (Kaiser-Bunbury & al. 2017). Regarding modularity in behavior-behavior networks, the division of the network in groups that are more related to each other (modules) is similar to the concept of roles developed by Hölldobler & Wilson (1990), stated as a “set of behavioral acts, linked by relatively high transition probabilities”. Inactivity may occur as a result of time delays associated with searching for or switching tasks (Leighton & al. 2017). Thus, we also use a variation of the behavior-behavior network which considers inactivity frequency between the behavioral acts as links to quantify its influence.

Our study uses the trap-jaw ant *Odontomachus chelifer* (Latreille, 1802) as a model organism. We investigate the organization of division of labor in colonies of *O. chelifer*. In the congener *Odontomachus brunneus* (Patton), dominant individuals are more likely to reside in the central areas of the nest, where they take care of the brood, while subordinate individuals are pushed towards the edges, where they are more likely to forage (Powell & Tschinkel 1999). This process of division of labor has been called “Interaction-based task allocation” (Powell & Tschinkel 1999). Similarly, the division of labor in *O. chelifer* could be based on interactions between workers resulting in spatial fidelity and we take this into account. Thus, we study the organization of division of labor in *Odontomachus chelifer* as complex networks. In particular, we describe the individual influence of behaviors and workers through different network analysis (i.e., specialization, centrality, modularity and nestedness) applied to behavior-worker and behavior-behavior networks at the same time.

## Methods

### Field Collection and Culture Methods

The species chosen for this study is the ant *Odontomachus chelifer* (Latreille, 1802). The genus *Odontomachus* (Ponerinae) is characterized by large body size (≈ 12-15 mm in length) and a powerful articulated jaw, and usually forms small colonies (Latreille 1804, Patek & al. 2006, Spagna & al. 2008). The species is distributed from Mexico to the northeast of Argentina (Brown 1976) and has a generalist diet (Raimundo & al. 2009, Núñez & al. 2011). In this study, five colonies of *Odontomachus chelifer* were collected in forest fragments at the campus of the Universidade Federal do Paraná and the Museu de História Natural do Capão da Imbuia, Curitiba, Paraná, Brazil. The field-collected colonies usually had less than 100 workers (colonies with 75, 60, 30, 34 and 7 workers; mean ± sd: 41 ± 26). In the laboratory, colonies were transferred to artificial plaster nests, where they were kept under stable environmental conditions (20°C under constant light with 600≈lux and humidity at 60%).

Internal dimensions of the cavity were 19.5 x 15 x 2 cm (width x depth x height) divided in two chambers. All the colonies were supplied with water *ad libitum*, pieces of mealworms and artificial diet (Bhatkar & Whitcomb 1970). In all colonies, ants arranged themselves in a pattern similar to that observed by Powell & Tschinkel (1999) for *O. brunneus*, where the chamber furthest from the entrance contained the brood and queen, creating three distinct zones: the “brood zone”, “broodless zone” (all the other areas within the nest), and “foraging zone” (area outside the nest). Colonies were allowed to adjust to laboratory conditions for one month before the beginning of observations. All workers were marked individually with combinations of oil-based Testors^®^ paint (one spot on the head, one on the mesosoma, and two on the gaster) for individual worker recognition. All colonies were allowed to adjust to laboratory conditions for at least one month before any focal workers were marked, and the colonies were left for an additional week after marking before observations began. The colony size was not altered for the experiments. Voucher specimens of the ants from each colony studied were deposited in the collection of the Department of Zoology, Univerrsidade Federal do Paraná (DZUP), Curitiba, Brazil. The species was morphologically identified following the key of Brown (1976).

### Behavioral observations

Three queenright monogynous colonies of the five collected in the field were chosen for observation based on their apparent health and status (i.e., a large brood pile and the presence of a queen). Each worker from the three colonies (n = 60, 30, and 7 workers, henceforth colonies A, B, and C, respectively) was observed through scan sampling at 10-min intervals for ten hours, divided into two observation sessions by an interval of two days (five hours each; between 9:00 and 19:00 *per* trial). In each trial, the zone chambers were systematically scanned, noting the behavioral state in order to ensure the correct behavioral notation of all the ants. The observations were recorded with a digital camcorder (JVC GZ-HM320SUB) placed above the colonies. After the videos were analyzed, all the recorded behaviors (5820 recorded activities) were double-checked by a second person to ensure accurate recordings of ant identities across the observations (Data S1 A-C, as digital supplementary materials to this article, at the journal’s web pages). Individual behavioral repertoires were created (see Tab. 1 for a complete list of the behavioral acts and definitions; Script S1 and Data S2, as digital supplementary material to this article, at the journal’s web pages) Behavioral acts were classified as either ‘active’ (antennation, brood care, carrying brood, carrying debris, carrying food, feeding, foraging/patrolling, grooming, being groomed, self-grooming and walking) or ‘inactive’ (immobile and not otherwise engaged in any active task).

**Tab. 1:** List of possible behavioral acts observed in the colonies (divided in two classes; active and inactive) of *Odontomachus chelifer* (Latreille, 1802), the acronym used in the figures and tables, and descriptions of each behavior.

| Class | Behavioral acts | Acronym | Description |
| --- | --- | --- | --- |
| Active | Antennation | at | Contact with another worker with the antenna |
|  | Brood care | bc | Manipulating brood (feeding, grooming) |
|  | Carrying brood | cb | Moving brood |
|  | Carrying debris | cd | Carrying/manipulating a stone within the nest in any way (moving, pushing, pulling) |
|  | Carrying food | cf | Manipulating food inside and outside the nest |
|  | Feeding | fd | Feeding inside nest (brought back by foragers) |
|  | Foraging/Patrolling | fp | Located outside of the nest (foraging or patrolling) |
|  | Grooming | g | Grooming another ant |
|  | Being groomed | bg | Be groomed by another ant |
|  | Self-grooming | sg | Grooming itself |
|  | Walking | wl | Walking within the nest |
| Inactive | Inactivity | in | When the ant is immobile within the nest and not engaged in “any” active behavior (more than 10s) |

### Networks

Networks are depictions of adjacency matrices, in which an element within the matrix *a_ij_* with a value equal to zero means the absence of interaction, and any value ≥ 1 indicates the number of interactions between the elements of the network (Data S3-16 & Script S1-6, as digital supplementary materials to this article, at the journal’s web pages). Two different kinds of networks were considered in this study: worker-behavior networks (WBNs) and behavior-behavior networks (BBNs). WBNs characterize the relationship between two sets of nodes; workers and their respective behavioral repertoires. It is an undirected bipartite graph, in which links (edges) are defined whenever an individual performs a specific behavior. Alternatively, BBNs connect every behavior to the one performed immediately after it, with workers as links. It is a unipartite di-graph, representing temporal and directional interaction between each behavior. Networks were analyzed considering all the behaviors observed as nodes, excluding inactivity behavior. Inactivity was excluded as a node because its disproportionately high prevalence obscured other network patterns. Nevertheless, the influence of inactivity was quantified in two ways. The first one was calculated as the normalized proportion of the raw inactivity for each ant (named *Ii* index) during behavioral observations. The second one is a variation of the BBNs, where behaviors were linked by the frequency of inactivity behaviors between them (i.e., BBNs of inactivity). Thus, the link between two nodes (tasks) of the BBNs of inactivity was quantified as the presence (1) or absence (0) of the behavioral interaction, and inactivity was the additional (1 + *n*) or the only weight of the link.

### Network metrics

A series of measures were used from network analysis in the WBNs and BBNs, all analyses were performed using R 3.6.0 (R Development Core Team 2019). Graph visualization was created using both the igraph version 1.2.5 and bipartite version 2.15 packages (Csardi & Nepusz 2006, Dormann & al. 2009). The organization within the networks was explored, i.e., the roles of individual nodes (node-level metrics), as well as on a global scale (network-level metrics). Each metric had its values compared to the ones obtained from random networks generated by a specific null model (each null model used in our study is explained in the section: Statistical analysis and comparison with nulls models). The chosen metrics could be divided into five categories: Specialization (only for WBNs), centrality (only for BBNs), modularity and nestedness (for both WBNs and BBNs).

#### 1) Specialization

Two specialization network measures based on interaction frequencies in the WTNs were used: the *d’* and *H_2_’* metrics (Blüthgen & al. 2006), which represent scale-independent indices to characterize specialization in ecological networks at node and group-levels, respectively (Data S17-19 as digital supplementary materials to this article, at the journal’s web pages). Originally, both measures were proposed to quantify specialization in ecological plant-pollinator networks. The *d’* index is derived from the Kullback-Leibler distance (such as Shannon’s diversity index) and quantifies how strongly a behavior (or worker) deviates from a null model which assumes behavioral allocation in proportion to the workers and behaviors available (more details; Blüthgen & al. 2006). The *d’* index ranges from 0 (no specialization) to 1 (full specialization) and can be calculated at worker level (*d’_indv_*) or behavior level (*d’_behavior_*). For the entire network, the degree of specialization considering both parties (e.g. behaviors and workers) can be determined with the *H’_2_* index (Blüthgen & al. 2006, 2008). *H’_2_* was used in the context of division of labor for the first time by Pasquaretta & Jeanson (2018). It describes to which extent the worker-behavior interactions deviate from those that would be expected from a neutral configuration given the workers and tasks marginal totals. *H’_2_* ranges between 0 (no specialization) to 1 (complete specialization). The *d’* and *H_2_’* measures were calculated by the R package bipartite version 2.15 in R (Dormann & al. 2008). Furthermore, Gorelick et al. (2004) created two indices based on normalized mutual entropy, which have been used in several empirical and theoretical studies of division of labor (e.g., Jeanson & al. 2007, Dornhaus 2008, Santoro & al. 2019). While they were not created in the context of network theory, they are implemented in adjacent matrices such as graphs. These metrics quantify specialization from individuals and behaviors and were named DOL_indv_ and DOL_behav_ (also known as DOL_task_), respectively. The two indices range between 0 (no division of labor) to 1 and were indirectly compared to the *H_2_’* index (because as the *H_2_’* index, they are calculated for the entire network).

#### 2) Centrality

Betweenness centrality and degree centrality were used to study the patterns of flow information across the behavioral acts (BBNs). Betweenness centrality is a measure of how often a node is located on the shortest (geodesic) path between other nodes in the network (Freeman 1979). Thus, it measures the degree to which the node (behavior) functions as a potential point of control of communication (i.e., bridge) among the other nodes within a network. In unweighted networks (where the original betweenness centrality was proposed), all links have the same weight, thus the shortest path for interaction between two nodes is through the smallest number of intermediate nodes. Differently, most of the new centrality measures proposed for weighted networks have been solely focused on edge weights, and not on the number of links, a central component of the original measure. Due to this issue, the betweenness centrality proposed by Opsahl & al. (2010) was used, which considers both the number and the strength of links (weight). The relative importance of these two aspects in the metric is controlled by the tuning parameter (α), which goes from 0 to 1. Alpha was set to 0.5 to consider both factors with the equal proportions (Script S7-9 as digital supplementary materials to this article, at the journal’s web pages). In order to differentiate nodes with higher betweenness centrality from the others, tasks with a betweenness centrality above the third quartile of the data (>75%, i.e. two or three behavioral acts with the highest scores) were considered bridges. After the identification of the bridges according to betweenness centrality, we analyzed if the frequency of behavior switches between behavioral acts was random in comparison with a uniform discrete distribution of the data (i.e. equally distributed frequency among the behaviors). This information could show if the existence of a bridge is an epiphenomenon due to its higher frequency compared to other behaviors or if it is the result from selectiveness towards some specific behaviors (Data S20 and Script S10 as digital supplementary materials to this article, at the journal’s web pages). Degree centrality was applied to the BBNs of inactivity and used to describe the latency of the activity among the tasks, i.e., the higher the degree centrality of a node, the higher the latency (inactivity) around it. Originally, degree centrality is simply the count of how many connections (i.e., links) a node has in a binary network. The degree has generally been extended to the sum of weights in weighted networks (Barrat & al. 2004, Newman 2004, Opsahl & al. 2008) and labeled node strength. In order to combine both degree and strength, the degree centrality metric proposed by Opsahl & al. (2010) was utilized, which, as the betweenness centrality proposed by the same authors, uses a tuning parameter (α) to set the relative importance of the number of ties compared to link weights. The α tuning parameter was set to 0.5 to consider both factors with equal proportions. Degree centrality was divided as in and out-degree centrality for directed graphs (such as BBNs of inactivity). As the names imply, in-degree point toward and out-degree away from the given node. Behavioral acts with an in-degree and out-degree centrality above the third quartile of the data (> 75%, i.e. two or three behavioral acts with the highest scores) were regarded as inactivity hubs (i.e., with inactivity converging to the node as a link) or inactivity spreaders (i.e., with inactivity leaving the node as a link), respectively. While the centrality measures for all observed behaviors were computed and compared with a null model (Tab. 2, 3 and 4), in this study only behaviors considered bridges, inactivity hubs and spreaders were reported (Script S11-13 as digital supplementary materials to this article, at the journal’s web pages). Betweenness centrality and degree centrality were calculated using the R package tnet version 3.0.16 (Opsahl 2009).

**Tab 2.** Betweenness centrality values from the BBNs (behavior-behavior networks) of all the colonies (A, B and C) of *Odontomachus chelifer* (Latreille, 1802). The behaviors (nodes) of all workers are considered. The behavioral acts are considered bridges if they have a betweenness centrality value above the third quartile of the data (>75%; two or three behavioral acts with the highest scores, signaled in bold). The *P*-values of the Z-scores (expressed as significant in bold) of the data compared to the null model (link and weight reshuffling null model) are also exposed.

| Behavioral acts | A | *P*-values | B | *P*-values | C | *P*-values |
| --- | --- | --- | --- | --- | --- | --- |
| wl | **14** | 0.13 | 0 | 0.53 | 5 | 0.23 |
| sg | **36** | 0.36 | **17** | 0.29 | 0 | 1 |
| g | 0 | 1 | **8** | 1 | --- | ----- |
| bg | 0 | 1 | 5 | 0.88 | --- | ----- |
| bc | **14** | **0.05** | 2 | 0.99 | 0 | 1 |
| cb | 0 | 1 | 3 | 0.07 | --- | ----- |
| fd | 0 | 1 | --- | ----- | **6** | 0.35 |
| cf | 0 | 1 | --- | ----- | 0 | ----- |
| fp | 0 | 1 | 0 | 1 | 5 | 0.20 |
| cd | 0 | 1 | --- | ----- | --- | ----- |
| at | --- | ----- | 0 | 1 | 0 | 1 |

#### 3) Modularity

Modularity was proposed by Newman (2006) to compute the strength and number of modules within a network, and it has been studied across different biological scales (Lorenz & al. 2011). Modules can be defined as groups of tightly connected nodes that are sparsely connected to other nodes in the network (Newman 2006). The modularity (*Q*) ranges from 0 (community structure not different from random) to 1 (complete separation between modules) (Data S21-22 & Script S2 as digital supplementary materials to this article, at the journal’s web pages). There are different algorithms available to detect modules in weighted bipartite and unipartite networks (Clauset & al. 2008, Dormann & Strauss 2014, Beckett 2016). In the WBNs, the DIRTLPAwb+ algorithm for optimizing bipartite modularity was used (Beckett 2016) and implemented in the R package bipartite version 2.15 (Dormann & al. 2008). We normalized the bipartite modularity values following Pasquaretta & Jeanson (2018). The algorithm used to search for modules in the BBNs is the Louvain method developed by Clauset & al. (2008) and implemented in the R package igraph version 1.2.5 (Csardi & Nepusz 2006) (Script S14-15 as digital supplementary materials to this article, at the journal’s web pages).

#### 4) Nestedness

Two different metrics to estimate nestedness of the WBNs and BBNs were used (Data S23-27 & Script S2-6 as digital supplementary materials to this article, at the journal’s web pages). The first metric was the weighted nestedness metric based on overlap and decreasing fill (WNODF), which is a modified version of the nestedness metric based on overlap and decreasing fill (NODF) which considers weighted bipartite networks instead of only binary ones (Almeida-Neto & al. 2008, Almeida-Neto & Ulrich 2011). WNODF nestedness score ranges from 0 (non-nested) to 100 (perfectly nested) and it was applied to the WTNs. The nestedness in the BBNs was quantified by the UNODF, the unipartite version of the NODF metric (Cantor & al. 2017). In completely non-nested networks, UNODF = 0, while in perfectly nested networks UNODF tends towards 1. Directed networks (such as the ones of this study) will have two different UNODF values (and interpretations), because the interactions in matrix elements *a_ij_* and *a_ij_* represent different things. These two different UNODF values could be divided in nestedness among rows (UNODFr) and nestedness among columns (UNODFc). UNODFr measures nestedness computing the pairwise overlap among rows and UNODFc the pairwise overlap among columns. Since the calculation of UNODF is made through binary networks, the UNODF index was measured for different cut-off values (such as Cantor & al. 2017). The metric was calculated without a cut-off to include all data (named UNODF 1), but also considering a cut-off of 10% of the data (named UNODF 2), in order to exclude behavioral acts which were not so frequent considering all others. WNODF and UNODF were calculated using the R packages bipartite version 2.15 and unodf version 1.2, respectively (Dormann & al. 2009, Cantor & al. 2017).

### Statistical analysis and comparison with null models

G-tests of goodness of fit (including post-hoc pairwise comparisons) and G-test of independence tests were used to compare the frequency of behavioral acts (Sokal & Rohlf 1981). In the G-test goodness of fit test, the null hypothesis considers that the number of observations in each behavioral act is equal to that predicted by a uniform discrete distribution of the data, and the alternative hypothesis is that the observed numbers differ from this expected distribution. The p-values of the multiple comparisons were adjusted to control the false discovery rate using the Benjamini-Hochberg procedure (Benjamini and Hochberg 1995). Correlation between *d’* and *Ii* was tested (Spearman correlation). There are several null models to generate random networks, from simpler to more sophisticated (Farine 2017). The Paterfield’s algorithm was used in the WBNs, as suggested by Pasquaretta & Jeanson (2018) for the division of labor for bipartite networks. This model generates random networks constraining the marginal sums (i.e., worker performance and behavior need are maintained), but links are randomly assigned between workers and behaviors. Since a model for the *Ii* index could not be obtained through random networks, for this specific metric of WBNs, its empirical distribution per colony was compared to a null model with a continuous normal distribution originated from the empirical data. In this model, values outside the empirical interquartile range were excluded, thus identifying individuals or behavioral acts that have *Ii* index values outside this range. There is no null model recommended in the literature for unipartite networks (BBNs) in a division of labor context, thus we considered three different null models extensively used in the literature and that had important properties to be considered in our work. To compare the values of modularity and betweenness centrality obtained in the original BBNs with those obtained from random networks, the link and weight reshuffling null model developed by Opsahl & al. 2008 was considered. Such model consists of reshuffling the network topology while preserving the degree distribution. The importance of maintaining the network degree distribution is that most real-world degree distributions are naturally skewed rather than having a uniform or Poisson distribution. Thus, preserving the same degree distribution of the original network makes the null model more realistic and comparable to the original network. In order to compare degree centrality, the weight reshuffling null model Opsahl & al. (2008) was considered. The weight reshuffling procedure consists of reshuffling the weights globally in the network (Opsahl & al. 2008). This null model maintains the topology of the observed network. Therefore, the number of ties originating from a node does not change. The null models used to verify nestedness were first developed for bipartite networks, so an adapted version of a model widely used in bipartite biological networks, named null model 2 (Bascompte & Jordano 2007) for unipartite binary networks (i.e., modified versions of the weighted BBNs; Cantor & al. 2017) was used. In this null model, the probability that a link connects two nodes is proportional to their corresponding degree. A conveniently property of this model is that it preserves key network features, such as the network size, connectance and degree distribution. The statistical significance of all the network measures compared to the random networks was evaluated based on the *Z*-score:

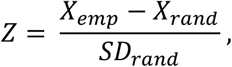

where *X*_emp_ is a metric extracted from the empirical networks (n = 1000), *X*_rand_ is the average value of the same metric obtained from random networks and SD_rand_ is the standard deviation of the metric obtained from the randomized networks. The *p* values from all the analyses were considered significant at *p* ≤ 0.05, but also described at different levels as well when necessary (i.e., ≤ 0.01 and ≤ 0.001). All the statistical analyses and null models were performed by the R software 3.6.0 (R Core Team 2019). The R packages rvaidememoire version 0.9-77 and stats version 4.0.2 were used to compute the G-test and Spearman correlation, respectively (R Core Team 2019, Hervé 2020). The null models for WBNs were created by the R package bipartite version 2.15 (Dormann & al. 2008). Two packages were used to create null models for BBNs, the TNET package version 03.0.16 was used to create the link and weight reshuffling model, and the weight reshuffling model (Opsahl 2009), the unodf package version 1.2 was used to create the adapted null model 2 (Cantor & al. 2017).

## Results

We analyzed the behavioral data in two main forms: as behavioral repertoires (i.e. colony time budgets) and as the inferences obtained from network analyses (WBNs and BBNs). The data was composed of 5,919 behavioral acts (including inactivity), WBNs resulting in 1,195 interactions, and BBNs resulting in 7,706 interactions.

### Behavioral repertoire

The behavioral repertoire performed by the workers was composed of 12 behavioral acts, considering inactivity (Fig. 1). The frequency of inactivity compared to the other behavioral acts was significantly higher across all colonies. Over the observation period, 81% of workers were inactive in colony A (*G*_(59)_ = 650; *P* < 0.001), 83% in colony B (*G*_(29)_ = 243; *P* < 0.001) and 61% in colony C (*G*_(6)_ = 26.9; *P* < 0.001) (Fig. 1). Among the behavioral acts, walking (≈28%) and self-grooming (≈24%) had frequencies significantly higher considering all the colonies (colony A, *G*_(10)_ = 747.03, *P* < 0.001; colony B, *G*_(8)_ = 233.96; *P* < 0.001; colony C, *G*_(6)_ = 138.78; *P* < 0.001). Brood care also had a higher frequency than the other behaviors but limited to the colony A and B (≈21) (Fig. 1). Moreover, we observed rare behaviors that could be characterized as dominance interactions, however, so few aggressive interactions were observed (n < 10; outside the time interval of the scan sampling) that they were not analyzed here, the dominance interactions observed for *O. chelifer* are very similar to those described by (Powell & Tschinkel 1999). It should be noted that brood care and carrying brood behavioral performance were only observed in the brood zone.

**Fig. 1:**
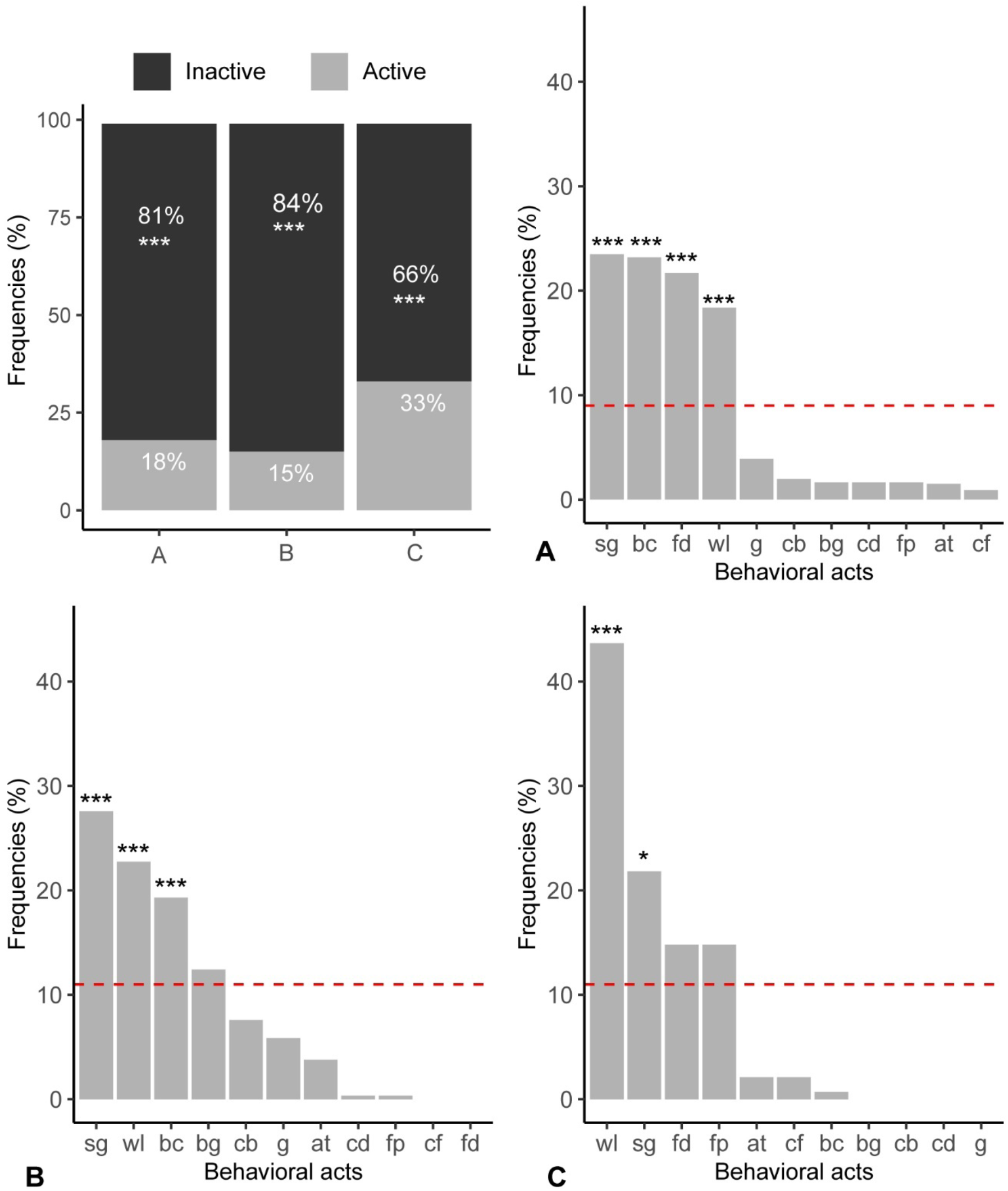
Frequencies of the behavioral acts observed in the colonies (A, B and C) of *Odontomachus chelifer* (Latreille, 1802). The stacked barplot in the right corner is representing the frequencies of inactive and active behavior classes, the other barplots are representing the frequencies among the active behavioral acts. The red dashed line is representing the value of the null model under a uniform distribution of the data. The significant statistical differences (post-hoc G-test) were signaled by *, ** and *** (*p* values less than 0.05, 0.01 and 0.001, respectively).

### Specialization

We found that *H_2_’* values in the colonies were significantly higher than the *H_2_’* obtained from random networks (Fig. 2 and 3, Colony A, *Z* = 29.3, *P* < 0.0001; Colony B, *Z* = 15.8, *P* < 0.0001; Colony C, *Z* = 10, *P* < 0.0001). The DOL_indv_ values in the colonies (Fig. 3) were also significantly higher than the correspondent ones obtained from random networks (Colony A, *Z* = 28.9, *P* < 0.0001; Colony B, *Z* = 15.4, *P* < 0.0001; Colony C, *Z* = 9.8, *P* < 0.0001). Following the same trend, DOL_behavior_ values (Fig. 3) were higher than those from random networks (Colony A, *Z* = 3.2, *P* < 0.0001; Colony B, *Z* = 2.8 *P* = 0.005; Colony C, *Z* = 9.8, *P* = 0.001). The distribution of *d*’_behavior_ and *d*’_indv_ values were clearly skewed, with similar median values between the colony A and B, and lower values in colony C. All the behaviors from each colony had a *d*’_behavior_ higher than then ones obtained from random networks (Fig. 4, *Z*-score, *P* < 0.05). Brood care (Colony A and B) and foraging/patrolling (Colony C) had the highest *d’*_behavior_ values among the behavioral acts and both were significantly higher than its respective ones from random networks (Fig. 4, *Z*-score, *P* < 0.05). Some individuals (23% in the colony A, 26% in the colony B and 28% in the colony C) had a *d*’_indv_ significantly higher than *d*’_indv_ obtained from random networks (Fig. 4, *Z*-score, *P* < 0.05). Moreover, a few individuals (25% in the colony A, 26% in the colony B and 14% in the colony C) had an *Ii* index significantly higher than *d*’_indv_ obtained from the null model (Fig. 4, *Z*-score, *P* < 0.05). There was no observed correlation between the *d*’_indv_ and the correspondent *Ii* index from each colony (Colony A, *r*_s_: −0.03, *P* = 0.76; Colony B, *r*_s_: −0.35, *P* = 0.06; Colony C, *r*_s_: 0.57, *P* = 0.10).

**Fig. 2:**
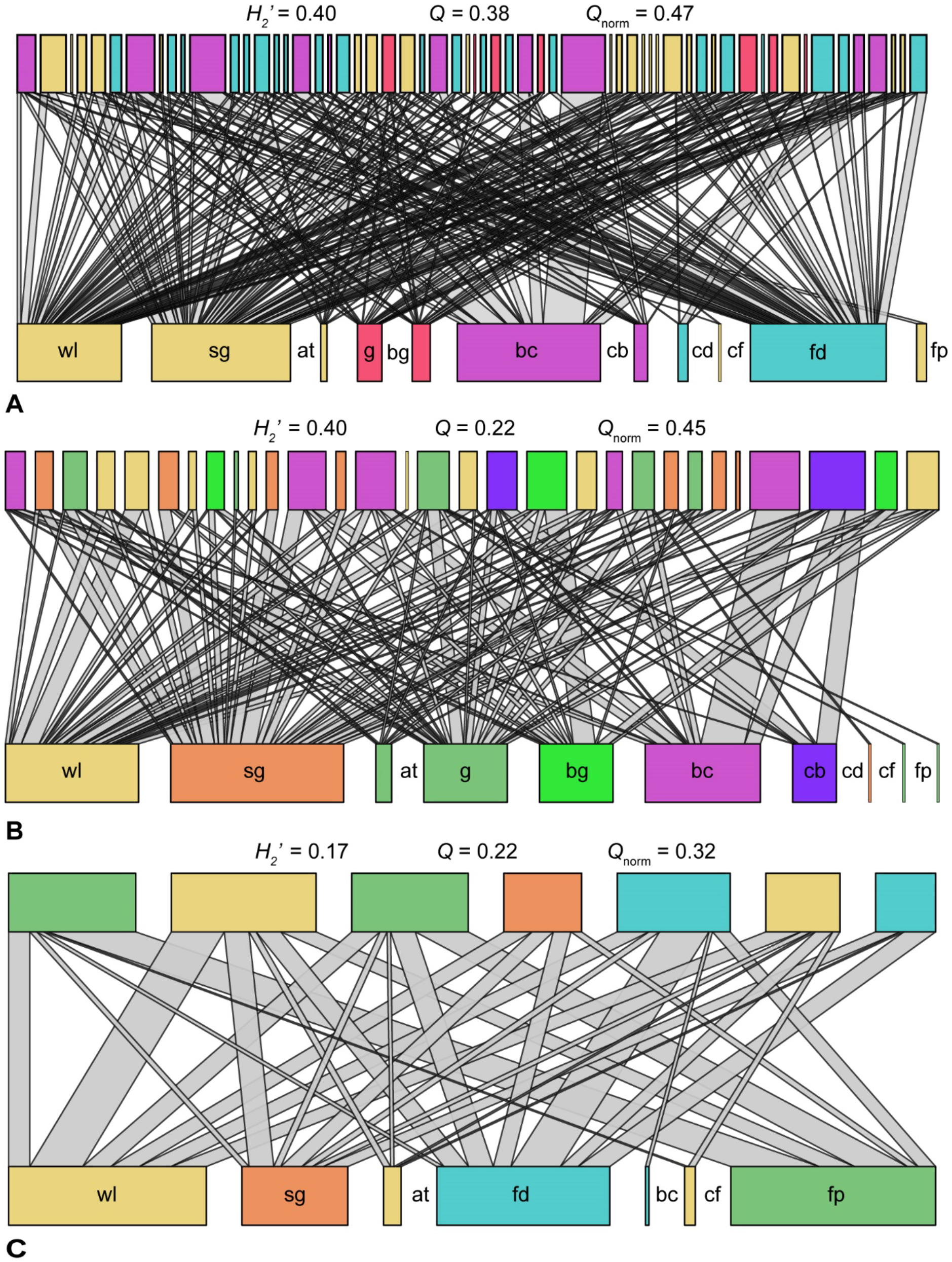
WBNs (worker-behavior networks) graphs observed from the colonies (A, B and C) of *Odontomachus chelifer* (Latreille, 1802). Upper and lower rectangles represent workers and behavioral acts, respectively. The width of each rectangle is proportional to the number of acts and the width of link indicates the frequency of interactions between behaviors and workers. For each network, numbers in upper rectangles represent worker identities. For each network, the value of *H’_2_* (specialization index), *Q*, and *Qnorm* (modularity indices) are given. The different modules of workers and behaviors are identified in different colors.

**Fig. 3:**
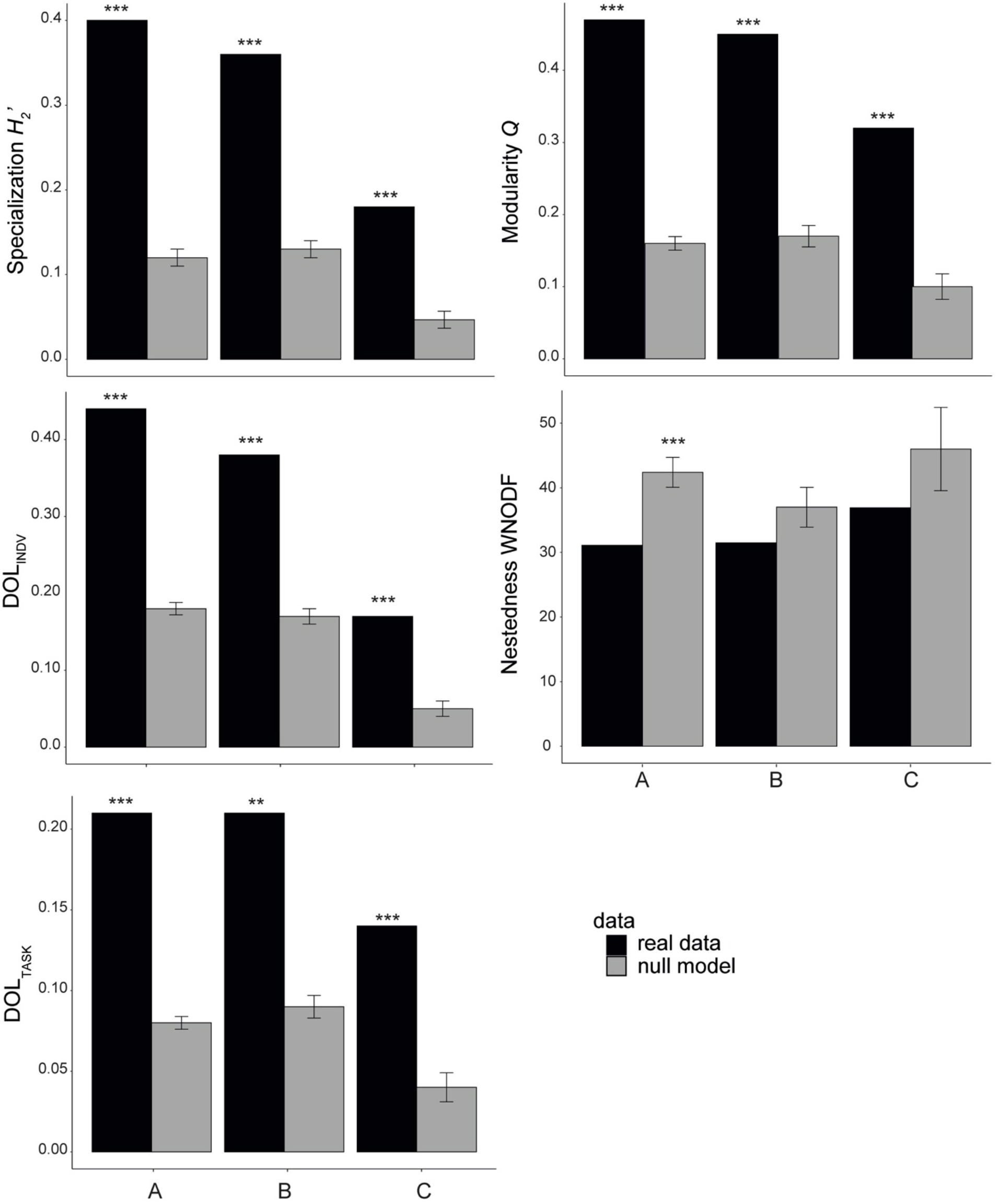
Network specialization (*H_2_’*, DOL_indv_, DOL_task_), modularity (*Q*) and weighted nestedness (WNODF) for WBNs (worker-behavior networks) from the colonies (A, B and C) of *Odontomachus chelifer* (Latreille, 1802). Black bars represent the original networks, while grey bars represent networks randomized and the respective standard deviation (SD). The significant statistical differences (Z-score) were signaled by *, ** and *** (*p* values less than 0.05, 0.01 and 0.001, respectively). The asterisks (*) above the bars mean significant differences between the original and the randomized networks or vice and versa.

**Fig. 4:**
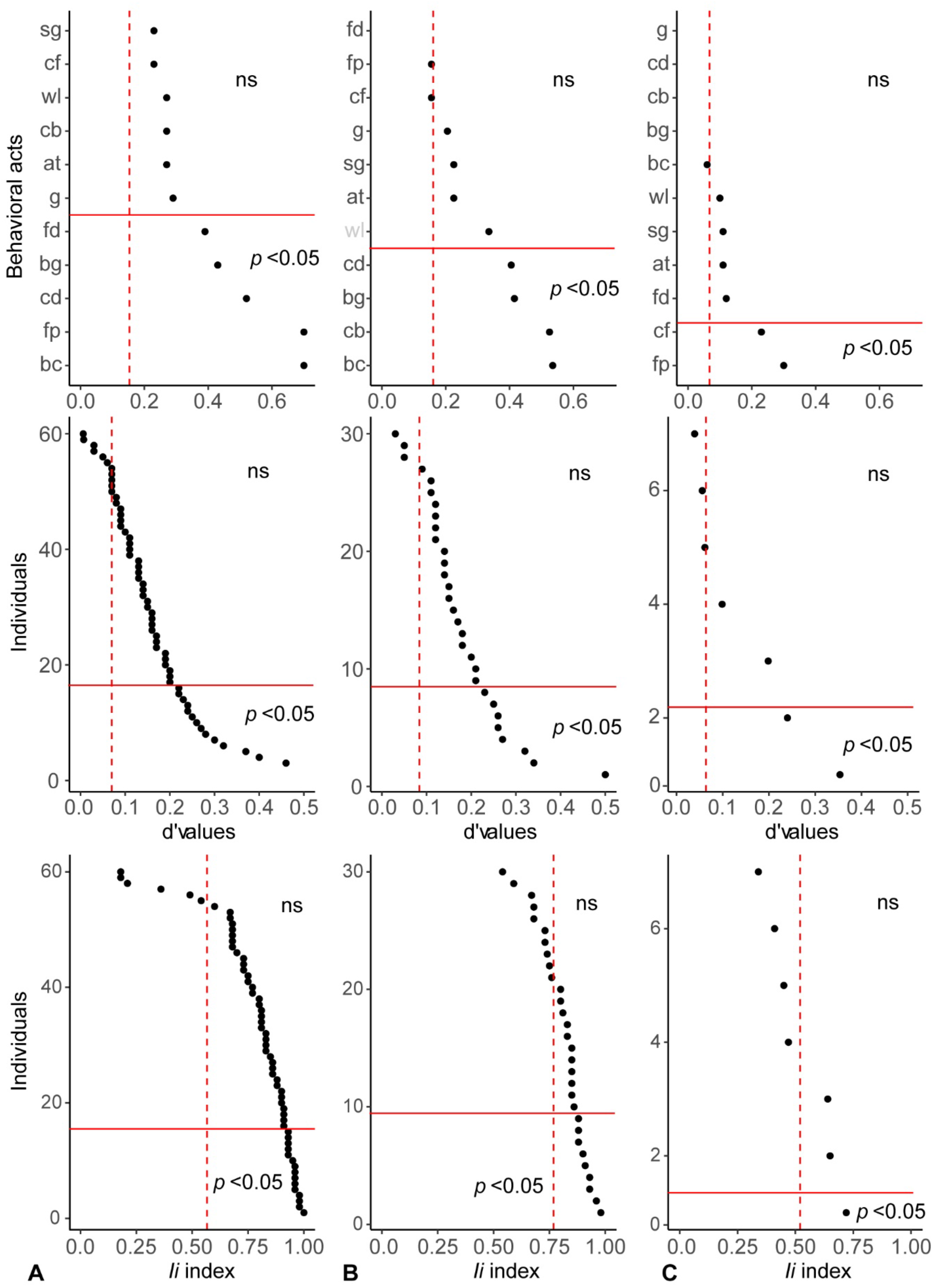
Data distribution (black dots) of the d’ values (d’_behavior_ and d’_indv_, i.e. metrics of specialization of behaviors and workers, respectively), and *Ii* (inactivity) index from WBNs (worker-behavior networks) of the colonies (A, B and C) of *Odontomachus chelifer* (Latreille, 1802). The y axis ticks in the d’_behavior_ mean each specific acronym representing the behavioral considered (presented in the Tab. 1). The red dashed line represents the mean of each null model considered. The red line divides significant data points (*p* <0.05).

### Centrality

The centrality measures allowed us to identify the influence of each behavior (i.e., bridges, inactivity hubs and spreaders) within BBNs. In colony A, brood care, self-grooming and walking were bridges. (Fig. 5A). In colony B, self-grooming and grooming were bridges (Fig. 5B). Differently, feeding was the only bridge behavior in colony C (Fig. 5C). The bridge behavior of brood care (colony A) had a betweenness centrality value significantly different from those obtained by the null model (Fig. 5A and Tab. 2). The frequencies of behavior switching significantly diverged from the self-interaction among the behaviors (*G*_(10)_ = 24; *P* < 0.001). Moreover, during active behavior switching, we observed that the frequency of behavioral switch of bridges was not uniform (Fig. 6). The bridges self-grooming, walking and feeding had significant interactions among themselves (*G*_(9)_ = 24, 23 and 45; *P* < 0.001), while the bridge brood-care had a significant interaction with carrying brood (*G*_(9)_ = 20; *P* < 0.001), and grooming had significant interaction with self-grooming (*G*_(9)_ = 22; *P* < 0.001) (Fig. 6).

**Fig. 5:**
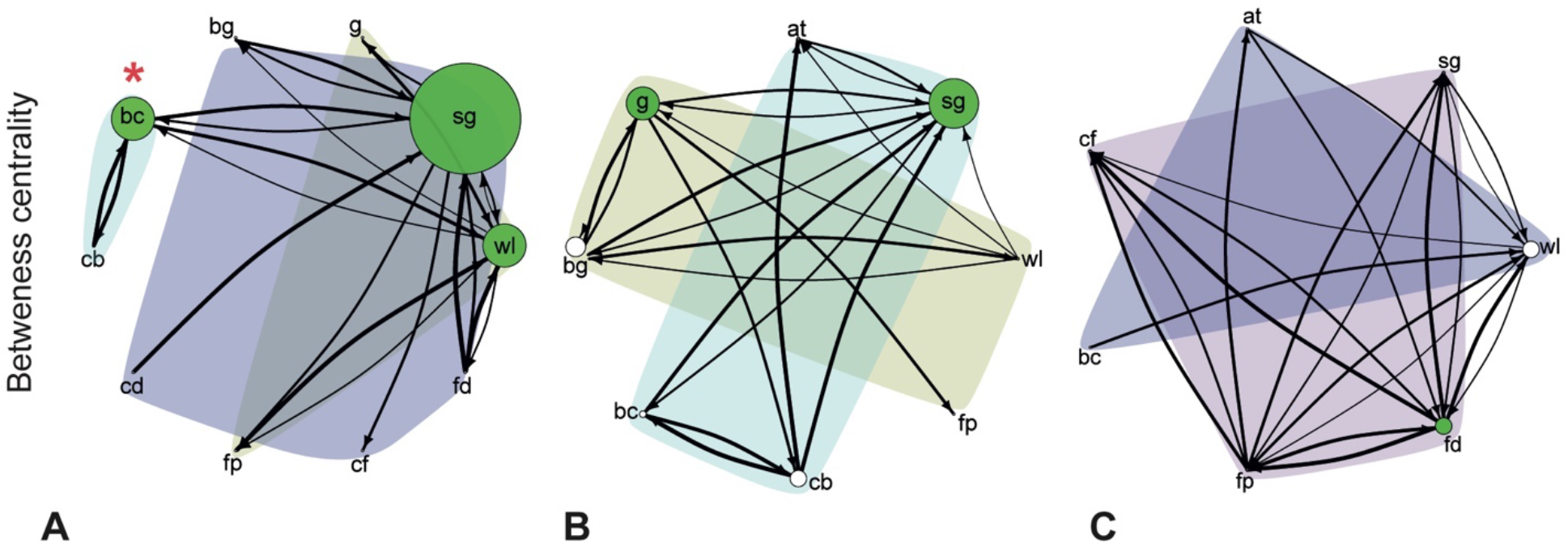
BBNs (behavior-behavior networks) graphs from the colonies (A, B and C) of *Odontomachus chelifer* (Latreille, 1802). The nodes represent behavioral acts and the links between them the interactions between behaviors from the workers. The width of each link indicates the frequency of interactions between behaviors and workers. The size of the nodes represents the betweenness centrality values of the nodes (the larger the node, the higher the betweenness centrality value obtained), nodes colored as green are bridges (i.e., nodes with betweenness centrality above the third quartile of the data, >75%, i.e. two or three behavioral acts with the highest scores). Bridges signaled with a red asterisk (*) are statistically significant (*p* <0.05) compared to the random networks. The different modules of tasks are identified in different colors.

**Fig. 6:**
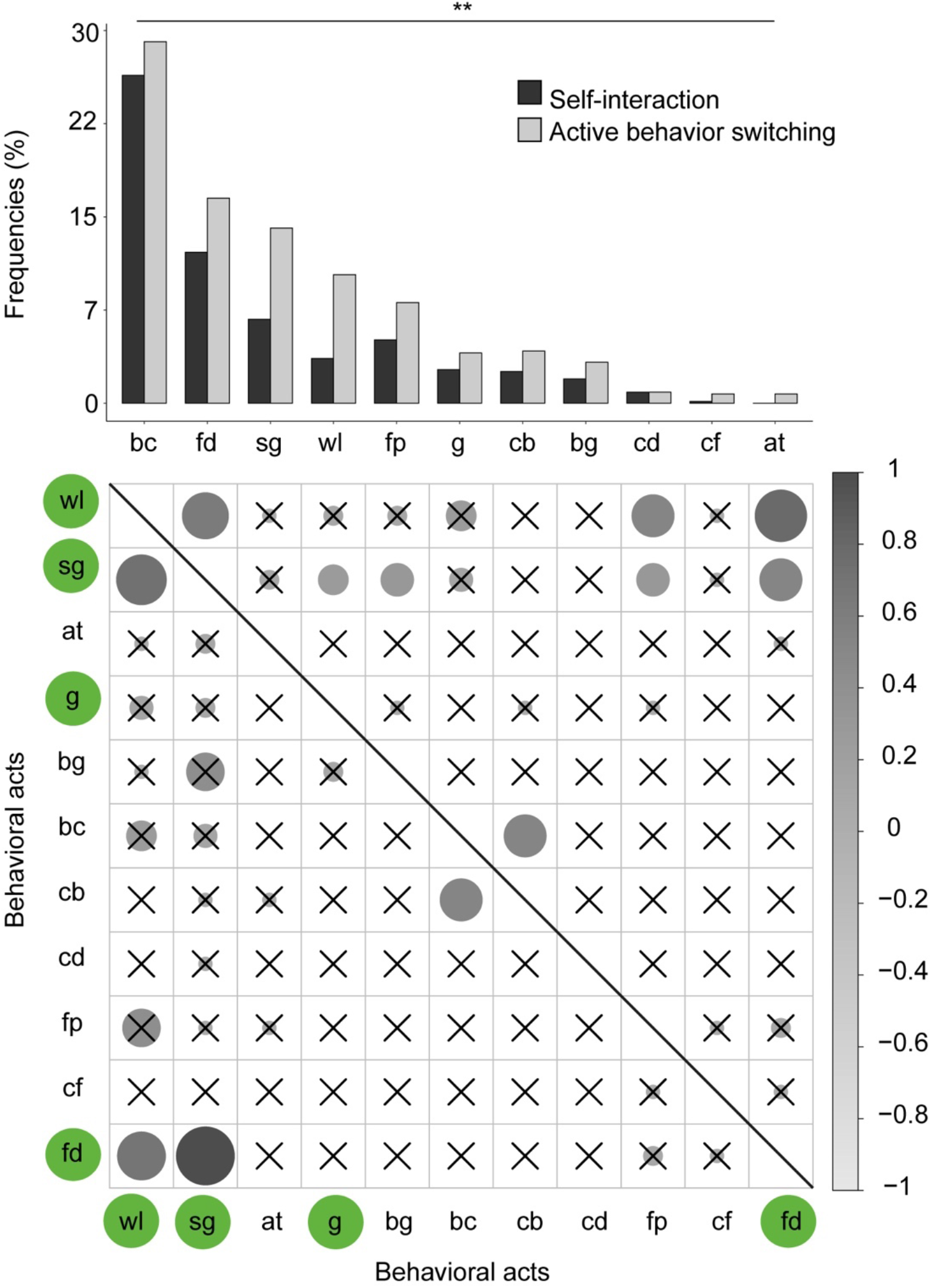
Comparison of the absolute frequency of the BBNs (behavior-behavior networks) for each active behavioral act, when they are in self-interaction or when occurs behavior switching with other active behaviors (statistical significance signaled by **, i.e., *p* < 0.01, upper plot). Adjacency matrix of the BBNs absolute frequencies during active behavior switching (lower plot). The higher the normalized value (−1 to 1), stronger the color and bigger the circle size. Bridges (i.e., nodes with betweenness centrality above the third quartile of the data, >75%) are colored as green. Circles within the matrix without an X had a frequency significantly higher (*p* <0.05) than the other behaviors of each column (post-hoc G test).

In colony A and B, self-grooming and grooming were inactivity hubs, with walking being an inactivity hub for colony A as well (Fig. 7A,B). In colony C, walking and feeding were the inactivity hubs (Fig. 7C). Inactivity hubs were not statistically different than the random networks, with the exception of grooming in colony B (Fig. 7 and Tab. 3). Feeding and self-grooming were inactivity spreaders in colony A (Fig. 7A). In colony B, grooming and brood care were inactive spreaders (Fig. 7B). In colony C, foraging/patrolling and feeding were inactivity spreaders (Fig. 7C). In colony A, self-grooming and feeding were significant inactivity spreaders compared to random networks. Differently, in colony B, brood-care was the only inactivity spreader significantly different from the null model (Fig. 7 and Tab. 4). Colony C did not have inactivity spreaders that were statistically higher than random networks (Fig. 7 and Tab. 4).

**Fig. 7:**
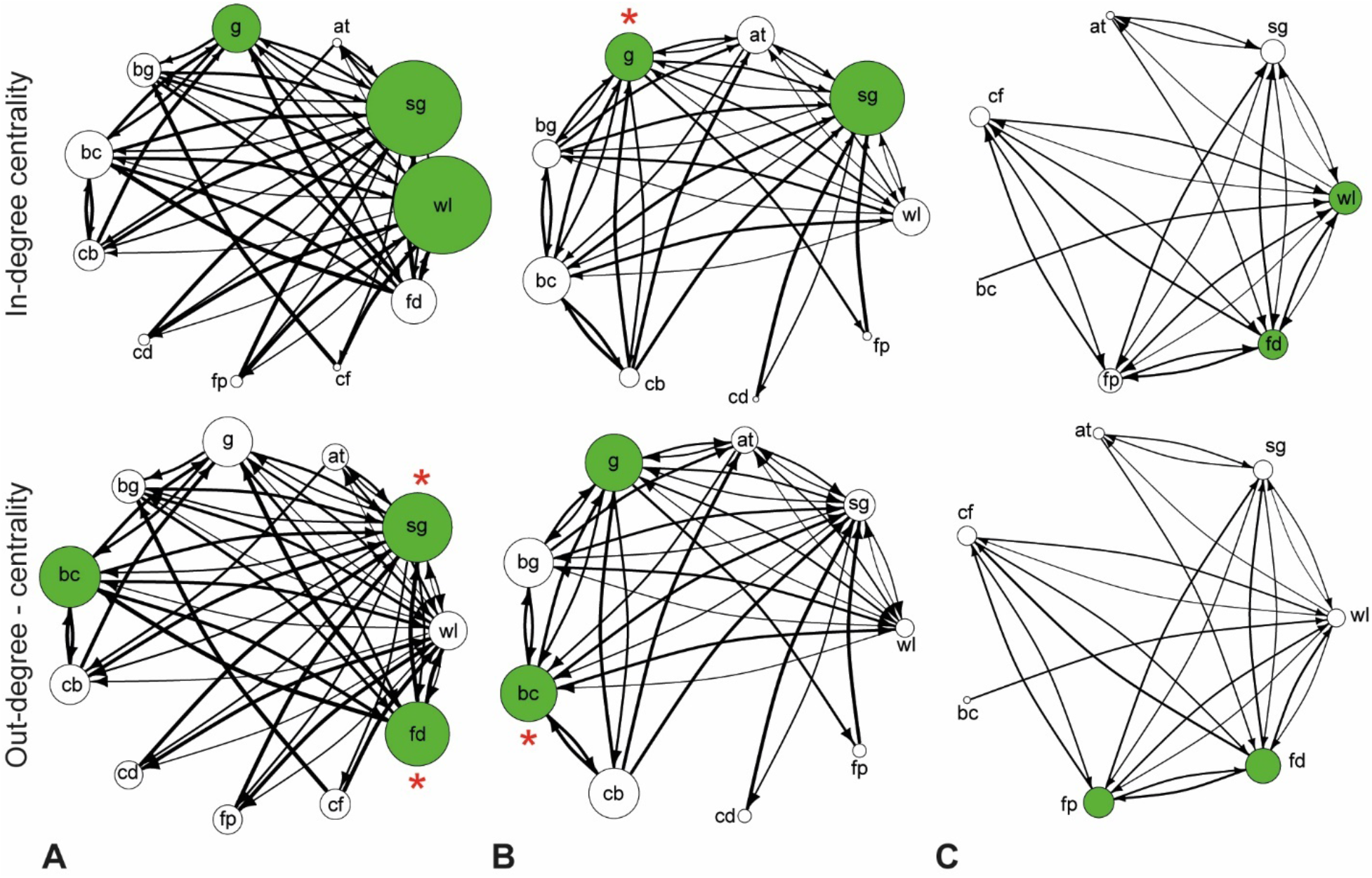
BBNs (behavior-behavior networks) of inactivity in-degree and out-degree graphs from the colonies (A, B and C) of *Odontomachus chelifer* (Latreille, 1802). The nodes represent behavioral acts and the links between them the inactivity between behaviors from the workers. The width of each link indicates the frequency of inactivity between behaviors and workers. The size of the nodes represents the in-degree and out-degree centrality values of the nodes (the larger the node, the higher the in-degree and out-degree centrality value obtained), nodes colored as green are inactivity hubs or spreaders (i.e., nodes with in-degree or out-degree centrality above the third quartile of the data, >75%, i.e. two or three behavioral acts with the highest scores, respectively). Inactivity hubs and spreaders signaled with a red asterisk (*) are statistically significant (*p* <0.05) compared to the random networks.

**Tab 3.**
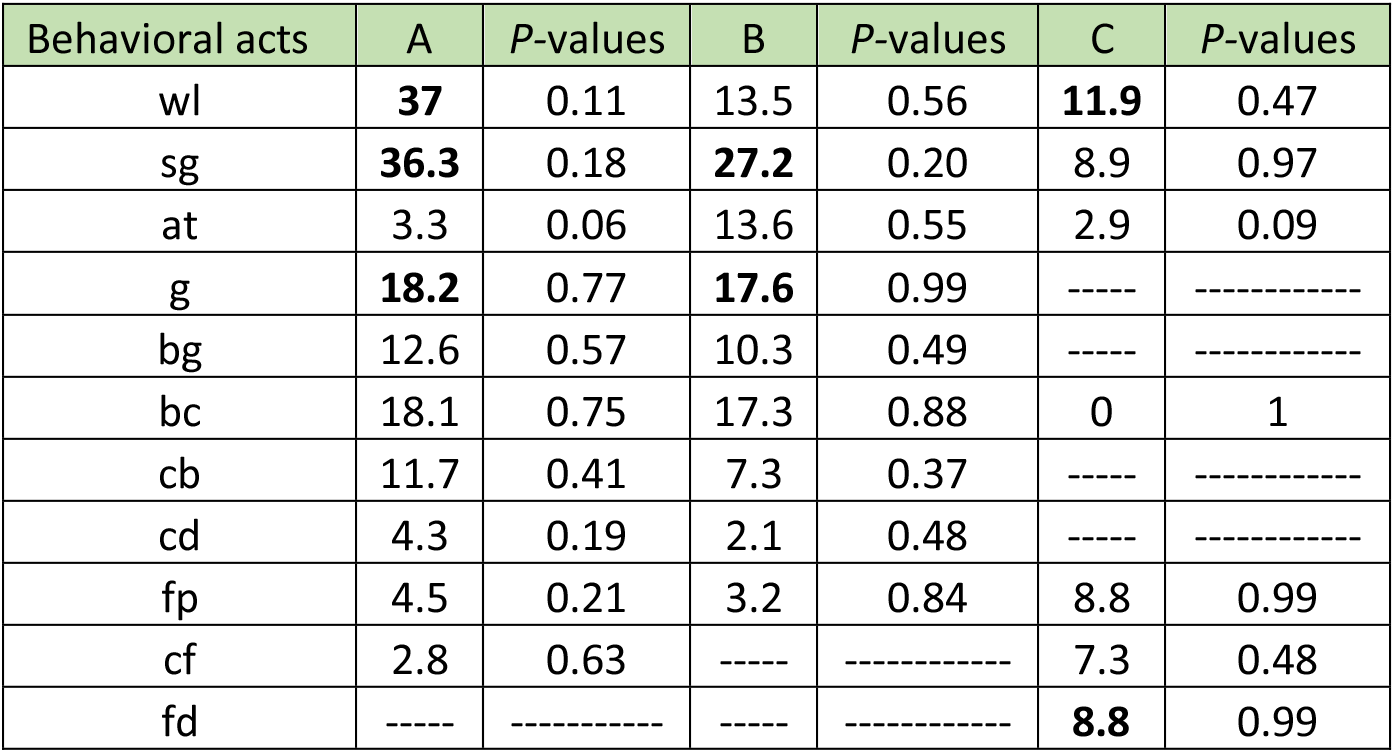
Degree centrality values (In-degree) from the BBNs (behavior-behavior networks) of all the colonies (A, B and C) of *Odontomachus chelifer* (Latreille, 1802). The behaviors (nodes) of all workers are considered. The behavioral acts are considered inactivity hubs if they have an in-degree centrality value above the third quartile of the data (> 75%; two or three behavioral acts with the highest scores, signaled in bold). The *P*-values of the Z-scores (expressed as significant in bold) of the data compared to the null model (weight reshuffling null model) are also exposed.

**Tab 4.**
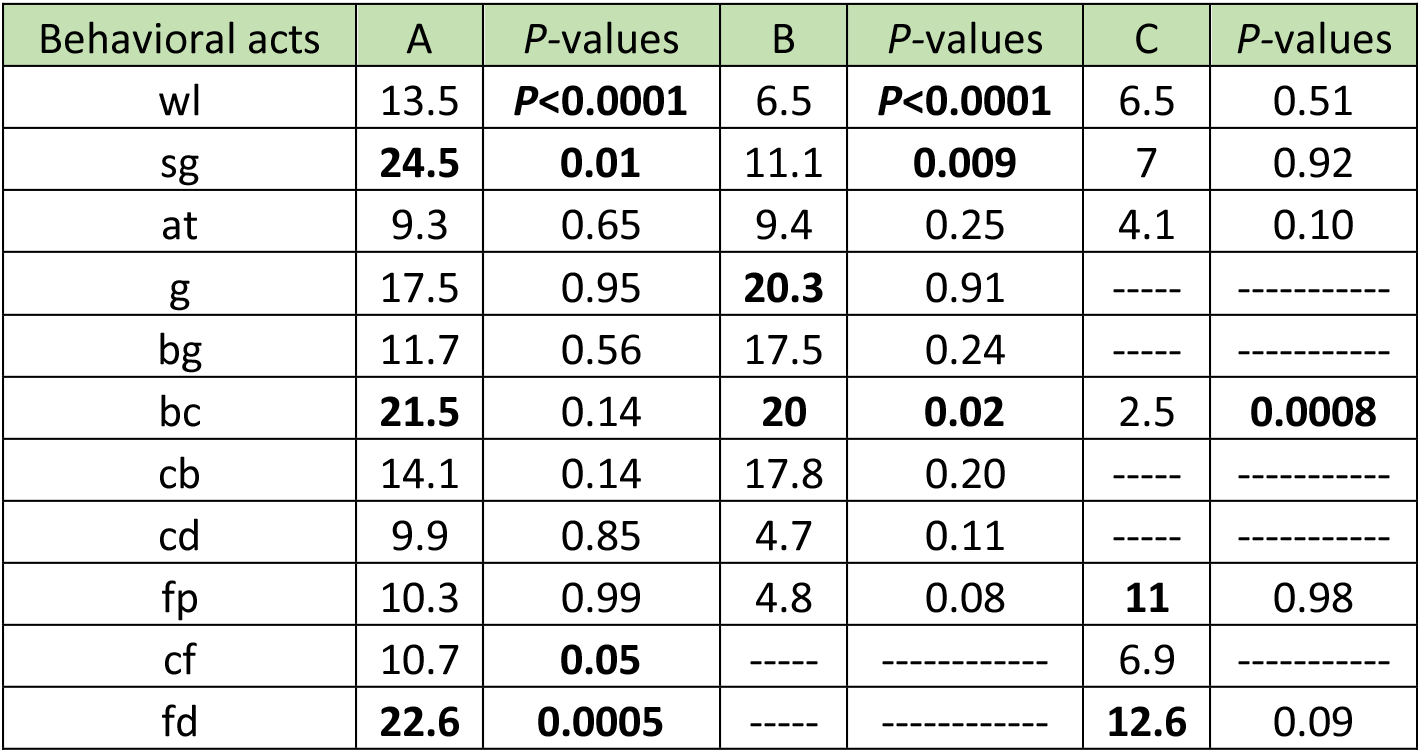
Degree centrality values (Out-degree) from the BBNs (behavior-behavior networks) of all the colonies (A, B and C) of *Odontomachus chelifer* (Latreille, 1802). The behaviors (nodes) of all workers are considered. The behavioral acts are considered inactivity spreaders if they have an out-degree centrality values above the third quartile of the data (>75%; two or three behavioral acts with the highest scores, signaled in bold). The *P*-values of the Z-scores (expressed as significant in bold) of the data compared to the null model (weight reshuffling null model) are also exposed.

### Modularity

Modularity values for the WBNs (*Q*_norm_) in all the colonies (Fig. 2) were significantly higher than the *Q*_norm_ obtained from random networks (Colony A, *Z* = 29.7, *P* < 0.0001; Colony B, *Z* = 12.54, *P* < 0.0001; Colony C, *Z* = 6.25, *P* < 0.0001). WBNs were organized into four modules in colony A (module 1: wl, sg, at and fp; module 2: g and bg, module 3: bc and cb, module 4: cd and fd) and C (module 1: wl, at and cf; module 2: sg, module 3: fd and bc, module 4: cd and fd) and six modules in colony B (module 1: wl cf; module 2: sg, module 3: at and g, module 4: bg, module 5: bc, module 6: cb, cd, cf and fp), the composition of the modules (number of individuals and behaviors interacting) differed between all colonies (Fig. 2). BBNs modularity values were not significantly higher compared to those obtained from random networks in colony A and C, but significant in colony B for all workers, (Fig. 8). BBNs in colony A were organized into three modules (Fig. 5, module 1: bc and cb; module 2: wl, fp and g; module 3: sg, fd, cf, cd, and bg). All BBNs were organized in two modules (Fig. 5) in colony B (module 1: g, bg, wl and fp; module 2: at, sg, bc and cb) and C (module 1: bc, at and wl; module 2: sg, cb, fp and fd). The composition of the modules (the interacting behaviors) within each colony differs with each worker composition.

**Fig. 8:**
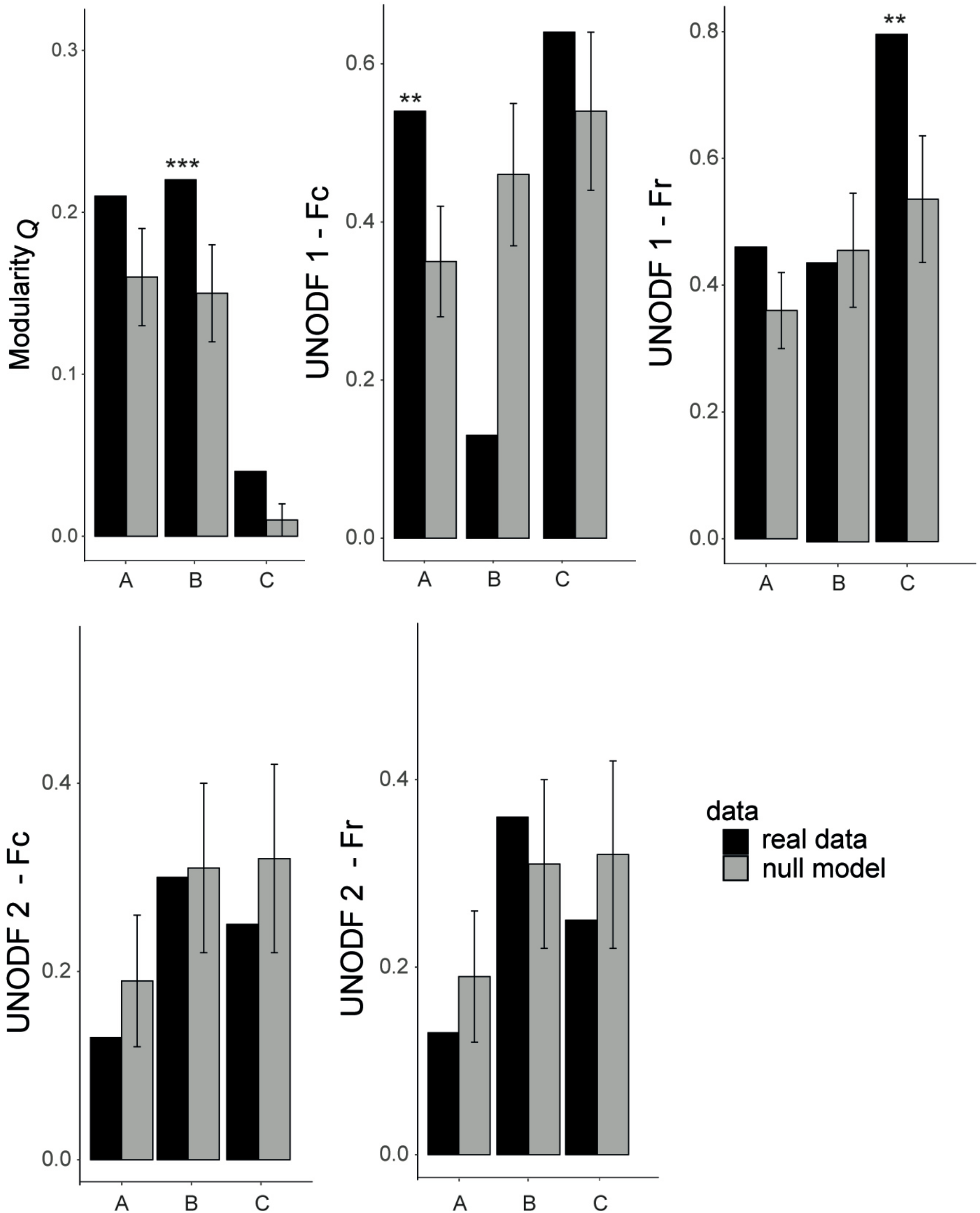
Network modularity (*Q*) and weighted nestedness (UNODF) for BBNs (behavior-behavior networks) from the colonies (A, B and C) of *Odontomachus chelifer* (Latreille, 1802). Black bars represent the original networks, while grey bars represent networks randomized and the respective standard deviation (SD). UNODF 1 is the metric calculated without a cut-off and UNODF 2 is the metric calculated with a cut-off of 10%. The significant statistical differences (Z-score) were signaled by *, ** and *** (*p* values less than 0.05, 0.01 and 0.001, respectively). The asterisks (*) above the bars mean significant differences between the original and the randomized networks or vice and versa.

### Nestedness

Nestedness (WNODF) value was significantly lower than those obtained from random networks (also known as an anti-nested pattern, but there are criticism about using this term, see Almeida-Neto & al. 2006) in colony A, while the values in colony B and C were not statistically significant (Fig. 3, Colony A, *Z* = −5, *P* < 0.0001; Colony B, *Z* = −1.64, *P* < 0.0001; Colony C, *Z* = −1.65, *P* < 0.0001). Nestedness (UNODF) values revealed that in general, the colonies did not have a nested structure in both the cut-off conditions considered from BBNs (Fig. 8). However, a nested structure was significantly present when considering all workers in colony A (UNODFc 1, *Z* = 2.59, *P* < 0.009) and C (UNODFr 1, *Z* = 2.39, *P* < 0.01).

## Discussion

This study shows that colonies of *Odontomachus chelifer* interact in structured networks that are consistent across colonies of different sizes. In short, inactivity is the most performed behavior in *O. chelifer*, whereas some behavioral acts such as walking, self-grooming (for all colonies) and brood care (for colony A and B) are more frequent. Dominance interactions are present but rarely observed. There is specialization within the colonies, with some individuals and behaviors more specialized than others. Centrality measures showed that some behaviors could have more impact in the network organization. Complex patterns such as modularity, nested and significantly not nested structures were observed in WBNs and BBNs. Our results are manifold and will be discussed with detail in turn.

Inactivity is by far the most recurrent behavior observed in all the colonies of *O. chelifer*. High inactivity frequency among ant workers (> 50% of the behaviors performed per colony) is a very widespread phenomena observed in several ant species, both in field and laboratory studies (Lindauer 1952, Herbers 1983, Herbers & Cunningham 1983, Cole 1986, Hölldobler & Wilson 1990, Schmid-Hempel 1990, Dornhaus & al. 2008, Dornhaus 2008, Dornhaus & al. 2009, Charbonneau & Dornhaus 2015). However, the role of inactivity is not usually considered to be part of task allocation strategies or colony organization (for exceptions, see Herbers 1981, Fresneau 1984, Cole 1986, Corbara & al. 1989, Retana & Cerdá 1990, Retana & Cerdá 1991, Charbonneau & Dornhaus 2015). It appears that, as already discussed in the literature, inactivity could have larger importance in the context of division of labor, given that some workers specialize in inactivity (Charbonneau & Dornhaus 2015). In *O. chelifer*, while some workers were clearly more inactive than others, the degree of inactivity performed by a worker (*Ii* index) was not directly associated with its degree of specialization (*d*’_indv_). This suggests that, while inactivity is the most performed behavior, its influence on behavior or task allocation is not quantified by its simple execution. The role of inactivity as a link between behavior switching will be discussed further through the analysis of inactivity hubs and spreaders.

Among the active behavioral acts, self-grooming and walking were on average the most performed behaviors by workers of all the colonies. Self-grooming is a self-maintenance (cleaning) task and a regulator of chemical signaling from hydrocarbons constituents (Soroker & al. 1998, Lahav & al. 1999). The hydrocarbon constituents of the postpharyngeal gland are sequestered by internal transport as well as from the body surface by self-grooming (Soroker & al. 1994, 1995a,b). Thus, the link between the postpharyngeal gland and body surface enables the ants continuously to refresh, and subsequently update their epicuticular hydrocarbons. The chemical signaling made by cuticular hydrocarbons and enhanced by self-grooming is probably a mechanism to maintain reproductive skew within the colonies. The fertility signal from chemical signaling communicates information that increase the individuals’ fitness (Keller & Nonacs 1993). Walking has innate importance in the performance of tasks, based upon the consequent movement of the ant to a designated task outside its position (Charbonneau & al. 2013), so it is not surprising that is one of the most performed behaviors by the workers.

Brood-care was significantly more frequent in larger colonies (A and B). Ants extensively present in the brood zone could have a higher reproductive status due to a close relationship with brood care and the production of eggs. This phenomenon is observed in *O. brunneus*, where social rank based on reproduction (proximity of allocated zone to the brood) is correlated to ovarian condition (Powell & Tschinkel 1999, Smith & al. 2012). While dominance interactions could have a role in the division of labor and emergence of hierarchy (Sasaki et al. 2016), i.e., reinforcing dominance/submission among workers, even suppressing worker brain dopamine (Shimoji et al. 2017), they were rare and not extensively observed in *O. chelifer*. This suggest that dominant behaviors are not a crucial process or the only one responsible for the division of labor in *O. chelifer*. Perhaps dominant interactions are important in a larger scale of time, or during few exceptions that demand a kind of control within the colony between workers (such as sudden changes in the food supply of the colony), or even in parallel with other kinds of behavioral control which do not demand physical contact (e.g., self-grooming). In addition, aggression (dominance interactions) and signaling by cuticular hydrocarbons enhanced by self-grooming are two possible modalities used to regulate reproduction in *O. chelifer*. They are both strong indicators of reproductive capacity (for aggression this was already discussed for vertebrates: Hrdy & Hrdy 1976; and insects: West-Eberhard 1967).

We observed that the division of labor of *O. chelifer* is organized through the significant presence of specialization within all studied colonies. The connection between specialization and subsequent better behavioral performance is scarce and contradictory in the literature (O’Donnell & Jeanne 1992, Dornhaus 2008, Russell & al. 2017, Santoro & al. 2019), but specialization must be important for other reasons within the colony organization. Specialization could be expected when you have interaction-based task allocation, such as observed for *O. brunneus* (Powell & Tschinkell 1999) and possibly to some degree (as already discussed) for *O. chelifer*. In the context of interaction-based task allocation, behavioral roles are naturally restricted to particular zones of the colony, therefore, allocation to a particular zone, through dominance interaction (or other processes, such as fertility signaling by self-grooming), ensure role specialization. We hypothesized that the behavioral acts which had more specialization across the colonies (brood care and foraging/patrolling) could be the most affected by this kind of dynamics (in the literature they are usually considered tasks). Risky and costly tasks for social insects, such as solitary foraging/patrolling (O’Donnell & Jeanne 1992, Perry & al. 2015) are made by specialists in *O. chelifer*, that inserted in the context of interaction-based task allocation have a lower social rank (i.e., a higher distance of the brood zone). Brood care specialization may also be the result of a reproductive hierarchy, differently than foraging/patrolling, brood care and carrying brood are probably performed by workers with higher social rank (within the brood zone). Moreover, the skewed distribution of *d*’_indv_ values clearly show that some workers are more specialized than others, where the worker force of the colony is composed by a mix of generalists and specialists, such pattern appears to be widespread in social insects (e.g., Jandt & al. 2009, Santoro & al. 2019). A partial division of labor, where generalists coexist with specialists, could be structurally important for the division of labor, for instance, such arrangement could generate more flexibility in the performance of tasks (Jandt & al. 2009). A mathematical model developed by D’orazio & Waite (2007) demonstrates that errors committed by generalist workers are few compared with the success of the group in general, thus the inefficiency and error-prone generalists may also be a fundamental feature of many of the social insect systems, as observed in wasps (Forsyth 1978, Jeanne 1986, Karsai & Wenzel 2000) and stingless bees (Hofstede & Sommeijer 2006). Moreover, there is a lot of possible explanation for the presence of specialization within colonies of eusocial insects, for instance, increased spatial efficiency for ants (Sendova & Franks 2005), or reduction of other switching costs (Chittka & al. 1997). It is also possible that specialization optimizes the performance of multistep tasks (Jeanne 1986). Together or separately, these processes may generate colony-level fitness advantages stemming from division of labor, even without observable improvement in individual efficiency (Dornhaus 2008).

While the interpretation of significance for the network-level metrics is quite intuitive, for instance, a significant positive nestedness Z-score indicates that the network is nested, and a significant negative one indicates a value less nested than randomized networks, this is not the case for node-level metrics, such as the betweenness and degree centrality (in and out-degree) measures from behavioral acts. A problem observed for centrality measures is that even a node with a negative Z-score, still has a higher centrality value than the other ones within the random network. Thus, we evaluate the significance of the comparison of the empirical centrality values (which were classified as bridges, inactivity hubs, and spreaders) to the random networks as simply a significant difference, rather than an attempt to interpret a positive or negative z-score. Information flow across tasks was intermediated by a set of different nodes (bridges), which varied accordingly to each colony. A pattern of occurrence of bridges was more visible than inactivity hubs and inactivity spreaders, which varied a lot across the colonies. Self-grooming had a prominent role (i.e., bridge) in the larger colonies (A and B), as well as feeding in the smaller colony (Colony C). Differently, Charbonneau & al. 2013 observed that walking had higher betweenness centrality compared to all other tasks for *Temnothorax rugatulus* being a significant bridge in our classification. This suggests that *O. chelifer* workers did not wander around the nest as much to switch behaviors. Self-grooming as a bridge between other behaviors gives evidence to the already discussed hypothesis of self-grooming as a reproductive regulator, with a crucial role for the maintenance of task allocation of the colony. Some association between behaviors were self-evident due to its inherent link (e.g. grooming and self-grooming) or logical sequence (after returning to the nest, the ant will walk within the nest prior to other behavior). Nevertheless, self-grooming had interesting associations, for instance, between walking and brood care. Self-grooming associated with walking could mean that the ant while moving within the nest display its reproductive status through self-grooming, same when manipulating the brood. Thus, self-grooming could be performed between behavioral acts to ensure reproductive status between workers, for instance, maintaining nurses (i.e., workers performing brood care and carrying brood) as nurses, and foragers as foragers. In colony C, however, feeding is a significant bridge. This gives another perspective of the phenomena of higher food intake in colony C, which occur between other behavior and that the bridge role is adjustable to the colony needs.

Modularity analysis offers the great advantage of providing a quantitative method to identify modules of preferentially interacting workers and behaviors and among behaviors. WBNs presented significant modularity, which means the existence of exclusive interactions between workers and behaviors. Modularity is thought to increase stability in ecological communities (May 1972; Krause & al. 2003; Teng & McCann 2004; however, see Pimm 1979). Similarly, the existence of modules in WBNs could generate stability as well: maybe if a worker is lost, another one from the same module could replace it minimizing the loss of a possible specialist. Differently, BBNs presented mixed results, with only colony B showing significant modularity. Among the modules of colony B, the association between them appears to be random, with the exception of brood care and carrying brood. They are usually performed intermittently between each other, frequently classified together as nursing behavior in the literature. The apparent randomness of the modular composition within BBNs could indicate the capacity of task flexibilization across the colonies of *O. chelifer*. However, our results showed that the behaviors are prone to interact more with some behaviors than others, thus modules if formed are apparently not representative of behavioral flexibilization (at least at a high degree).

WBNs were not nested, and colony A even was significantly less nested than randomized networks. The lack of a nested structure may indicate that the division of labor dynamics is vulnerable to worker loss, because a nested structure has been shown to protect against destruction of the network (Fortuna & Bascompte 2006, Burgos & al. 2007). In a nested network, the core group of generalists are more robust to loss of network individuals, they could easily substitute another worker. However, the relatively low degree of specialization present in the colonies (i.e., *H’*_2_ ≤ 0.40) might increase robustness (Pocock & al. 2012) and the workers within modules might fulfill similar interaction functions. Non-nested structures have been often observed in weighted ecological networks, and anti-nested patterns while rarer in nature (Staniczenko & al. 2013), were observed in interactions between fungi and plants (Bahram & al. 2014, Toju & al. 2014, 2015, Jacobsen & al. 2018), which could be explained by competitive exclusion (Toju & al. 2015). While competitive exclusion does not make sense in the context of division of labor, it could be analogous to the formation of strict modules without much connection with other nodes, since colony A had higher modularity compared to the other colonies. Some significant nested structures occurred in BBNs in colony A and C. Nestedness implies a hierarchy in the linking rules of the network system, so there is heterogeneity in the number of interactions among its elements. Furthermore, in ecological systems a nested structure is related to stability (e.g., Memmott & al. 2004, Burgos & al. 2007, Bastola & al. 2009). The presence of nestedness in behavioral allocation of ants could be viewed as a steady process, where individual adaptation could slightly change due to the necessities of the colony at some specific interval of time. Therefore, workers concentrate on some specific behaviors, but new ones could be performed trough colonial necessity. This pattern is foreseen by some mathematical models in social insects (e.g., Wilson 1985, Robinson 1987a, 1987b, 1992, Robinson & Page 1988, Calabi 1988, Detrain & Pasteels 1991, Page & Robinson 1991, Detrain & Pasteels 1992 Bonabeau & al. 1996).

The view of the ant colony as a complex system is not something new (Gordon 2010), but such view needs the use of a conceptual framework which simultaneously captures the complexity present within patterns of the colony and provides tools to analytically interpret the observed behavioral processes. The use of worker-behavior and behavior-behavior interactions constitute another layer of complexity for exploring the mechanisms that underlie individual variation within a network. It should be emphasized that this not exclude the assumption that interaction among workers (social interactions) are involved in task allocation, which could be analyzed together with worker-behavior and behavior-behavior interactions. The use of network concepts such as specialization, centrality, modularity and nestedness proved to be interesting for the description of the roles of the behaviors and workers in the organization of the division of labor. Furthermore, as previously suggested by Lewinsohn & al. (2006), simultaneously looking at several network patterns can substantially advance our understanding of the architecture of networks as well. A hindrance to the development of studies like this is the difficulty to account and quantify the real number of the tasks displayed by all the workers of the colony. Our study is still based on manual annotation of behaviors, while such approach is effective, it is time demanding and impractical for larger colonies or periods of time. Clever approaches such as the use of spatial fidelity for the determination of task performance (Mersch & al. 2013), while interesting to the study of several behavioral phenomena, it falls short to determine correct task performance or include behaviors that probably have an impact in colony organization (e.g., self-grooming, feeding). New improvements in approaches using automatic tracking by machine learning (which could encode and quantify behaviors; see Hong & al. 2015) could provide alternatives for this experimental gap. We expect even more holistic studies in the future, comparing species with different colony sizes to help the description of individual differences between workers.

## Supporting information

Appendix, as digital supplementary material to this article

## Acknowledgements

We acknowledge financial support to FMN and MEB from CNPq/MCT, which provided graduate fellowships (132204/2011-8 and 132004/2010-0, respectively). We are grateful to Simon Garnier for his constructive and helpful suggestions on an earlier version of our work.

## Appendix, as digital supplementary material to this article, at the journal’s web pages

Data S1a. Raw data for each individual (IN1-IN60) from the colony A of *Odontomachus chelifer* (Latreille, 1802).

Data S1b. Raw data for each individual (IN1-IN30) from the colony B of *Odontomachus chelifer* (Latreille, 1802).

Data S1c. Raw data for each individual (IN1-IN7) from the colony C of *Odontomachus chelifer* (Latreille, 1802).

Data S2. Frequencies of inactive and active behavior classes of *Odontomachus chelifer* (Latreille, 1802).

Data S3. WBN (worker-behavior network) from the colony A of *Odontomachus chelifer* (Latreille, 1802).

Data S4. WBN (worker-behavior network) considering inactivity behavior from the colony A of *Odontomachus chelifer* (Latreille, 1802).

Data S5. WBN (worker-behavior network) from the colony B of *Odontomachus chelifer* (Latreille, 1802).

Data S6. WBN (worker-behavior network) considering inactivity behavior from the colony B of *Odontomachus chelifer* (Latreille, 1802).

Data S7. WBN (worker-behavior network) from the colony C of *Odontomachus chelifer* (Latreille, 1802).

Data S8. WBN (worker-behavior network) considering inactivity behavior from the colony C of *Odontomachus chelifer* (Latreille, 1802).

Data S9. BBNs (behavior-behavior networks) from each individual of the colonies (A, B and C) of *Odontomachus chelifer* (Latreille, 1802).

Data S10. BBNs (behavior-behavior networks) of inactivity from each individual of the colonies (A, B and C) of *Odontomachus chelifer* (Latreille, 1802).

Data S11. BBN (behavior-behavior network) from the colony A of *Odontomachus chelifer* (Latreille, 1802).

Data S12. BBN (behavior-behavior network) considering inactivity behavior from the colony A of *Odontomachus chelifer* (Latreille, 1802).

Data S13. BBN (behavior-behavior network) from the colony B of *Odontomachus chelifer* (Latreille, 1802).

Data S14. BBN (behavior-behavior network) considering inactivity behavior from the colony B of *Odontomachus chelifer* (Latreille, 1802).

Data S15. BBN (behavior-behavior network) from the colony C of *Odontomachus chelifer* (Latreille, 1802).

Data S16. BBN (behavior-behavior network) considering inactivity behavior from the colony C of *Odontomachus chelifer* (Latreille, 1802).

Data S17. *H2* index (specialization index) from the colonies (A, B and C) of *Odontomachus chelifer* (Latreille, 1802) compared to the values obtained from the null models considered.

Data S18. DOLi (specialization index for individuals) from the colonies (A, B and C) of *Odontomachus chelifer* (Latreille, 1802) compared to the values obtained from the null models considered.

Data S19. DOLt (specialization index for tasks) from the colonies (A, B and C) of *Odontomachus chelifer* (Latreille, 1802) compared to the values obtained from the null models considered.

Data S20. Matrix of *P*-values (post-hoc G-test) from the absolute frequency of the BBNs (behavior-behavior networks) for each active behavioral act, when they are in self-interaction or when occurs behavior switching with other active behaviors.

Data S21. Modularity (*Q*) values from the colonies (A, B and C) of *Odontomachus chelifer* (Latreille, 1802) compared to the values obtained from the null models considered.

Data S22. Modularity (*Q*) values and standard deviation (SD) from BBNs (behavior-behavior networks) of the colonies (A, B and C) of *Odontomachus chelifer* (Latreille, 1802) and the null model considered.

Data S23. WNODF (weighted nestedness) from the colonies (A, B and C) of *Odontomachus chelifer* (Latreille, 1802) compared to the values obtained from the null models considered.

Data S24. Nestedness among columns considering all data (UNODFc) and standard deviation (SD) of the BBNs (behavior-behavior networks) from the colonies of *Odontomachus chelifer* (Latreille, 1802) and the null model considered.

Data S25. Nestedness among rows considering all data (UNODFr) and standard deviation (SD) from BBNs (behavior-behavior networks) of the colonies of *Odontomachus chelifer* (Latreille, 1802) and the null model considered.

Data S26. Nestedness among columns considering a cut-off of 10% of the data (UNODFc) and standard deviation (SD) from BBNs (behavior-behavior networks) of the colonies of *Odontomachus chelifer* (Latreille, 1802) and the null model considered.

Data S27. Nestedness among rows considering a cut-off of 10% of the data (UNODFr) and standard deviation (SD) from BBNs (behavior-behavior networks) of the colonies of *Odontomachus chelifer* (Latreille, 1802) and the null model considered.

Script S1. R script for the analysis of the behavioral repertoires from the colonies of *Odontomachus chelifer* (Latreille, 1802).

Script S2. R script for the analysis of the WBNs (worker-behavior networks) from the colonies of *Odontomachus chelifer* (Latreille, 1802).

Script S3. R script to read each individual BBNs (behavior-behavior networks) from the colonies of *Odontomachus chelifer* (Latreille, 1802).

Script S4. R script for the creation of the BBN (behavior-behavior network) and UNODF analysis (weighted nestedness) from the colony A of *Odontomachus chelifer* (Latreille, 1802).

Script S5. R script for the creation of BBN (behavior-behavior network) and UNODF analysis (weighted nestedness) from the colony B of *Odontomachus chelifer* (Latreille, 1802).

Script S6. R script for the creation of BBN (behavior-behavior network) and UNODF analysis (weighted nestedness) from the colony C of *Odontomachus chelifer* (Latreille, 1802).

Script S7. R script for the creation of BBNs (behavior-behavior networks) null models by the TNET package version 03.0.16.

Script S8. R script for the creation of BBNs (behavior-behavior networks) null models of inactivity by the TNET package version 03.0.16.

Script S9. R script of the comparison of the absolute frequency of the BBNs (behavior-behavior networks) for each active behavioral act, when they are self-interacting or when occurs behavior switching with other active behaviors.

Script S10. R script of the BBNs (behavior-behavior networks) null models for betweenness centrality from the colonies of *Odontomachus chelifer* (Latreille, 1802).

Script S11. R script of the BBNs (behavior-behavior networks) null models for degree centrality from the colony A of *Odontomachus chelifer* (Latreille, 1802).

Script S12. R script of the BBNs (behavior-behavior networks) null models for degree centrality from the colony B of *Odontomachus chelifer* (Latreille, 1802).

Script S13. R script of the BBNs (behavior-behavior networks) null models for degree centrality from the colony C of *Odontomachus chelifer* (Latreille, 1802).

Script S14. R script of the BBNs (behavior-behavior networks), modularity *(Q)* values and weighted nestedness (UNODF) from the colonies of *Odontomachus chelifer* (Latreille, 1802).

Script S15. R script of the BBNs (behavior-behavior networks) null models of the modularity (*Q*) values from the colonies of *Odontomachus chelifer* (Latreille, 1802).

