## Appendix, as digital supplementary material to this article for "Worker-behavior and behavior-behavior interaction networks in the trap-jaw ant *Odontomachus chelifer* (Latreille, 1802) (Hymenoptera: Formicidae)": Script S1.R

```
library(magritrr)
library(dplyr)
library(ggplot2)
library(RVAideMemoire)
library(ggpubr)
library(reshape2)

A_im<-c(2996, 664)
B_im<-c(1541, 290)
C_im<-c(286, 142) 

behavfreq_im$A_im = behavfreq_im$A_im / sum(behavfreq_im$A_im)*100
behavfreq_im$B_im = behavfreq_im$B_im / sum(behavfreq_im$B_im)*100
behavfreq_im$C_im = behavfreq_im$C_im / sum(behavfreq_im$C_im)*100

behaviors_im <-c("im", "other")
behavfreq_im <- behavfreq_im %>% `row.names<-`(behaviors_im)
behavfreq_im <- melt(behavfreq_im)
behavfreq_im<-as.data.frame(cbind(A_im,B_im,C_im))
##

behavfreq_im<-read.table("Data S2.txt", header = TRUE, row.names=NULL)

im_stack<- ggplot(behavfreq_im, aes(x=factor(colonies), y= value)) + geom_bar(aes(fill= behav), stat="identity", position="stack") + theme_classic() + labs(x = "Colonies", y = "Frequencies (%)")+
  scale_fill_grey() + geom_text(aes(y=value, label=value), vjust=1.6, color="white")


behaviors_in<-c("im", 'wl', 'sg', 'at', 'bg', 'fd', 'g', 'bc', 'cb', 'cd', 'fp', 'cf')
A<-c(2996, 122,156,10,11,144,26,154,13,11,11,6)
B<-c(1541, 66,80,11,36,NA,17,56,22,1,1,NA)
C<-c(286, 62,31,3,NA,21,NA,1,NA,NA,21,3) 

behavfreq_with_in<-as.data.frame(cbind(A,B,C))
behavfreq_with_in <- behavfreq_with_in %>% `row.names<-`(behaviors_in)

#stats comparison with ianctivity versus other behaviors
G.multcomp(behavfreq_with_in$A, p.method = "BH")
G.multcomp(na.omit(behavfreq_with_in$B), p.method = "BH")
G.multcomp(na.omit(behavfreq_with_in$C), p.method = "BH")

##########################

behaviors<-c('wl', 'sg', 'at', 'bg', 'fd', 'g', 'bc', 'cb', 'cd', 'fp', 'cf')
A<-c(122,156,10,11,144,26,154,13,11,11,6)
B<-c(66,80,11,36,NA,17,56,22,1,1,NA)
C<-c(62,31,3,NA,21,NA,1,NA,NA,21,3) 

behavfreq<-as.data.frame(cbind(A,B,C))

behavfreq$A_pct = behavfreq$A / sum(behavfreq$A)
behavfreq$B_pct = behavfreq$B / sum(behavfreq$B)
behavfreq$C_pct = behavfreq$C / sum(behavfreq$C)

behavfreq = mutate(behavfreq, A_pct = A/sum(A, na.rm=TRUE)*100, 
                   B_pct = B/sum(B, na.rm = TRUE)*100,
                   C_pct = C/sum(C, na.rm = TRUE)*100)

behavfreq = behavfreq %>% rowwise() %>% mutate(Avg = (mean(c(A_pct, B_pct, C_pct), na.rm = T)))  

behavfreq <- behavfreq %>% `row.names<-`(behaviors)

Font_My_Theme = theme(
  axis.title.x = element_text(size = 10),
  axis.title.y = element_text(size = 10),
  axis.text.x = element_text(size = 10),
  axis.text.y = element_text(size = 10))

#all<- ggplot(data=behavfreq , aes(x=reorder(behaviors, -Avg), y=Avg)) + geom_bar(stat="identity", fill="black") +
#theme_classic() + labs(x = "Behavioral acts", y = "%") +  ylim(0, 45) + geom_hline(yintercept=10.22, linetype="dashed", color = "red") + Font_My_Theme +
#geom_text(aes(label=Avg), vjust=-0.3, size=3.5)

ca <- ggplot(data=behavfreq , aes(x=reorder(behaviors, -A_pct), y=A_pct)) + geom_bar(stat="identity", fill="black")+
theme_classic() + labs(x = "Behavioral acts", y = "Frequencies (%)") +  ylim(0, 45) + geom_hline(yintercept=9, linetype="dashed", color = "red") + Font_My_Theme 

cb <- ggplot(data=behavfreq , aes(x=reorder(behaviors, -B_pct), y=B_pct)) + geom_bar(stat="identity", fill="black")+
theme_classic() + labs(x = "Behavioral acts", y = "Frequencies (%)") +  ylim(0, 45) + geom_hline(yintercept=11, linetype="dashed", color = "red") + Font_My_Theme  

cc <- ggplot(data=behavfreq , aes(x=reorder(behaviors, -C_pct), y=C_pct)) + geom_bar(stat="identity", fill="black")+
theme_classic() + labs(x = "Behavioral acts", y = "Frequencies (%)") +  ylim(0, 45) + geom_hline(yintercept=11, linetype="dashed", color = "red") + Font_My_Theme 

#stats behavior frequency
G.theo.multcomp(behavfreq$A, p.method = "BH")
G.theo.multcomp(na.omit(behavfreq$B), p.method = "BH")
G.theo.multcomp(na.omit(behavfreq$C), p.method = "BH")
###################

#Figure 1 unedited 
ggarrange(im_stack, ca, cb, cc, labels = c("all", "A", "B", "C"), legend = "bottom", font.label = list(size = 12))
```
