## Appendix, as digital supplementary material to this article for "Worker-behavior and behavior-behavior interaction networks in the trap-jaw ant *Odontomachus chelifer* (Latreille, 1802) (Hymenoptera: Formicidae)": Script S2.R

```
#load packages
library(bipartite)
library(igraph)
library(ggplot2)
library(dplyr)
library(ggpubr)

#font theme for the plots 
Font_My_Theme = theme(axis.title.x = element_text(size = 10), axis.title.y = element_text(size = 10),
  axis.text.x = element_text(size = 10), axis.text.y = element_text(size = 10))

#WBNs - read data from the colonies 
WBNs_a<-read.table("Data S3.txt", header=TRUE)
WBNs_b<-read.table("Data S5.txt", header=TRUE)
WBNs_c<-read.table("Data S7.txt", header=TRUE)

#plot IBN network - Figure 1 elements - colored version was made within an image editor
plotweb(WBNs_a, method="normal")
plotweb(WBNs_b, method="normal")
plotweb(WBNs_c, method="normal")

#Z-score 
net.score = function(obsval, nullval) {
  (obsval - mean(nullval))/sd(nullval)
} 

#P values of z-score
add.pvalues = function(zscore) {
  2*pnorm(-abs(zscore))
}

#Generate null models using Patefield's algorithm for colony a, b and c.
netnulls.r2d_WBNs_a <- nullmodel(WBNs_a, N=1000, method="r2dtable")
netnulls.r2d_WBNs_b <- nullmodel(WBNs_b, N=1000, method="r2dtable")
netnulls.r2d_WBNs_c<- nullmodel(WBNs_c, N=1000, method="r2dtable")

######################
#WBNs H2 values
H2fun(WBNs_a) #0.40
H2fun(WBNs_b) #0.36
H2fun(WBNs_c) #0.18

#generating H2 values for the random networks 
h2.nulls_a <- lapply(netnulls.r2d_WBNs_a, H2fun) 
h2.nulls_b <- lapply(netnulls.r2d_WBNs_b, H2fun) 
h2.nulls_c <- lapply(netnulls.r2d_WBNs_c, H2fun) 

H2.nulls_a<-unlist(lapply(h2.nulls_a, "[[", 1))
H2.nulls_b<-unlist(lapply(h2.nulls_b, "[[", 1)) 
H2.nulls_c<-unlist(lapply(h2.nulls_c, "[[", 1)) 

net.score(0.40, H2.nulls_a); add.pvalues(29.38277) # z-score 29.38277  p = 9.117781e-190
net.score(0.36, H2.nulls_b); add.pvalues(15.55624) # z-score 15.83643  p = 1.744477e-56
net.score(0.18, H2.nulls_c); add.pvalues(3.233083) # z-score 10.06042 p = 0.00122462

######
#plot of the empirical H2 values versus  the distribution of the null models
H2<-read.table("Data S17.txt", header = TRUE)
H2$Data <- factor(H2$Data, levels = c("H2", "null_model"))

H2plot<- ggplot(H2, aes(x=colonies, y=H2_value, fill=Data)) + 
  geom_bar(stat="identity", color="black", 
           position=position_dodge()) +
  geom_errorbar(aes(ymin=H2_value-sd_data, ymax=H2_value+sd_data), width=.2,
                position=position_dodge(.9)) + theme_classic()  + scale_fill_grey()

###################################

#IBN d' values - Calculates standardised specialisation index d' (d prime) for tasks
#colony A
dfun(WBNs_a)
dfun(WBNs_b)
dfun(WBNs_c)

############################ d tasks
behaviors<-c('wl', 'sg', 'at', 'bg', 'g', 'bc', 'cb', 'cd', 'cf', 'fd', 'fp')
#dtask
#           wl    sg    at    bg    g     bc     cb   cd    cf     fd   fp
dbehavior_a<-c(0.27, 0.23, 0.27, 0.43, 0.29, 0.70, 0.27, 0.52, 0.23, 0.39, 0.70)
#           wl    sg     at    bg    g     bc    cb    cd   cf   fd     fp       
dbehavior_b<-c(0.34, 0.23, 0.23, 0.42, 0.21, 0.54, 0.53, 0.41, 0.16, NA, 0.16)
#           wl    sg    at  bg   g    bc   cb  cd   cf     fd    fp     
dbehavior_c<-c(0.10, 0.11, 0.11, NA, NA, 0.06, NA, NA, 0.23, 0.12, 0.30)
#
dbehavior_all<-as.data.frame(cbind(dbehavior_a,dbehavior_b, dbehavior_c))
dbehavior_all <- dbehavior_all %>% `row.names<-`(behaviors)

dbehavior.nulls_a <- lapply(netnulls.r2d_WBNs_a, dfun) #
dbehavior.nulls_b <- lapply(netnulls.r2d_WBNs_b, dfun) #
dbehavior.nulls_c <- lapply(netnulls.r2d_WBNs_c, dfun) #

dbehavior_prime.nulls_a<-unlist(lapply(dbehavior.nulls_a, "[[", 1))
dbehavior_prime.nulls_b<-unlist(lapply(dbehavior.nulls_b, "[[", 1))
dbehavior_prime.nulls_c<-unlist(lapply(dbehavior.nulls_c, "[[", 1))

#z-score dbehavior colony A - significant ones bg, bc, cd, fd, fp  
#wl
net.score(0.27, dbehavior_prime.nulls_a); add.pvalues(1.16) # z-score 1.16 p = 0.24
#sg
net.score(0.23, dbehavior_prime.nulls_a); add.pvalues(0.67) #z-score 2.16 p = 0.50
#at
net.score(0.27, dbehavior_prime.nulls_a); add.pvalues(1.16) # z-score 2.72 p = 0.24
#bg
net.score(0.43, dbehavior_prime.nulls_a); add.pvalues(3.10) # z-score 3.10 p = 0.001935206
#g
net.score(0.29, dbehavior_prime.nulls_a); add.pvalues(1.40) # z-score 1.40 p = 0.16
#bc
net.score(0.70, dbehavior_prime.nulls_a); add.pvalues(6.38) # z-score 6.38 p = 1.77088e-10
#cb
net.score(0.27, dbehavior_prime.nulls_a); add.pvalues(1.16) # z-score 1.16 p = 0.2460488
#cd
net.score(0.52, dbehavior_prime.nulls_a); add.pvalues(4.20) # z-score 4.20 p =  2.66915e-05
#cf
net.score(0.23, dbehavior_prime.nulls_a); add.pvalues(0.67) # z-score 2.16 p = 0.50
#fd
net.score(0.39, dbehavior_prime.nulls_a); add.pvalues(2.62) # z-score 2.62 p = 0.008
#fp
net.score(0.70, dbehavior_prime.nulls_a); add.pvalues(6.38) # z-score 6.38 p = 1.77088e-10

#########################
#              wl    sg     at    bg    g     bc    cb    cd    cf   fd   fp       
dbehavior_b<-c(0.34, 0.23, 0.23, 0.42, 0.21, 0.54, 0.53, 0.41, 0.16, NA, 0.16)
#z-score dtask col b # significative from 0.23-  significant ones bg, cd, bc, cb
#wl
net.score(0.34, dbehavior_prime.nulls_b); add.pvalues(1.58) # z-score 1.58 p = 0.11
#sg
net.score(0.23, dbehavior_prime.nulls_b); add.pvalues(0.54) # z-score 2.11 p = 0.58
#at
net.score(0.23, dbehavior_prime.nulls_b); add.pvalues(0.54) # z-score 2.11 p = 0.58
#bg
net.score(0.42, dbehavior_prime.nulls_b); add.pvalues(2.34) # z-score 2.34 p = 0.01
#g
net.score(0.21, dbehavior_prime.nulls_b); add.pvalues(0.36) # z-score 0.36 p = 0.71
#bc
net.score(0.54, dbehavior_prime.nulls_b); add.pvalues(3.47) # z-score 3.47 p = 0.0005204584
#cb
net.score(0.53, dbehavior_prime.nulls_b); add.pvalues(3.37) # z-score 3.37 p = 0.0007516818
#cd
net.score(0.41, dbehavior_prime.nulls_b); add.pvalues(2.24) # z-score 2.24 p = 0.02
#cf
net.score(0.16, dbehavior_prime.nulls_b); add.pvalues(-0.11) # z-score -0.11 p = 0.91
#fp
net.score(0.16, dbehavior_prime.nulls_b); add.pvalues(-0.11) # z-score -0.11 p = 0.91

#########################
#z-score dbehavior col c  significant above 0.23
#                wl    sg    at  bg   g   bc   cb  cd   cf    fd    fp     
dbehavior_c<-c(0.10, 0.11, 0.11, NA, NA, 0.06, NA, NA, 0.23, 0.12, 0.30)
#z-score dtask col b # significant ones fp and cf

#wl
net.score(0.10, dbehavior_prime.nulls_c); add.pvalues(0.54) # z-score 0.54 p = 0.58
#sg
net.score(0.11, dbehavior_prime.nulls_c); add.pvalues(0.70) #z-score 0.68 p = 0.48
#at
net.score(0.11, dbehavior_prime.nulls_c); add.pvalues(0.68) #z-score 0.68 p = 0.49
#bc
net.score(0.06, dbehavior_prime.nulls_c); add.pvalues(-0.09) # z-score -0.09 p = 0.92
#cf
net.score(0.23, dbehavior_prime.nulls_c); add.pvalues(2.57) # z-score 2.57 p = 0.01
#fp
net.score(0.12, dbehavior_prime.nulls_c); add.pvalues(0.84) # z-score 0.84 p = 0.40
#fp
net.score(0.30, dbehavior_prime.nulls_c); add.pvalues(3.67) # z-score 3.67 p = 0.0002425505

#############################################
ca_dbehav<-ggplot(data=dbehavior_all , aes(x=reorder(behaviors, -dbehavior_a), y=dbehavior_a)) + geom_point(stat="identity", fill="black")+
  theme_classic() + labs(x = "Behavioral acts", y = "d'behavior") + ylim(0, 0.7) + geom_hline(yintercept=0.17, linetype="dashed", color = "red") + Font_My_Theme + coord_flip()

cb_dbehav<-ggplot(data=dbehavior_all , aes(x=reorder(behaviors, -dbehavior_b), y=dbehavior_b)) + geom_point(stat="identity", fill="black")+
  theme_classic() + labs(x = "Behavioral acts", y = "d'behavior") + ylim(0, 0.7) + geom_hline(yintercept=0.17, linetype="dashed", color = "red") + Font_My_Theme + coord_flip()

cc_dbehav<-ggplot(data=dbehavior_all , aes(x=reorder(behaviors, -dbehavior_c), y=dbehavior_c)) + geom_point(stat="identity", fill="black")+
  theme_classic() + labs(x = "Behavioral acts", y = "d' values") + ylim(0, 0.7) + geom_hline(yintercept=0.06, linetype="dashed", color = "red") + Font_My_Theme + coord_flip()

#####################################################################

#IBN d' values - Calculates standardised specialisation index d' (d prime) for workers - d'worker
dfun(t(WBNs_a))
dfun(t(WBNs_b))
dfun(t(WBNs_c))

dindv_a<-c(0.07, 0.46, 0.006, 0.09, 0.13, 0.09, 0.05, 0.11, 0.11, 0.22, 0.20, 0.08, 0.16, 0.14, 0.08, 0.07, 0.24, 0.20, 0.20, 0.25, 0.24, 0.10, 0.007, 0.16, 0.16, 0.11, 0.17, 0.27, 0.40, 0.13, 0.13, 0.14, 0.37, 0.26, 0.22, 0.51, 0.51, 0.06, 0.16, 0.11, 0.19, 0.03, 0.03, 0.09, 0.13, 0.17, 0.19, 0.19, 0.28, 0.14, 0.30, 0.07, 0.23, 0.32, 0.09, 0.07, 0.17, 0.15, 0.07, 0.15)
dindv_b<-c(0.14, 0.09, 0.12, 0.17, 0.27, 0.18, 0.11, 0.15, 0.05, 0.11, 0.20, 0.15, 0.14, 0.03, 0.05, 0.21, 0.25, 0.34, 0.26, 0.14, 0.12, 0.16, 0.23, 0.32, 0.18, 0.12, 0.50, 0.21, 0.26, 0.12)
dindv_c<-c(0.23283579, 0.03206649, 0.04862776, 0.05424779, 0.09153465, 0.19119154, 0.34639081)

#we transpose the network to be able to calculate the d' values for the individuals
netnulls.r2d_WBNn_a_t <- nullmodel(t(WBNs_a), N=1000, method="r2dtable")
netnulls.r2d_WBNn_b_t <- nullmodel(t(WBNs_b), N=1000, method="r2dtable")
netnulls.r2d_WBNn_c_t <- nullmodel(t(WBNs_c), N=1000, method="r2dtable")

dindv.nulls_a <- lapply(netnulls.r2d_WBNn_a_t, dfun) #
dindv.nulls_b <- lapply(netnulls.r2d_WBNn_b_t, dfun) #
dindv.nulls_c <- lapply(netnulls.r2d_WBNn_c_t, dfun) #

dindv_prime.nulls_a<-unlist(lapply(dindv.nulls_a, "[[", 1))
dindv_prime.nulls_b<-unlist(lapply(dindv.nulls_b, "[[", 1))
dindv_prime.nulls_c<-unlist(lapply(dindv.nulls_c, "[[", 1))

dindv_a<-sort(dindv_a, decreasing = TRUE)
dindv_b<-sort(dindv_b, decreasing = TRUE)
dindv_c<-sort(dindv_c, decreasing = TRUE)

dindv_a<-as.data.frame(dindv_a)
dindv_b<-as.data.frame(dindv_b)
dindv_c<-as.data.frame(dindv_c) 

#z-score dindv verify in the data where the values start to be significant 
dindv_a
net.score(0.22, dindv_prime.nulls_a); add.pvalues(1.99) # significative after 0.22 p 0.04
dindv_b
net.score(0.23, dindv_prime.nulls_b); add.pvalues(2.11)  # significative after 0.23 p 0.03
dindv_c
net.score(0.23, dindv_prime.nulls_c); add.pvalues(2.11) # significative after 0.23 p 0.03

ca_dindv<-ggplot(data=dindv_a, aes(y=dindv_a, x = seq(1, length(dindv_a)))) + geom_point(stat="identity", fill="black")+
  theme_classic() + labs(x = "Individuals", y = "d' values") + ylim(0, 0.5) + geom_hline(yintercept=0.07, linetype="dashed", color = "red") + Font_My_Theme + coord_flip()

cb_dindv<-ggplot(data=dindv_b, aes(y=dindv_b, x = seq(1, length(dindv_b)))) + geom_point(stat="identity", fill="black")+
  theme_classic() + labs(x = "Individuals", y = "d' values") + ylim(0, 0.5) +geom_hline(yintercept=0.08, linetype="dashed", color = "red") + Font_My_Theme + coord_flip()

cc_dindv<-ggplot(data=dindv_c, aes(y=dindv_c, x = seq(1, length(dindv_c)))) + geom_point(stat="identity", fill="black")+
  theme_classic() + labs(x = "Individuals", y = "d' values") + ylim(0, 0.5) + geom_hline(yintercept=0.06, linetype="dashed", color = "red") + Font_My_Theme + coord_flip()

#Ii
Ii_ca<-c(0.67, 0.54, 0.98, 0.83, 0.75, 0.80, 0.49, 0.18, 0.18, 0.93, 0.36, 0.85, 0.90, 0.73, 0.91, 0.93, 0.68, 0.86, 0.93, 0.77, 0.88, 
0.80, 0.77, 0.73, 0.91, 0.68, 0.83, 0.95, 0.98, 0.90, 0.83, 0.86, 0.73, 0.88, 0.86, 1, 0.21, 0.98, 0.90, 0.81, 0.96, 0.96, 0.96,
0.67, 0.91, 0.81, 0.93, 0.75, 0.68, 0.96, 0.83, 0.68, 0.96, 0.60, 0.81, 0.81, 0.68, 0.93, 0.91, 0.70)
Ii_cb<-c(0.83, 0.85, 0.80, 0.85, 0.80, 0.83, 0.93, 0.85, 0.85, 0.93, 0.90, 0.68, 0.91, 0.68, 0.98, 0.75, 0.85, 0.73, 0.67, 0.76, 0.86,
         0.74, 0.88, 0.88, 0.88, 0.96, 0.59, 0.54, 0.81, 0.73)
Ii_cc<-c(0.41, 0.34, 0.45, 0.64, 0.47, 0.65, 0.72)


#generate null model based on the values within the first and third interquantiles
dist_Il_ca<-runif(1000, min=0.72, max=0.91) 
dist_Il_cb<-runif(1000, min=0.74, max=0.88) 
dist_Il_cc<-runif(1000, min=0.43, max=0.64)
mean(as.matrix(Ii_ca)); mean(as.matrix(Ii_cb)); mean(as.matrix(Ii_cc))  

Ii_ca<-sort(Ii_ca, decreasing = TRUE)
Ii_cb<-sort(Ii_cb, decreasing = TRUE)
Ii_cc<-sort(Ii_cc, decreasing = TRUE)

Ii_ca<-as.data.frame(Ii_ca)
Ii_cb<-as.data.frame(Ii_cb)
Ii_cc<-as.data.frame(Ii_cc)

#z-score Ii index verify in the data where the p values start to be significant 
Ii_ca
net.score(0.93, dist_Il_ca); add.pvalues(2.03) # significative after 0.93 p 0.04
Ii_cb
net.score(0.90, dist_Il_cb); add.pvalues(2.24)  # significative after 0.88 p 0.02
Ii_cc
net.score(0.65, dist_Il_cc); add.pvalues(1.93) #  # significative after 1.93 p 0.05

ca_Iindv<-ggplot(data=Ii_ca, aes(y=Ii_ca, x = seq(1, length(Ii_ca)))) + geom_point(stat="identity", fill="black")+
  theme_classic() + labs(x = "Individuals", y = "Ii index") + ylim(0, 1) + geom_hline(yintercept=0.78, linetype="dashed", color = "red") + Font_My_Theme  + coord_flip()

cb_Iindv<-ggplot(data=Ii_cb, aes(y=Ii_cb, x = seq(1, length(Ii_cb)))) + geom_point(stat="identity", fill="black")+
  theme_classic() + labs(x = "Individuals", y = "Ii index") + ylim(0, 1) + geom_hline(yintercept=0.81, linetype="dashed", color = "red") + Font_My_Theme + coord_flip()

cc_Iindv<-ggplot(data=Ii_cc, aes(y=Ii_cc, x = seq(1, length(Ii_cc)))) + geom_point(stat="identity", fill="black")+
  theme_classic() + labs(x = "Individuals", y = "Ii index") + ylim(0, 1) + geom_hline(yintercept=0.52, linetype="dashed", color = "red") + Font_My_Theme + coord_flip() 


#######################################33
#plot d'behavior, d'indv and Ii index - Figure 4 Unedited 
ggarrange(ca_dbehav, cb_dbehav, cc_dbehav, ca_dindv, cb_dindv, cc_dindv, ca_Iindv, cb_Iindv, cc_Iindv, labels = c("A", "B", "C"), legend = "bottom", font.label = list(size = 12))


######
  #modularity
  modplot_a<-computeModules(WBNs_a) #Q=0.38 Qmax=0.80 Qnorm= 0.47
  #modplot_a<-metaComputeModules(WBNs_a, N=1, method="Beckett")
  #plotModuleWeb(modplot_a, rank=TRUE, weighted=TRUE)
  modplot_b<-computeModules(WBNs_b) #Q=0.37 Qmax=0.81 Qnorm= 0.45
  #modplot_b<-metaComputeModules(IBNn2, N=1000, method="Beckett")
  #plotModuleWeb(modplot2,  rank=TRUE, weighted=TRUE)
  modplot_c<-computeModules(WBNs_c) #Q=0.23 Qmax=0.71 Qnorm= 0.32
  #modplot_c<-metaComputeModules(IBNn3, N=1000, method="Beckett")
  #plotModuleWeb(modplot3,  rank=TRUE, weighted=TRUE)
  
  #netnulls.r2d_IBNn_a <- nullmodel(IBNn_a, N=1000, method="r2dtable")
  #netnulls.r2d_IBNn_b <- nullmodel(IBNn_b, N=1000, method="r2dtable")
  #netnulls.r2d_IBNn_c <- nullmodel(IBNn_c, N=1000, method="r2dtable")
  
  modules.nulls_a <- sapply(netnulls.r2d_WBNs_a, computeModules) #Q=0.16
  modules.nulls_b <- sapply(netnulls.r2d_WBNs_b, computeModules) #Q=0.17
  modules.nulls_c <- sapply(netnulls.r2d_WBNs_c , computeModules) #Q=0.15
  
  like.nulls_a <- sapply(modules.nulls_a, function(x) x@likelihood)
  like.nulls_b <- sapply(modules.nulls_b, function(x) x@likelihood)
  like.nulls_c <- sapply(modules.nulls_c, function(x) x@likelihood)
  
  rowMeans(cbind(like.nulls_a,  like.nulls_b, like.nulls_c))
           
  #qplot(like.nulls_a)+geom_vline(xintercept=0.38,col="red") + geom_histogram(bins=100) + Font_My_Theme + theme_classic()
  
#z-score modularity
  net.score(0.38, like.nulls_a); add.pvalues(0.40); # z-score 23.70 p = 3.605688e-124
  net.score(0.37, like.nulls_b); add.pvalues(0.36) # z-score 12.54169 p = 4.41479e-36
  net.score(0.23, like.nulls_c); add.pvalues(0.18) # z-score 6.25 p = 4.104527e-10
  
  
  ##### plot modularity 
  mod_Q<-read.table("Data S20.txt", header = TRUE)
  mod_Q$Data <- factor(mod_Q$Data, levels = c("Q", "null_model"))
  
  mod_Q_plot<-ggplot(mod_Q, aes(x=colonies, y=Q_value, fill=Data)) + 
    geom_bar(stat="identity", color="black", 
             position=position_dodge()) +
    geom_errorbar(aes(ymin=Q_value-sd_data, ymax=Q_value+sd_data), width=.2,
                  position=position_dodge(.9)) + theme_classic() + scale_fill_grey()
###############################################################################

# DOL index 
  ## R code to compute DOL statistics (Gorelick et al. 2004)
  
  DOLi.DOLt<-function (net){
    if(sum(net)>1)
      net<-net/sum(net)
    ColSum<-colSums(net)
    RowSum<-rowSums(net)
    pi.pj= RowSum%o%ColSum 
    I=sum(net*log(net/pi.pj),na.rm=T)
    Ht=-sum(RowSum*log(RowSum),na.rm=T)
    Hi=-sum(ColSum*log(ColSum),na.rm=T)
    DOLi=I/Ht
    DOLt=I/Hi
    return(c(DOLi,DOLt))
  }
  
DOLi.DOLt(WBNs_a) #0.44 0.21
DOLi.DOLt(WBNs_b) #0.38 0.21
DOLi.DOLt(WBNs_c) #0.17 0.14

DOL.nulls_a <- lapply(netnulls.r2d_WBNs_a, DOLi.DOLt) 
DOL.nulls_b <- lapply(netnulls.r2d_WBNs_b, DOLi.DOLt) 
DOL.nulls_c <- lapply(netnulls.r2d_WBNs_c, DOLi.DOLt) 

DOLi.nulls_a<-unlist(lapply(DOL.nulls_a, "[[", 1))
DOLi.nulls_b<-unlist(lapply(DOL.nulls_b, "[[", 1)) 
DOLi.nulls_c<-unlist(lapply(DOL.nulls_c, "[[", 1)) 

DOLt.nulls_a<-unlist(lapply(DOL.nulls_a, "[[", 2))
DOLt.nulls_b<-unlist(lapply(DOL.nulls_b, "[[", 2)) 
DOLt.nulls_c<-unlist(lapply(DOL.nulls_c, "[[", 2))

#DOLi
net.score(0.44, DOLi.nulls_a); add.pvalues(28.97569) # z-score 28.9 p = <0.0001 
net.score(0.38, DOLi.nulls_b); add.pvalues(15.43334) # z-score 15.4 p = <0.0001 
net.score(0.17, DOLi.nulls_c);add.pvalues(9.8) # z-score 9.8  p <0.0001 

#qplot(DOLi.nulls_n1)+geom_vline(xintercept=0.44,col="red") + geom_histogram(bins=100) + Font_My_Theme + theme_classic()

#DOLt
net.score(0.21, DOLt.nulls_a);add.pvalues(3.226994)  # z-score 28.17  p = 0.001
net.score(0.21, DOLt.nulls_b);add.pvalues(2.805712)# z-score 14.7 p = 0.005
net.score(0.14, DOLt.nulls_c);add.pvalues(7.220763)# z-score 9.8  p <0.0001 

#qplot(DOLt.nulls_n1)+geom_vline(xintercept=0.44,col="red") + geom_histogram(bins=100) + Font_My_Theme + theme_classic()

DOLi<-read.table("Data S18.txt", header=TRUE)

#plot DOLi
DOLi<-read.table("Data S18.txt", header = TRUE)
DOLi$Data <- factor(DOLi$Data, levels = c("DOLi", "null_model"))

DOLi_plot<-ggplot(DOLi, aes(x=colonies, y=DOL_value, fill=Data)) + 
  geom_bar(stat="identity", color="black", position=position_dodge()) +
  geom_errorbar(aes(ymin=DOL_value-sd_data, ymax=DOL_value+sd_data), width=.2, position=position_dodge(.9)) + 
  theme_classic() + scale_fill_grey()

DOLt<-read.table("Data S19.txt", header=TRUE)

#plot DOLt
DOLt<-read.table("Data S19.txt", header = TRUE)
DOLt$Data <- factor(DOLt$Data, levels = c("DOLt", "null_model"))

DOLt_plot <- ggplot(DOLt, aes(x=colonies, y=DOL_value, fill=Data)) + 
  geom_bar(stat="identity", color="black", position=position_dodge()) +
  geom_errorbar(aes(ymin=DOL_value-sd_data, ymax=DOL_value+sd_data), width=.2,
                position=position_dodge(.9)) + theme_classic() + scale_fill_grey()

#nestedness IBN
#netnulls.r2d_IBNn1 <- nullmodel(IBNn_a, N=1000, method="r2dtable")
#netnulls.r2d_IBNn2 <- nullmodel(IBNn_b, N=1000, method="r2dtable")
#netnulls.r2d_IBNn3 <- nullmodel(IBNn_c, N=1000, method="r2dtable")

obsn_a <- unlist(networklevel(WBNs_a, index="weighted NODF")) #WNODF 31.1 WINE 0.59
nulln_a <- unlist(sapply(netnulls.r2d_WBNs_a, networklevel, index="weighted NODF"))#42.4  #z-score -5 p = 0

obsn_b <- unlist(networklevel(WBNs_b, index="weighted NODF")) #WNODF 31.5 WINE 0.63
nulln_b <- unlist(sapply(netnulls.r2d_WBNs_b, networklevel, index="weighted NODF")) #37

obsn_c <- unlist(networklevel(WBNs_c, index="weighted NODF")) #WNODF 36.9 WINE 0.32
nulln_c <- unlist(sapply(netnulls.r2d_WBNs_c, networklevel, index="weighted NODF")) #46

#z-score WNODF
net.score(obsn_a, nulln_a); add.pvalues(-4.904562) #-4.94 p = <0.001
net.score(obsn_b, nulln_b); add.pvalues(-1.926077) #-1.92 p = 0.054
net.score(obsn_c, nulln_c); add.pvalues(-1.366671) #-1.36 p = 0.17

#####
WNODF<-read.table("Data S21.txt", header = TRUE)

WNODF$Data <- factor(WNODF$Data, levels = c("WNODF", "null_model"))

WNODF_plot<-ggplot(WNODF, aes(x=colonies, y=WNODF_value, fill=Data)) + 
  geom_bar(stat="identity", color="black", 
           position=position_dodge()) +
  geom_errorbar(aes(ymin=WNODF_value-sd_data, ymax=WNODF_value+sd_data), width=.2,
                position=position_dodge(.9)) + theme_classic() +   scale_fill_grey()
#######################################

plot(density(nulln_a), xlim=c(min(obsn_a, min(nulln_a)), max(obsn_a, max(nulln_a))), 
     main="comparison of observed with null model Patefield")
abline(v=obsn_a, col="red", lwd=2)    

plot(density(nulln_b), xlim=c(min(obsn_b, min(nulln_b)), max(obsn_b, max(nulln_b))), 
     main="comparison of observed with null model Patefield")
abline(v=obsn_b, col="red", lwd=2)

plot(density(nulln_c), xlim=c(min(obsn_c, min(nulln_c)), max(obsn_c, max(nulln_c))), 
     main="comparison of observed with null model Patefield")
abline(v=obsn_c, col="red", lwd=2)


#plot d'behavior, d'indv and Ii index - Unedited figure 2
ggarrange(H2plot, DOLi_plot, DOLt_plot, mod_Q_plot, WNODF_plot, font.label = list(size = 12),  widths = c(2), ncol = 2, nrow =3)
```
