## Appendix, as digital supplementary material to this article for "Worker-behavior and behavior-behavior interaction networks in the trap-jaw ant *Odontomachus chelifer* (Latreille, 1802) (Hymenoptera: Formicidae)": Script S3.R

```
#BTN1 open files from Data S9 A
BTN1<-read.table("BTN1.txt", header=TRUE)
BTN2<-read.table("BTN2.txt", header=TRUE)
BTN3<-read.table("BTN3.txt", header=TRUE)
BTN4<-read.table("BTN4.txt", header=TRUE)
BTN5<-read.table("BTN5.txt", header=TRUE)
BTN6<-read.table("BTN6.txt", header=TRUE)
BTN7<-read.table("BTN7.txt", header=TRUE)
BTN8<-read.table("BTN8.txt", header=TRUE)
BTN9<-read.table("BTN9.txt", header=TRUE)
BTN10<-read.table("BTN10.txt", header=TRUE)
BTN11<-read.table("BTN11.txt", header=TRUE)
BTN12<-read.table("BTN12.txt", header=TRUE)
BTN13<-read.table("BTN13.txt", header=TRUE)
BTN14<-read.table("BTN14.txt", header=TRUE)
BTN15<-read.table("BTN15.txt", header=TRUE)
BTN16<-read.table("BTN16.txt", header=TRUE)
BTN17<-read.table("BTN17.txt", header=TRUE)
BTN18<-read.table("BTN18.txt", header=TRUE)
BTN19<-read.table("BTN19.txt", header=TRUE)
BTN20<-read.table("BTN20.txt", header=TRUE)
BTN21<-read.table("BTN21.txt", header=TRUE)
BTN22<-read.table("BTN22.txt", header=TRUE)
BTN23<-read.table("BTN23.txt", header=TRUE)
BTN24<-read.table("BTN24.txt", header=TRUE)
BTN25<-read.table("BTN25.txt", header=TRUE)
BTN26<-read.table("BTN26.txt", header=TRUE)
BTN27<-read.table("BTN27.txt", header=TRUE)
BTN28<-read.table("BTN28.txt", header=TRUE)
BTN29<-read.table("BTN29.txt", header=TRUE)
BTN30<-read.table("BTN30.txt", header=TRUE)
BTN31<-read.table("BTN31.txt", header=TRUE)
BTN32<-read.table("BTN32.txt", header=TRUE)
BTN33<-read.table("BTN33.txt", header=TRUE)
BTN34<-read.table("BTN34.txt", header=TRUE)
BTN35<-read.table("BTN35.txt", header=TRUE)
BTN36<-read.table("BTN36.txt", header=TRUE)
BTN37<-read.table("BTN37.txt", header=TRUE)
BTN38<-read.table("BTN38.txt", header=TRUE)
BTN39<-read.table("BTN39.txt", header=TRUE)
BTN40<-read.table("BTN40.txt", header=TRUE)
BTN41<-read.table("BTN41.txt", header=TRUE)
BTN42<-read.table("BTN42.txt", header=TRUE)
BTN43<-read.table("BTN43.txt", header=TRUE)
BTN44<-read.table("BTN44.txt", header=TRUE)
BTN45<-read.table("BTN45.txt", header=TRUE)
BTN46<-read.table("BTN46.txt", header=TRUE)
BTN47<-read.table("BTN47.txt", header=TRUE)
BTN48<-read.table("BTN48.txt", header=TRUE)
BTN49<-read.table("BTN49.txt", header=TRUE)
BTN50<-read.table("BTN50.txt", header=TRUE)
BTN51<-read.table("BTN51.txt", header=TRUE)
BTN52<-read.table("BTN52.txt", header=TRUE)
BTN53<-read.table("BTN53.txt", header=TRUE)
BTN54<-read.table("BTN54.txt", header=TRUE)
BTN55<-read.table("BTN55.txt", header=TRUE)
BTN56<-read.table("BTN56.txt", header=TRUE)
BTN57<-read.table("BTN57.txt", header=TRUE)
BTN58<-read.table("BTN58.txt", header=TRUE)
BTN59<-read.table("BTN59.txt", header=TRUE)
BTN60<-read.table("BTN60.txt", header=TRUE)

#BTNa
BTNa_total<-BTN1+BTN2+BTN3+BTN4+BTN5+BTN6+BTN7+BTN8+BTN9+BTN10+BTN11+BTN12+BTN13+BTN14+BTN15+BTN16+BTN17+BTN18+BTN19+BTN20+
  BTN21+BTN22+BTN23+BTN24+BTN25+BTN26+BTN27+BTN28+BTN29+BTN30+BTN31+BTN32+BTN33+BTN34+BTN35+BTN36+BTN37+BTN38+BTN39+BTN40+
  BTN41+BTN42+BTN43+BTN44+BTN45+BTN46+BTN47+BTN48+BTN50+BTN51+BTN52+BTN53+BTN54+BTN55+BTN56+BTN57+BTN58+BTN59+BTN60
#################################

#BTNb open files from Data S9 B
BTN1<-read.table("BTN1.txt", header=TRUE)
BTN2<-read.table("BTN2.txt", header=TRUE)
BTN3<-read.table("BTN3.txt", header=TRUE)
BTN4<-read.table("BTN4.txt", header=TRUE)
BTN5<-read.table("BTN5.txt", header=TRUE)
BTN6<-read.table("BTN6.txt", header=TRUE)
BTN7<-read.table("BTN7.txt", header=TRUE)
BTN8<-read.table("BTN8.txt", header=TRUE)
BTN9<-read.table("BTN9.txt", header=TRUE)
BTN10<-read.table("BTN10.txt", header=TRUE)
BTN11<-read.table("BTN11.txt", header=TRUE)
BTN12<-read.table("BTN12.txt", header=TRUE)
BTN13<-read.table("BTN13.txt", header=TRUE)
BTN14<-read.table("BTN14.txt", header=TRUE)
BTN15<-read.table("BTN15.txt", header=TRUE)
BTN16<-read.table("BTN16.txt", header=TRUE)
BTN17<-read.table("BTN17.txt", header=TRUE)
BTN18<-read.table("BTN18.txt", header=TRUE)
BTN19<-read.table("BTN19.txt", header=TRUE)
BTN20<-read.table("BTN20.txt", header=TRUE)
BTN21<-read.table("BTN21.txt", header=TRUE)
BTN22<-read.table("BTN22.txt", header=TRUE)
BTN23<-read.table("BTN23.txt", header=TRUE)
BTN24<-read.table("BTN24.txt", header=TRUE)
BTN25<-read.table("BTN25.txt", header=TRUE)
BTN26<-read.table("BTN26.txt", header=TRUE)
BTN27<-read.table("BTN27.txt", header=TRUE)
BTN28<-read.table("BTN28.txt", header=TRUE)
BTN29<-read.table("BTN29.txt", header=TRUE)
BTN30<-read.table("BTN30.txt", header=TRUE)

#BTNC2
BTNb_total<-BTN1+BTN2+BTN3+BTN4+BTN5+BTN6+BTN7+BTN8+BTN9+BTN10+BTN11+BTN12+BTN13+BTN14+BTN15+BTN16+BTN17+BTN18+BTN19+BTN20+
  BTN21+BTN22+BTN23+BTN24+BTN25+BTN26+BTN27+BTN28+BTN29+BTN30

#######################
#BTN3 open files from Data S9 C
BTN1<-read.table("BTN1.txt", header=TRUE)
BTN2<-read.table("BTN2.txt", header=TRUE)
BTN3<-read.table("BTN3.txt", header=TRUE)
BTN4<-read.table("BTN4.txt", header=TRUE)
BTN5<-read.table("BTN5.txt", header=TRUE)
BTN6<-read.table("BTN6.txt", header=TRUE)
BTN7<-read.table("BTN7.txt", header=TRUE)

#BTNC3
BTNc_total<-BTN1+BTN2+BTN3+BTN4+BTN5+BTN6+BTN7

######################################

#BTNaI open files from Data S10 A
I1<-read.table("I1.txt", header=TRUE)
I2<-read.table("I2.txt", header=TRUE)
I3<-read.table("I3.txt", header=TRUE)
I4<-read.table("I4.txt", header=TRUE)
I5<-read.table("I5.txt", header=TRUE)
I6<-read.table("I6.txt", header=TRUE)
I7<-read.table("I7.txt", header=TRUE)
I8<-read.table("I8.txt", header=TRUE)
I9<-read.table("I9.txt", header=TRUE)
I10<-read.table("I10.txt", header=TRUE)
I11<-read.table("I11.txt", header=TRUE)
I12<-read.table("I12.txt", header=TRUE)
I13<-read.table("I13.txt", header=TRUE)
I14<-read.table("I14.txt", header=TRUE)
I15<-read.table("I15.txt", header=TRUE)
I16<-read.table("I16.txt", header=TRUE)
I17<-read.table("I17.txt", header=TRUE)
I18<-read.table("I18.txt", header=TRUE)
I19<-read.table("I19.txt", header=TRUE)
I20<-read.table("I20.txt", header=TRUE)
I21<-read.table("I21.txt", header=TRUE)
I22<-read.table("I22.txt", header=TRUE)
I23<-read.table("I23.txt", header=TRUE)
I24<-read.table("I24.txt", header=TRUE)
I25<-read.table("I25.txt", header=TRUE)
I26<-read.table("I26.txt", header=TRUE)
I27<-read.table("I27.txt", header=TRUE)
I28<-read.table("I28.txt", header=TRUE)
I29<-read.table("I29.txt", header=TRUE)
I30<-read.table("I30.txt", header=TRUE)
I31<-read.table("I31.txt", header=TRUE)
I32<-read.table("I32.txt", header=TRUE)
I33<-read.table("I33.txt", header=TRUE)
I34<-read.table("I34.txt", header=TRUE)
I35<-read.table("I35.txt", header=TRUE)
I36<-read.table("I36.txt", header=TRUE)
I37<-read.table("I37.txt", header=TRUE)
I38<-read.table("I38.txt", header=TRUE)
I39<-read.table("I39.txt", header=TRUE)
I40<-read.table("I40.txt", header=TRUE)
I41<-read.table("I41.txt", header=TRUE)
I42<-read.table("I42.txt", header=TRUE)
I43<-read.table("I43.txt", header=TRUE)
I44<-read.table("I44.txt", header=TRUE)
I45<-read.table("I45.txt", header=TRUE)
I46<-read.table("I46.txt", header=TRUE)
I47<-read.table("I47.txt", header=TRUE)
I48<-read.table("I48.txt", header=TRUE)
I49<-read.table("I49.txt", header=TRUE)
I50<-read.table("I50.txt", header=TRUE)
I51<-read.table("I51.txt", header=TRUE)
I52<-read.table("I52.txt", header=TRUE)
I53<-read.table("I53.txt", header=TRUE)
I54<-read.table("I54.txt", header=TRUE)
I55<-read.table("I55.txt", header=TRUE)
I56<-read.table("I56.txt", header=TRUE)
I57<-read.table("I57.txt", header=TRUE)
I58<-read.table("I58.txt", header=TRUE)
I59<-read.table("I59.txt", header=TRUE)
I60<-read.table("I60.txt", header=TRUE)

BTNCaI_total<-I1+I2+I3+I4+I5+I6+I7+I8+I9+I10+I11+I12+I13+I14+I15+I16+I17+I18+I19+I20+
  I21+I22+I23+I24+I25+I26+I27+I28+I29+I30+I31+I32+I33+I34+I35+I36+I37+I38+I39+I40+
  I41+I42+I43+I44+I45+I46+I47+I48+I49+I50+I51+I52+I53+I54+I55+I56+I57+I58+I59+I60

#BTNbI open files from Data S10 B
I1<-read.table("I1.txt", header=TRUE)
I2<-read.table("I2.txt", header=TRUE)
I3<-read.table("I3.txt", header=TRUE)
I4<-read.table("I4.txt", header=TRUE)
I5<-read.table("I5.txt", header=TRUE)
I6<-read.table("I6.txt", header=TRUE)
I7<-read.table("I7.txt", header=TRUE)
I8<-read.table("I8.txt", header=TRUE)
I9<-read.table("I9.txt", header=TRUE)
I10<-read.table("I10.txt", header=TRUE)
I11<-read.table("I11.txt", header=TRUE)
I12<-read.table("I12.txt", header=TRUE)
I13<-read.table("I13.txt", header=TRUE)
I14<-read.table("I14.txt", header=TRUE)
I15<-read.table("I15.txt", header=TRUE)
I16<-read.table("I16.txt", header=TRUE)
I17<-read.table("I17.txt", header=TRUE)
I18<-read.table("I18.txt", header=TRUE)
I19<-read.table("I19.txt", header=TRUE)
I20<-read.table("I20.txt", header=TRUE)
I21<-read.table("I21.txt", header=TRUE)
I22<-read.table("I22.txt", header=TRUE)
I23<-read.table("I23.txt", header=TRUE)
I24<-read.table("I24.txt", header=TRUE)
I25<-read.table("I25.txt", header=TRUE)
I26<-read.table("I26.txt", header=TRUE)
I27<-read.table("I27.txt", header=TRUE)
I28<-read.table("I28.txt", header=TRUE)
I29<-read.table("I29.txt", header=TRUE)
I30<-read.table("I30.txt", header=TRUE)
####################

BTNbI_total<-I1+I2+I3+I4+I5+I6+I7+I8+I9+I10+I11+I12+I13+I14+I15+I16+I17+I18+I19+I20+
  I21+I22+I23+I24+I25+I26+I27+I28+I29+I30


#BTN3I open files from Data S10 C 
I1<-read.table("I1.txt", header=TRUE)
I2<-read.table("I2.txt", header=TRUE)
I3<-read.table("I3.txt", header=TRUE)
I4<-read.table("I4.txt", header=TRUE)
I5<-read.table("I5.txt", header=TRUE)
I6<-read.table("I6.txt", header=TRUE)
I7<-read.table("I7.txt", header=TRUE)

#BTNC3I

BTNCc_Itotal<- I1+ I2+ I3 + I4 + I5+ I6 + I7


#betweness centarlity values - above 3 quartile 
#c1
c1<-c(14,36,0,0,14,0,0,0,0,0,0)
quantile(c1, 0.75)

#c2
c2<-c(0,17,0,8,5,2,3,0)
quantile(c2, 0.75)

#c3
c3<-c(5,0,0,0,0,5,6)
quantile(c3,0.75)


#degree centrality - above quantile 3 
#c1 out
c1out<-c(13.5,24.5,9.3,17.5,11.7,21.5,14.1,9.9,10.3,10.7,22.6)
quantile(c1out,0.75); quantile(c1out,0.25)

#c2 out
c2out<-c(6.5,11.1,9.4,20.3,17.5,20,17.8,4.7,4.8)
quantile(c2out,0.75); quantile(c2out,0.25)

#c3 out
c3out<-c(6.5,7,4.1,6.9,2.5,11,12.6)
quantile(c3out,0.75); quantile(c3out,0.25)

#c1 in 
c1in<-c(37,36.3,3.3,18.2,12.6,18.1,11.7,4.3,4.5,2.8,17)
quantile(c1in,0.75); quantile(c1in,0.25)

#c2 in
c2in<-c(13.5,27.2,13.6,17.6,10.3,17.3,7.3,2.1,3.2)
quantile(c2in,0.75); quantile(c2in,0.25)

#c3 in
c3in<-c(11.9,8.9,2.9,7.3,0,8.8,10.8)
quantile(c3in,0.75); quantile(c3in,0.25)
```
