## Appendix, as digital supplementary material to this article for "Worker-behavior and behavior-behavior interaction networks in the trap-jaw ant *Odontomachus chelifer* (Latreille, 1802) (Hymenoptera: Formicidae)": Script S4.R

```
# colony a = BTNC1 graph and UNODF
library(igraph)
library(RVAideMemoire)
library(BBmisc)
library(unodf)

full_BBN_a <-read.table("Data S11.txt", header=TRUE)
BTNC1total<-as.matrix(full_BBN_a) #from the script read_BTNS
#remove at 
BTNC1totalat <- BTNC1total[-c(3), ]
BTNC1totalat <- BTNC1totalat[,-3 ]

BTNC1totalig<-graph_from_adjacency_matrix(BTNC1total)
BTNC1totalig<-graph_from_adjacency_matrix(BTNC1totalat)

l <- layout_in_circle(BTNC1totalig)

BTNC1totaligs_b<-simplify(BTNC1totalig, remove.loops = TRUE, remove.multiple=TRUE)
#BTN1centrality<-betweenness(BTNC1totaligs_b,  normalized = TRUE)

node.size<-setNames(c(14, 36, 0, 0, 14, 0, 0, 0, 0, 0),c("WL", "SG","G", "BG", "BC", "CB", "CD", "FP", "CF", "FD"))

E(BTNC1totaligs_b)$width <- 1+E(BTNC1totaligs_b)/5
E(BTNC1totaligs_b)$weight <- E(BTNC1totaligs_b)$width

plot(BTNC1totaligs_b, layout=l, edge.arrow.size=.7, edge.curved=.15, edge.color="black", vertex.color="white",
     vertex.label.color="black", mark.groups=communities(BTN1mod), vertex.size=(node.size*1.5))

#####################################################3
BTNC1totaligs<-as.undirected(BTNC1totaligs_b, mode = c("collapse"), edge.attr.comb = "sum")
BTN1mod<-cluster_louvain(BTNC1totaligs) #0.22

#E(BTNC1totaligs)$weight <- E(BTNC1totaligs)$width

plot(BTNC1totaligs_b, layout=l, edge.arrow.size=.7, edge.curved=.15, edge.color="black", vertex.color="white",
     vertex.label.color="black", mark.groups=communities(BTN1mod), vertex.size=(node.size*1.5))

#UNODF BTN1###########################################################
BTNC1total_norm<-BTNC1totalat/sum(BTNC1totalat); BTNC1total_norm

BTNC1total_bin<-cutoff(BTNC1total_norm, cut = seq(0, 0)); BTNC1total_bin_0.01<-cutoff(BTNC1total_norm, cut = seq(0.01, 0))

#####UNODF BTNC1total_bin
BTNC1total_bin<-as.data.frame(BTNC1total_bin); BTNC1total_bin<-as.matrix(BTNC1total_bin)

BTNC1total_bin_unodf<-unodf(BTNC1total_bin) #UNODFc 0.5432099  #  UNODFr 0.4691358

####UNODF BTNC1total_bin_0.1
BTNC1total_bin_0.01<-as.data.frame(BTNC1total_bin_0.01); BTNC1total_bin_0.01<-as.matrix(BTNC1total_bin_0.01)

BTNC1total_bin_unodf_0.01<-unodf(BTNC1total_bin_0.01) #UNODFc 0.1333333  #  UNODFr 0.1333333

######################################## UNODF null model 
iter=1000
netnulls.BTNC1total_bin<-null2(BTNC1total_bin, iter); netnulls.BTNC1total_bin<-array_tree(netnulls.BTNC1total_bin, 3)

netnulls.BTNC1total_bin_0.01<-null2(BTNC1total_bin_0.01, iter, symmetric=TRUE); netnulls.BTNC1total_bin_0.01<-array_tree(netnulls.BTNC1total_bin_0.01, 3)

unodf.nulls_n1_bin <- lapply(netnulls.BTNC1total_bin, unodf); unodf.nulls_n1_bin_0.01 <- lapply(netnulls.BTNC1total_bin_0.01, unodf)

##########################################################
unodf_nullmodel_n1_Fc<-lapply(unodf.nulls_n1_bin, function(x) x[1,2]) #UNODFc
unodf_nullmodel_n1_Fr<-lapply(unodf.nulls_n1_bin, function(x) x[1,2]) #UNODFr

unodf_nullmodel_n1_Fc<-as.data.frame(unlist(unodf_nullmodel_n1_Fc)); unodf_nullmodel_n1_Fc<-as.numeric(unlist(unodf_nullmodel_n1_Fc)) #

net.zscore = function(obsval, nullval) {
  (obsval - mean(nullval))/sd(nullval)  
} 

zscore<-net.zscore(0.5432099, unodf_nullmodel_n1_Fc); zscore #2.59
net.metric.pvalue <- 2*pnorm(-abs(zscore)); net.metric.pvalue # 0.009

unodf_nullmodel_n1_Fr<-as.data.frame(unlist(unodf_nullmodel_n1_Fr)); unodf_nullmodel_n1_Fr<-as.numeric(unlist(unodf_nullmodel_n1_Fr)) #

zscore<-net.zscore(0.4691358, unodf_nullmodel_n1_Fr); zscore
net.metric.pvalue <- 2*pnorm(-abs(zscore)); net.metric.pvalue # 
########################

unodf_nullmodel_n1_0.01_Fc<-lapply(unodf.nulls_n1_bin_0.01, function(x) x[1,2]) #UNODFc
unodf_nullmodel_n1_0.01_Fr<-lapply(unodf.nulls_n1_bin_0.01, function(x) x[1,2]) #UNODFr

unodf_nullmodel_n1_0.01_Fc<-as.data.frame(unlist(unodf_nullmodel_n1_0.01_Fc)); unodf_nullmodel_n1_0.01_Fc<-as.numeric(unlist(unodf_nullmodel_n1_0.01_Fc)) #

zscore<-net.zscore(0.1333333, unodf_nullmodel_n1_0.01_Fc); zscore
net.metric.pvalue <- 2*pnorm(-abs(zscore)); net.metric.pvalue # 

unodf_nullmodel_n1_0.01_Fr<-as.data.frame(unlist(unodf_nullmodel_n1_0.01_Fr)); unodf_nullmodel_n1_0.01_Fr<-as.numeric(unlist(unodf_nullmodel_n1_0.01_Fr)) #

zscore<-net.zscore(0.1333333, unodf_nullmodel_n1_0.01_Fr); zscore
net.metric.pvalue <- 2*pnorm(-abs(zscore)); net.metric.pvalue # 
```
