## Appendix, as digital supplementary material to this article for "Worker-behavior and behavior-behavior interaction networks in the trap-jaw ant *Odontomachus chelifer* (Latreille, 1802) (Hymenoptera: Formicidae)": Script S5.R

```
# colony b = BTNC2 graph and UNODF
library(igraph)
library(RColorBrewer)
library(BBmisc)
library(unodf)

BTNC2total<-as.matrix(BTNb_total) #from the script read_BTNS

#remove cdfp
BTNC2totalcdfp <- BTNC2total[-c(8), ]
BTNC2totalcdfp <- BTNC2totalcdfp[,-8]

BTNC2totalig<-graph_from_adjacency_matrix(BTNC2total)
BTNC2totalig<-graph_from_adjacency_matrix(BTNC2totalcdfp)

l <- layout_in_circle(BTNC2totalig)

#E(BTNC2totalig)$width <- 1+E(BTNC2totalig)/45
#E(BTNC2totalig)$weight <- E(BTNC2totalig)$width

BTNC2totaligs_b<-simplify(BTNC2totalig, remove.loops = TRUE, remove.multiple=TRUE)
BTN2centrality<-betweenness(BTNC2totaligs_b,  normalized = TRUE)

node.size<-setNames(c(0, 17, 0, 8, 5, 2, 3, 0),c("WL", "SG","AT", "G", "BG", "BC", "CB", "FP"))

E(BTNC2totaligs_b)$width <- 1+E(BTNC2totaligs_b)/5
E(BTNC2totaligs_b)$weight <- E(BTNC2totaligs_b)$width

################################333

BTNC2totaligs<-as.undirected(BTNC2totaligs_b, mode = c("collapse"), edge.attr.comb = "sum")
BTN2mod <- cluster_louvain(BTNC2totaligs) #0.21

E(BTNC2totaligs)$weight <- E(BTNC2totaligs)$width

plot(BTNC2totaligs_b, layout=l, edge.arrow.size=.7, edge.curved=.15, edge.color="black", vertex.color="white",
     vertex.label.color="black", mark.groups=communities(BTN2mod), vertex.size=(node.size*1.5))

#plot(BTNC2totaligs, layout=l, edge.arrow.size=.8, edge.curved=.15, edge.color="black", vertex.color="white",
#     vertex.label.color="black", mark.groups=communities(BTN2mod), vertex.size=(node.size*2.5))

#UNODF BTN2###########################################################
BTNC2total_norm<-BTNC2totalcdfp/sum(BTNC2totalcdfp)
BTNC2total_norm
BTNC2total_bin<-cutoff(BTNC2total_norm, cut = seq(0, 0))
BTNC2total_bin_0.01<-cutoff(BTNC2total_norm, cut = seq(0.01, 0))

#####UNODF BTNC2total_bin
BTNC2total_bin<-as.data.frame(BTNC2total_bin)
BTNC2total_bin<-as.matrix(BTNC2total_bin)

unodf(BTNC2total_bin) #UNODFc 0.44  #  UNODFr 0.48

####UNODF BTNC2total_bin_0.1
BTNC2total_bin_0.01<-as.data.frame(BTNC2total_bin_0.01)
BTNC2total_bin_0.01<-as.matrix(BTNC2total_bin_0.01)

unodf(BTNC2total_bin_0.01) #UNODFc 0.30  #  UNODFr 0.36

######################################## UNODF null model 
iter=1000
netnulls.BTNC2total_bin<-null2(BTNC2total_bin, iter); netnulls.BTNC2total_bin<-array_tree(netnulls.BTNC2total_bin, 3)

netnulls.BTNC2total_bin_0.01<-null2(BTNC2total_bin_0.01, iter); netnulls.BTNC2total_bin_0.01<-array_tree(netnulls.BTNC2total_bin_0.01, 3)

unodf.nulls_n2_bin <- lapply(netnulls.BTNC2total_bin, unodf); unodf.nulls_n2_bin_0.01 <- lapply(netnulls.BTNC2total_bin_0.01, unodf)

##########################################################
unodf_nullmodel_n2_Fc<-lapply(unodf.nulls_n2_bin, function(x) x[1,2]) #UNODFc
unodf_nullmodel_n2_Fr<-lapply(unodf.nulls_n2_bin, function(x) x[1,2]) #UNODFr

unodf_nullmodel_n2_Fc<-as.data.frame(unlist(unodf_nullmodel_n2_Fc)); unodf_nullmodel_n2_Fc<-as.numeric(unlist(unodf_nullmodel_n2_Fc)) #

net.zscore = function(obsval, nullval) {
  (obsval - mean(nullval))/sd(nullval)  
} 

zscore<-net.zscore(0.44, unodf_nullmodel_n2_Fc); zscore
net.metric.pvalue <- 2*pnorm(-abs(zscore)); net.metric.pvalue # 

unodf_nullmodel_n2_Fr<-as.data.frame(unlist(unodf_nullmodel_n2_Fr)); unodf_nullmodel_n2_Fr<-as.numeric(unlist(unodf_nullmodel_n2_Fr)) #

zscore<-net.zscore(0.48, unodf_nullmodel_n2_Fr); zscore
net.metric.pvalue <- 2*pnorm(-abs(zscore)); net.metric.pvalue # 
########################

unodf_nullmodel_n2_0.01_Fc<-lapply(unodf.nulls_n2_bin_0.01, function(x) x[1,2]) #UNODFc
unodf_nullmodel_n2_0.01_Fr<-lapply(unodf.nulls_n2_bin_0.01, function(x) x[1,2]) #UNODFr

unodf_nullmodel_n2_0.01_Fc<-as.data.frame(unlist(unodf_nullmodel_n2_0.01_Fc)); unodf_nullmodel_n2_0.01_Fc<-as.numeric(unlist(unodf_nullmodel_n2_0.01_Fc)) #

zscore<-net.zscore(0.30, unodf_nullmodel_n2_0.01_Fc); zscore
net.metric.pvalue <- 2*pnorm(-abs(zscore)); net.metric.pvalue # 

unodf_nullmodel_n2_0.01_Fr<-as.data.frame(unlist(unodf_nullmodel_n2_0.01_Fr)); unodf_nullmodel_n2_0.01_Fr<-as.numeric(unlist(unodf_nullmodel_n2_0.01_Fr)) #

zscore<-net.zscore(0.36, unodf_nullmodel_n2_0.01_Fr); zscore
net.metric.pvalue <- 2*pnorm(-abs(zscore)); net.metric.pvalue # 
```
