## Appendix, as digital supplementary material to this article for "Worker-behavior and behavior-behavior interaction networks in the trap-jaw ant *Odontomachus chelifer* (Latreille, 1802) (Hymenoptera: Formicidae)": Script S6.R

```
## colony c = BTNC3 graph and UNODF
library(igraph)
library(RColorBrewer)
library(BBmisc)
library(unodf)

full_BBN_c <-read.table("Data S15.txt", header=TRUE)
BTNC3total<-as.matrix(full_BBN_c) #from the script read_BTNS
BTNC3totalig<-graph_from_adjacency_matrix(BTNC3total)

l <- layout_in_circle(BTNC3totalig)

#E(BTNC3totalig)$width <- 1+E(BTNC3totalig)/45
#E(BTNC3totalig)$weight <- E(BTNC3totalig)$width

BTNC3totaligs_b<-simplify(BTNC3totalig, remove.loops = TRUE, remove.multiple=TRUE)
BTN3centrality<-betweenness(BTNC3totaligs_b,  normalized = TRUE)

node.size<-setNames(c(5, 0, 0, 0, 0, 0, 5, 6),c("WL", "SG","AT", "CF", "BC", "FP", "FD"))

E(BTNC3totaligs_b)$width <- 1+E(BTNC3totaligs_b)/5
E(BTNC3totaligs_b)$weight <- E(BTNC3totaligs_b)$width

###########################3
BTNC3totaligs<-as.undirected(BTNC3totaligs_b, mode = c("collapse"), edge.attr.comb = "sum")
BTN3mod <- cluster_louvain(BTNC3totaligs) # 

E(BTNC3totaligs)$weight <- E(BTNC3totaligs)$width

plot(BTNC3totaligs_b, layout=l, edge.arrow.size=.7, edge.curved=.15, edge.color="black", vertex.color="white",
     vertex.label.color="black", mark.groups=communities(BTN3mod), vertex.size=(node.size*1.5))

#plot(BTNC3totaligs, layout=l, edge.arrow.size=.7, edge.curved=.15, edge.color="black", vertex.color="white",
#     vertex.label.color="black", mark.groups=communities(BTN3mod), vertex.size=(BTN3centrality*2.5))

#UNODF BTN3###########################################################
BTNC3total_norm<-BTNC3total/sum(BTNC3total)
BTNC3total_norm
BTNC3total_bin<-cutoff(BTNC3total_norm, cut = seq(0, 0))
BTNC3total_bin_0.01<-cutoff(BTNC3total_norm, cut = seq(0.01, 0))

#####UNODF BTNC3total_bin
BTNC3total_bin<-as.data.frame(BTNC3total_bin)
BTNC3total_bin<-as.matrix(BTNC3total_bin)

unodf(BTNC3total_bin) #UNODFc 0.64  #  UNODFr 0.80

####UNODF BTNC1total_bin_0.1

netnulls.BTNC3total_bin_0.01<-null2(BTNC3total_bin_0.01, iter, symmetric=TRUE); netnulls.BTNC3total_bin_0.01<-array_tree(netnulls.BTNC3total_bin_0.01, 3)
BTNC3total_bin_0.01<-as.data.frame(BTNC3total_bin_0.01)
BTNC3total_bin_0.01<-as.matrix(BTNC3total_bin_0.01)

unodf(BTNC3total_bin_0.01) #UNODFc 0.25  #  UNODFr 0.25
##################################################################

######################################## UNODF null model 
iter=1000
netnulls.BTNC3total_bin<-null2(BTNC3total_bin, iter); netnulls.BTNC3total_bin<-array_tree(netnulls.BTNC3total_bin, 3)

netnulls.BTNC3total_bin_0.01<-null2(BTNC3total_bin_0.01, iter); netnulls.BTNC3total_bin_0.01<-array_tree(netnulls.BTNC3total_bin_0.01, 3)

unodf.nulls_n3_bin <- lapply(netnulls.BTNC3total_bin, unodf); unodf.nulls_n3_bin_0.01 <- lapply(netnulls.BTNC3total_bin_0.01, unodf)

##########################################################
unodf_nullmodel_n3_Fc<-lapply(unodf.nulls_n3_bin, function(x) x[1,2]) #UNODFc
unodf_nullmodel_n3_Fr<-lapply(unodf.nulls_n3_bin, function(x) x[1,2]) #UNODFr

unodf_nullmodel_n3_Fc<-as.data.frame(unlist(unodf_nullmodel_n3_Fc)); unodf_nullmodel_n3_Fc<-as.numeric(unlist(unodf_nullmodel_n3_Fc)) #

net.zscore = function(obsval, nullval) {
  (obsval - mean(nullval))/sd(nullval)  
} 

zscore<-net.zscore(0.64, unodf_nullmodel_n3_Fc); zscore
net.metric.pvalue <- 2*pnorm(-abs(zscore)); net.metric.pvalue # 

unodf_nullmodel_n3_Fr<-as.data.frame(unlist(unodf_nullmodel_n3_Fr)); unodf_nullmodel_n3_Fr<-as.numeric(unlist(unodf_nullmodel_n3_Fr)) #

zscore<-net.zscore(0.80, unodf_nullmodel_n3_Fr); zscore
net.metric.pvalue <- 2*pnorm(-abs(zscore)); net.metric.pvalue # 
########################

unodf_nullmodel_n3_0.01_Fc<-lapply(unodf.nulls_n3_bin_0.01, function(x) x[1,2]) #UNODFc
unodf_nullmodel_n3_0.01_Fr<-lapply(unodf.nulls_n3_bin_0.01, function(x) x[1,2]) #UNODFr

unodf_nullmodel_n3_0.01_Fc<-as.data.frame(unlist(unodf_nullmodel_n3_0.01_Fc)); unodf_nullmodel_n3_0.01_Fc<-as.numeric(unlist(unodf_nullmodel_n3_0.01_Fc)) #

zscore<-net.zscore(0.25, unodf_nullmodel_n3_0.01_Fc); zscore
net.metric.pvalue <- 2*pnorm(-abs(zscore)); net.metric.pvalue # 

unodf_nullmodel_n3_0.01_Fr<-as.data.frame(unlist(unodf_nullmodel_n3_0.01_Fr)); unodf_nullmodel_n3_0.01_Fr<-as.numeric(unlist(unodf_nullmodel_n3_0.01_Fr)) #

zscore<-net.zscore(0.25, unodf_nullmodel_n2_0.01_Fr); zscore
net.metric.pvalue <- 2*pnorm(-abs(zscore)); net.metric.pvalue # 
```
