## Appendix, as digital supplementary material to this article for "Worker-behavior and behavior-behavior interaction networks in the trap-jaw ant *Odontomachus chelifer* (Latreille, 1802) (Hymenoptera: Formicidae)": Script S7.R

```
#BTNs total 1 = colony a; total 2 = colony b; total 3 = colony c
library(tnet)
#BTNC1totaligs_b_tnet<- cbind(get.edgelist(BTNC1totaligs_b, names=TRUE), E(BTNC1totaligs_b)$weight)
BTNC1totaligs_b_tnet<- cbind(get.edgelist(BTNC1totaligs_b, names=FALSE), E(BTNC1totaligs_b)$weight)
# 1 = WL, 2 = SG, 3= G, 4= BG, 5= BC, 6=BC, 7=CB, 8=CD, 9=FP, 10=FD

betweenness_w(BTNC1totaligs_b_tnet, alpha=0.5)
#
BTN1centrality_wl<-14
BTN1centrality_sg<-36
BTN1centrality_g<-0
BTN1centrality_bg<-0
BTN1centrality_bc<-14
BTN1centrality_cb<-0
BTN1centrality_cd<-0
BTN1centrality_fp<-0
BTN1centrality_cf<-0
BTN1centrality_fd<-0

############################################
#BTNC2totaligs_b_tnet<- cbind(get.edgelist(BTNC2totaligs_b, names=TRUE), E(BTNC2totaligs_b)$weight)
BTNC2totaligs_b_tnet<- cbind(get.edgelist(BTNC2totaligs_b, names=FALSE), E(BTNC2totaligs_b)$weight)
#node.size<-setNames(c(0, 15, 0, 10, 6, 2, 5, 0),c("WL", "SG","AT", "G", "BG", "BC", "CB", "FP"))

betweenness_w(BTNC2totaligs_b_tnet, alpha=0.5)
#
BTN2centrality_wl<-0
BTN2centrality_sg<-17
BTN2centrality_at<-0
BTN2centrality_g<-8
BTN2centrality_bg<-5
BTN2centrality_bc<-2  
BTN2centrality_cb<-3   
BTN2centrality_fp<-0   

#BTNC3totaligs_b_tnet<- cbind(get.edgelist(BTNC3totaligs_b, names=TRUE), E(BTNC3totaligs_b)$weight)
BTNC3totaligs_b_tnet<- cbind(get.edgelist(BTNC3totaligs_b, names=FALSE), E(BTNC3totaligs_b)$weight)
#node.size<-setNames(c(0, 15, 0, 10, 6, 2, 5, 0),c("WL", "SG","AT", "G", "BG", "BC", "CB", "FP"))

betweenness_w(BTNC3totaligs_b_tnet, alpha=0.5)
#
BTN3centrality_wl<-5
BTN3centrality_sg<-0
BTN3centrality_at<-0
BTN3centrality_cf<-0
BTN3centrality_bc<-0 
BTN3centrality_fp<-5   
BTN3centrality_fd<-6 

#####################################################################################################
############ graphs to igraph 
conversion <- function(x){
  tnet_igraph(x,  type="weighted one-mode tnet", directed=TRUE)
}

#total1######################
Nperm=1000
null.model_w_n1 <- lapply(seq_len(Nperm), function(x){  
  randomnet <- rg_reshuffling_w(BTNC1totaligs_b_tnet, option="weights") 
})

null.model_w_n1_igraph<-lapply(null.model_w_n1, conversion)


Nperm=1000
null.model_l_n1 <- lapply(seq_len(Nperm), function(x){  
  randomnet <- rg_reshuffling_w(BTNC1totaligs_b_tnet, option="links") 
})

null.model_l_n1_igraph<-lapply(null.model_l_n1, conversion)
########################################################################
###############################################################################
#total 2
Nperm=1000
null.model_w_n2 <- lapply(seq_len(Nperm), function(x){  
  randomnet <- rg_reshuffling_w(BTNC2totaligs_b_tnet, option="weights") 
})

null.model_w_n2_igraph<-lapply(null.model_w_n2, conversion)

Nperm=1000
null.model_l_n2 <- lapply(seq_len(Nperm), function(x){  
  randomnet <- rg_reshuffling_w(BTNC2totaligs_b_tnet, option="links") 
})

null.model_l_n2_igraph<-lapply(null.model_l_n2, conversion)


##################################################################################
#total 3
Nperm=1000
null.model_w_n3 <- lapply(seq_len(Nperm), function(x){  
  randomnet <- rg_reshuffling_w(BTNC3totaligs_b_tnet, option="weights") 
})

null.model_w_n3_igraph<-lapply(null.model_w_n3, conversion)

Nperm=1000
null.model_l_n3 <- lapply(seq_len(Nperm), function(x){  
  randomnet <- rg_reshuffling_w(BTNC3totaligs_b_tnet, option="links") 
})

null.model_l_n3_igraph<-lapply(null.model_l_n3, conversion)


##################################################################################
```
