## Appendix, as digital supplementary material to this article for "Worker-behavior and behavior-behavior interaction networks in the trap-jaw ant *Odontomachus chelifer* (Latreille, 1802) (Hymenoptera: Formicidae)": Script S8.R

```
#################################################################################################
#BBNs of inactivity - colony a
#BTNC1Itotaligs_b_tnet<- cbind(get.edgelist(BTNC1Itotaligs_b, names=TRUE), E(BTNC1totaligs_b)$weight)
BTNC1Itotaligs_b_tnet<- cbind(get.edgelist(BTNC1Itotaligs_b, names=FALSE), E(BTNC1Itotaligs_b)$weight)

BTNC1Itotaligs_b_tnet_degreeout<-degree_w(BTNC1Itotaligs_b_tnet,measure=c("output"), type="out", alpha=0.5)

BTNC1Itotaligs_b_tnet_degreein<-degree_w(BTNC1Itotaligs_b_tnet,measure=c("output"), type="in", alpha=0.5)

BTNC1Itotaligs_b_tnet_degreeout
#
BTN1Icentrality_degreeout_wl<-13.5
BTN1Icentrality_degreeout_sg<-24.5
BTN1Icentrality_degreeout_at<-9.3
BTN1Icentrality_degreeout_g<-17.5
BTN1Icentrality_degreeout_bg<-11.7
BTN1Icentrality_degreeout_bc<-21.5
BTN1Icentrality_degreeout_cb<-14.1
BTN1Icentrality_degreeout_cd<-9.9
BTN1Icentrality_degreeout_fp<-10.3
BTN1Icentrality_degreeout_cf<-10.7
BTN1Icentrality_degreeout_fd<-22.6

BTNC1Itotaligs_b_tnet_degreein
#
BTN1Icentrality_degreein_wl<-37
BTN1Icentrality_degreein_sg<-36.3
BTN1Icentrality_degreein_at<-3.3
BTN1Icentrality_degreein_g<-18.2
BTN1Icentrality_degreein_bg<-12.6
BTN1Icentrality_degreein_bc<-18.1
BTN1Icentrality_degreein_cb<-11.7
BTN1Icentrality_degreein_cd<-4.3
BTN1Icentrality_degreein_fp<-4.5
BTN1Icentrality_degreein_cf<-2.8
BTN1Icentrality_degreein_fd<-17
########################################
#1 = WL, 2 = SG, , 3=AT, 4=G, 5= BG, 6= BC, 7= CB, 8= CD, 9=CF, 10=FD

##############################################################################################
# BBNs of inactivity - colony b

BTNC2Itotaligs_b_tnet<- cbind(get.edgelist(BTNC2Itotaligs_b, names=FALSE), E(BTNC2Itotaligs_b)$weight)
BTNC2Itotaligs_b_tnet_degreeout<-degree_w(BTNC2Itotaligs_b_tnet,measure=c("output"), type="out", alpha=0.5)
BTNC2Itotaligs_b_tnet_degreein<-degree_w(BTNC2Itotaligs_b_tnet,measure=c("output"), type="in", alpha=0.5)

#
BTNC2Itotaligs_b_tnet_degreeout
#
BTN2Icentrality_degreeout_wl<-6.5
BTN2Icentrality_degreeout_sg<-11.1
BTN2Icentrality_degreeout_at<-9.4
BTN2Icentrality_degreeout_g<-20.3
BTN2Icentrality_degreeout_bg<-17.5
BTN2Icentrality_degreeout_bc<-20
BTN2Icentrality_degreeout_cb<-17.8 
BTN2Icentrality_degreeout_cd<-4.7
BTN2Icentrality_degreeout_fp<-4.8 

#
BTNC2Itotaligs_b_tnet_degreein
#
BTN2Icentrality_degreein_wl<-13.5
BTN2Icentrality_degreein_sg<-27.2
BTN2Icentrality_degreein_at<-13.6
BTN2Icentrality_degreein_g<-17.6
BTN2Icentrality_degreein_bg<-10.3
BTN2Icentrality_degreein_bc<-17.3  
BTN2Icentrality_degreein_cb<-7.3
BTN2Icentrality_degreein_cd<-2.1
BTN2Icentrality_degreein_fp<-3.2 
####################

######################################################################################################
#BBNs of inactivity - colony c
BTNC3Itotaligs_b_tnet<- cbind(get.edgelist(BTNC3Itotaligs_b, names=FALSE), E(BTNC3Itotaligs_b)$weight)

BTNC3Igenigs_b_tnet<- cbind(get.edgelist(BTNC3Igenigs_b, names=FALSE), E(BTNC3Igenigs_b)$weight)

BTNC3Iespigs_b_tnet<- cbind(get.edgelist(BTNC3Iespigs_b, names=FALSE), E(BTNC3Iespigs_b)$weight)

BTNC3Itotaligs_b_tnet_degreeout<-degree_w(BTNC3Itotaligs_b_tnet,measure=c("degree","output"), type="out", alpha=0.5)
BTNC3Igenigs_b_tnet_degreeout<-degree_w(BTNC3Igenigs_b_tnet,measure=c("output"), type="out", alpha=0.5)
BTNC3Iespigs_b_tnet_degreeout<-degree_w(BTNC3Iespigs_b_tnet,measure=c("output"), type="out", alpha=0.5)

BTNC3Itotaligs_b_tnet_degreein<-degree_w(BTNC3Itotaligs_b_tnet,measure=c("output"), type="in", alpha=0.5)
BTNC3Igenigs_b_tnet_degreein<-degree_w(BTNC3Igenigs_b_tnet,measure=c("output"), type="in", alpha=0.5)
BTNC3Iespigs_b_tnet_degreein<-degree_w(BTNC3Iespigs_b_tnet,measure=c("output"), type="in", alpha=0.5)


BTNC3Itotaligs_b_tnet_degreeout
#
BTN3Icentrality_degreeout_wl<-6.5
BTN3Icentrality_degreeout_sg<-7
BTN3Icentrality_degreeout_at<-4.1
BTN3Icentrality_degreeout_cf<-6.9
BTN3Icentrality_degreeout_bc<-2.5
BTN3Icentrality_degreeout_fp<-11  
BTN3Icentrality_degreeout_fd<-12.6

BTNC3Itotaligs_b_tnet_degreein

BTN3Icentrality_degreein_wl<-11.9
BTN3Icentrality_degreein_sg<-8.9
BTN3Icentrality_degreein_at<-2.9
BTN3Icentrality_degreein_cf<-7.3
BTN3Icentrality_degreein_bc<-0
BTN3Icentrality_degreein_fp<-8.8   
BTN3Icentrality_degreein_fd<-10.8

###########################################################

###
net.zscore = function(obsval, nullval) {
  (obsval - mean(nullval))/sd(nullval)  
} 
```
