## Appendix, as digital supplementary material to this article for "Worker-behavior and behavior-behavior interaction networks in the trap-jaw ant *Odontomachus chelifer* (Latreille, 1802) (Hymenoptera: Formicidae)": Script S9.R

```
#TOTAL load the TNET data first
#betweness null model 1
# 1 = WL, 2 = SG, 3= G, 4= BG, 5= BC, 6=CB, 7=CD, 8=FP, 9=CF, 10=FD

net.zscore = function(obsval, nullval) {
  (obsval - mean(nullval))/sd(nullval)  
} 

#colony 1 = colony A  #weights null model 1
Nperm=1000
null.model_w_n1 <- lapply(seq_len(Nperm), function(x){  
  randomnet <- rg_reshuffling_w(BTNC1totaligs_b_tnet, option="weights") 
})

bet_weights <- function(x){
  betweenness_w(x, alpha=0.5)
}

bet_nullmodel_w_n1<-lapply(null.model_w_n1, bet_weights)

#
BTN1centrality_all<-betweenness_w(BTNC1totaligs_b_tnet, alpha=1)
BTN1centrality_all<-(BTN1centrality_all[,2])

bet_nullmodel_w_n1_ALL<-lapply(bet_nullmodel_w_n1, function(x) x[,2]) #all
bt.all_c1<-as.data.frame(unlist(bet_nullmodel_w_n1_ALL))
bt.all_c1<-as.numeric(unlist(bt.all_c1)) 

zscore<-net.zscore(BTN1centrality_all, bt.all_c1)
zscore
net.metric.pvalue <- 2*pnorm(-abs(zscore))
net.metric.pvalue # 

#colony a named as colony 1 
#####node bet #weights null model 1

bet_nullmodel_w_n1_WL<-lapply(bet_nullmodel_w_n1, function(x) x[1,2]) #WL

bt.wl_c1<-as.data.frame(unlist(bet_nullmodel_w_n1_WL))
bt.wl_c1<-as.numeric(unlist(bt.wl_c1)) #mean 0.31

#praw_wl.c1<-(sum(abs(bt.wl_c1) > abs(BTN1centrality_wl)) + 1) / (length(bt.wl_c1) + 1) #p 1.88
#praw_wl.c1*2

zscore<-net.zscore(BTN1centrality_wl, bt.wl_c1)
zscore
net.metric.pvalue <- 2*pnorm(-abs(zscore))
net.metric.pvalue # 0.06

#
bet_nullmodel_w_n1_SG<-lapply(bet_nullmodel_w_n1, function(x) x[2,2]) #SG

bt.sg_c1<-as.data.frame(unlist(bet_nullmodel_w_n1_SG))
bt.sg_c1<-as.numeric(unlist(bt.sg_c1)) #mean 0.31

praw_sg.c1<-(sum(abs(bt.sg_c1) > abs(BTN1centrality_sg)) + 1) / (length(bt.sg_c1) + 1) #p 0.003
praw_sg.c1*2

zscore<-net.zscore(BTN1centrality_sg, bt.sg_c1)
zscore
net.metric.pvalue <- 2*pnorm(-abs(zscore))
net.metric.pvalue # 0.06

#
bet_nullmodel_w_n1_G<-lapply(bet_nullmodel_w_n1, function(x) x[3,2]) #G

bt.g_c1<-as.data.frame(unlist(bet_nullmodel_w_n1_G))
bt.g_c1<-as.numeric(unlist(bt.g_c1)) #mean 

praw_g.c1<-(sum(abs(bt.g_c1) >= abs(BTN1centrality_g)) + 1) / (length(bt.g_c1) + 1) #p 1
praw_g.c1*2

zscore<-net.zscore(BTN1centrality_g, bt.g_c1)
zscore
net.metric.pvalue <- 2*pnorm(-abs(zscore))
net.metric.pvalue # 0.95

#
bet_nullmodel_w_n1_BG<-lapply(bet_nullmodel_w_n1, function(x) x[4,2]) #BG

bt.bg_c1<-as.data.frame(unlist(bet_nullmodel_w_n1_BG))
bt.bg_c1<-as.numeric(unlist(bt.bg_c1)) #mean 

praw_bg.c1<-(sum(abs(bt.bg_c1) >= abs(BTN1centrality_bg)) + 1) / (length(bt.bg_c1) + 1) #p 1
praw_bg.c1

zscore<-net.zscore(BTN1centrality_bg, bt.bg_c1)
zscore
net.metric.pvalue <- 2*pnorm(-abs(zscore))
net.metric.pvalue # 

#
bet_nullmodel_w_n1_BC<-lapply(bet_nullmodel_w_n1, function(x) x[5,2]) #BC

bt.bc_c1<-as.data.frame(unlist(bet_nullmodel_w_n1_BC))
bt.bc_c1<-as.numeric(unlist(bt.bc_c1)) #mean 

praw_bc.c1<-(sum(abs(bt.bc_c1) >= abs(BTN1centrality_bc)) + 1) / (length(bt.bc_c1) + 1) #p 1
praw_bc.c1

zscore<-net.zscore(BTN1centrality_bc, bt.bc_c1)
zscore
net.metric.pvalue <- 2*pnorm(-abs(zscore))
net.metric.pvalue # 

#
bet_nullmodel_w_n1_CB<-lapply(bet_nullmodel_w_n1, function(x) x[6,2]) #CB

bt.cb_c1<-as.data.frame(unlist(bet_nullmodel_w_n1_CB))
bt.cb_c1<-as.numeric(unlist(bt.cb_c1)) #mean 

praw_cb.c1<-(sum(abs(bt.cb_c1) >= abs(BTN1centrality_cb)) + 1) / (length(bt.cb_c1) + 1) #p
praw_cb.c1

zscore<-net.zscore(BTN1centrality_cb, bt.cb_c1)
zscore
net.metric.pvalue <- 2*pnorm(-abs(zscore))
net.metric.pvalue #


bet_nullmodel_w_n1_CD<-lapply(bet_nullmodel_w_n1, function(x) x[7,2]) #CD
bet_nullmodel_w_n1_FP<-lapply(bet_nullmodel_w_n1, function(x) x[8,2]) #FP
bet_nullmodel_w_n1_CF<-lapply(bet_nullmodel_w_n1, function(x) x[9,2]) #CF
bet_nullmodel_w_n1_FD<-lapply(bet_nullmodel_w_n1, function(x) x[10,2]) #FD

################################################################################################
#colony 1/ colony a
#####node bet #weights and links null model 2
# 1 = WL, 2 = SG, 3= G, 4= BG, 5= BC, 6=CB, 7=CD, 8=FP, 9=CF, 10=FD
Nperm=1000
null.model_l_n1 <- lapply(seq_len(Nperm), function(x){  
  randomnet <- rg_reshuffling_w(BTNC1totaligs_b_tnet, option="links") 
})

bet_links <- function(x){
  betweenness_w(x, alpha=0.5)
}

bet_nullmodel_l_n1<-lapply(null.model_l_n1, bet_links)

BTN1centrality_all<-betweenness_w(BTNC1totaligs_b_tnet, alpha=0.5)
BTN1centrality_all<-mean(BTN1centrality_all[,2])

bet_nullmodel_l_n1_ALL<-lapply(bet_nullmodel_l_n1, function(x) x[,2]) #all
bt.all_c1<-as.data.frame(unlist(bet_nullmodel_l_n1_ALL))
bt.all_c1<-as.numeric(unlist(bt.all_c1)) 

zscore<-net.zscore(BTN1centrality_all, bt.all_c1)
zscore
net.metric.pvalue <- 2*pnorm(-abs(zscore))
net.metric.pvalue # 

##############node bet #links null model 2

bet_nullmodel_l_n1_WL<-lapply(bet_nullmodel_l_n1, function(x) x[1,2]) #WL

bt.wl_c1<-as.data.frame(unlist(bet_nullmodel_l_n1_WL))
bt.wl_c1<-as.numeric(unlist(bt.wl_c1)) #

#praw_wl.c1<-(sum(abs(bt.wl_c1) >= abs(BTN1centrality_wl)) + 1) / (length(bt.wl_c1) + 1) #p 1
#praw_wl.c1

zscore<-net.zscore(BTN1centrality_wl, bt.wl_c1)
zscore
net.metric.pvalue <- 2*pnorm(-abs(zscore))
net.metric.pvalue # 0.13

#
bet_nullmodel_l_n1_SG<-lapply(bet_nullmodel_l_n1, function(x) x[2,2]) #SG

bt.sg_c1<-as.data.frame(unlist(bet_nullmodel_l_n1_SG))
bt.sg_c1<-as.numeric(unlist(bt.sg_c1)) #

#praw_sg.c1<-(sum(abs(bt.sg_c1) >= abs(BTN1centrality_sg)) + 1) / (length(bt.sg_c1) + 1) #p 0.23
#praw_sg.c1

zscore<-net.zscore(BTN1centrality_sg, bt.sg_c1)
zscore
net.metric.pvalue <- 2*pnorm(-abs(zscore))
net.metric.pvalue # 0.13

#
bet_nullmodel_l_n1_G<-lapply(bet_nullmodel_l_n1, function(x) x[3,2]) #G

bt.g_c1<-as.data.frame(unlist(bet_nullmodel_l_n1_G))
bt.g_c1<-as.numeric(unlist(bt.g_c1)) #mean 

praw_g.c1<-(sum(abs(bt.g_c1) >= abs(BTN1centrality_g)) + 1) / (length(bt.g_c1) + 1) #p 1 
praw_g.c1

zscore<-net.zscore(BTN1centrality_g, bt.g_c1)
zscore
net.metric.pvalue <- 2*pnorm(-abs(zscore))
net.metric.pvalue # 1

#
bet_nullmodel_l_n1_BG<-lapply(bet_nullmodel_l_n1, function(x) x[4,2]) #BG

bt.bg_c1<-as.data.frame(unlist(bet_nullmodel_l_n1_BG))
bt.bg_c1<-as.numeric(unlist(bt.bg_c1)) #mean 

praw_bg.c1<-(sum(abs(bt.bg_c1) >= abs(BTN1centrality_bg)) + 1) / (length(bt.bg_c1) + 1) #p 1
praw_bg.c1

zscore<-net.zscore(BTN1centrality_bg, bt.bg_c1)
zscore
net.metric.pvalue <- 2*pnorm(-abs(zscore))
net.metric.pvalue # 1

#
bet_nullmodel_l_n1_BC<-lapply(bet_nullmodel_l_n1, function(x) x[5,2]) #BC

bt.bc_c1<-as.data.frame(unlist(bet_nullmodel_l_n1_BC))
bt.bc_c1<-as.numeric(unlist(bt.bc_c1)) #mean 

praw_bc.c1<-(sum(abs(bt.bc_c1) >= abs(BTN1centrality_bc)) + 1) / (length(bt.bc_c1) + 1) #p 1
praw_bc.c1

zscore<-net.zscore(BTN1centrality_bc, bt.bc_c1)
zscore
net.metric.pvalue <- 2*pnorm(-abs(zscore))
net.metric.pvalue # 1

#
bet_nullmodel_w_n1_CB<-lapply(bet_nullmodel_w_n1, function(x) x[6,2]) #CB
bet_nullmodel_w_n1_CD<-lapply(bet_nullmodel_w_n1, function(x) x[7,2]) #CD
bet_nullmodel_w_n1_FP<-lapply(bet_nullmodel_w_n1, function(x) x[8,2]) #FP
bet_nullmodel_w_n1_CF<-lapply(bet_nullmodel_w_n1, function(x) x[9,2]) #CF
bet_nullmodel_w_n1_FD<-lapply(bet_nullmodel_w_n1, function(x) x[10,2]) #FD

####################################################################################
#colony 2 / colony b 
#####node bet #weights null model 1
#node.size<-setNames(c(0, 15, 0, 10, 6, 2, 5, 0),c("WL", "SG","AT", "G", "BG", "BC", "CB", "FP"))

Nperm=1000
null.model_w_n2 <- lapply(seq_len(Nperm), function(x){  
  randomnet <- rg_reshuffling_w(BTNC2totaligs_b_tnet, option="weights") 
})

bet_weights <- function(x){
  betweenness_w(x, alpha=0.5)
}

bet_nullmodel_w_n2<-lapply(null.model_w_n2, bet_weights)

#colony 2
#####node bet #weights null model 1

bet_nullmodel_w_n2_WL<-lapply(bet_nullmodel_w_n2, function(x) x[1,2]) #WL

bt.wl_c2<-as.data.frame(unlist(bet_nullmodel_w_n2_WL))
bt.wl_c2<-as.numeric(unlist(bt.wl_c2)) 

praw_wl.c2<-(sum(abs(bt.wl_c2) >= abs(BTN2centrality_wl)) + 1) / (length(bt.wl_c2) + 1) #p 1
praw_wl.c2

zscore<-net.zscore(BTN2centrality_wl, bt.wl_c2)
zscore
net.metric.pvalue <- 2*pnorm(-abs(zscore))
net.metric.pvalue # 0.35

#
bet_nullmodel_w_n2_SG<-lapply(bet_nullmodel_w_n2, function(x) x[2,2]) #SG

bt.sg_c2<-as.data.frame(unlist(bet_nullmodel_w_n2_SG))
bt.sg_c2<-as.numeric(unlist(bt.sg_c2)) 

praw_sg.c2<-(sum(abs(bt.sg_c2) >= abs(BTN2centrality_sg)) + 1) / (length(bt.sg_c2) + 1) #p 0.0009
praw_sg.c2

zscore<-net.zscore(BTN2centrality_sg, bt.sg_c2)
zscore
net.metric.pvalue <- 2*pnorm(-abs(zscore))
net.metric.pvalue # 0.43

#
bet_nullmodel_w_n2_AT<-lapply(bet_nullmodel_w_n2, function(x) x[3,2]) #AT

bt.at_c2<-as.data.frame(unlist(bet_nullmodel_w_n2_AT))
bt.at_c2<-as.numeric(unlist(bt.at_c2)) 

praw_at.c2<-(sum(abs(bt.at_c2) >= abs(BTN2centrality_at)) + 1) / (length(bt.at_c2) + 1) #p 1
praw_at.c2

zscore<-net.zscore(BTN2centrality_at, bt.at_c2)
zscore
net.metric.pvalue <- 2*pnorm(-abs(zscore))
net.metric.pvalue # 0.97

#
bet_nullmodel_w_n2_G<-lapply(bet_nullmodel_w_n2, function(x) x[4,2]) #G

bt.g_c2<-as.data.frame(unlist(bet_nullmodel_w_n2_G))
bt.g_c2<-as.numeric(unlist(bt.g_c2)) 

praw_g.c2<-(sum(abs(bt.g_c2) >= abs(BTN2centrality_g)) + 1) / (length(bt.g_c2) + 1) #p 
praw_g.c2

zscore<-net.zscore(BTN2centrality_g, bt.g_c2)
zscore
net.metric.pvalue <- 2*pnorm(-abs(zscore))
net.metric.pvalue # 0.38

#
bet_nullmodel_w_n2_BG<-lapply(bet_nullmodel_w_n2, function(x) x[5,2]) #BG

bt.bg_c2<-as.data.frame(unlist(bet_nullmodel_w_n2_BG))
bt.bg_c2<-as.numeric(unlist(bt.bg_c2)) 

praw_bg.c2<-(sum(abs(bt.bg_c2) >= abs(BTN2centrality_bg)) + 1) / (length(bt.bg_c2) + 1) #p 
praw_bg.c2

zscore<-net.zscore(BTN2centrality_bg, bt.bg_c2)
zscore
net.metric.pvalue <- 2*pnorm(-abs(zscore))
net.metric.pvalue # 0.95

#
bet_nullmodel_w_n2_BC<-lapply(bet_nullmodel_w_n2, function(x) x[6,2]) #BC

bt.bc_c2<-as.data.frame(unlist(bet_nullmodel_w_n2_BC))
bt.bc_c2<-as.numeric(unlist(bt.bc_c2)) 

zscore<-net.zscore(BTN2centrality_bc, bt.bc_c2)
zscore
net.metric.pvalue <- 2*pnorm(-abs(zscore))
net.metric.pvalue # 0.39

praw_bc.c2<-(sum(abs(bt.bc_c2) >= abs(BTN2centrality_bc)) + 1) / (length(bt.bc_c2) + 1) #p 
praw_bc.c2

#
bet_nullmodel_w_n2_CB<-lapply(bet_nullmodel_w_n2, function(x) x[7,2]) #CB

bt.cb_c2<-as.data.frame(unlist(bet_nullmodel_w_n2_CB))
bt.cb_c2<-as.numeric(unlist(bt.bc_c2)) 

praw_cb.c2<-(sum(abs(bt.cb_c2) >= abs(BTN2centrality_cb)) + 1) / (length(bt.cb_c2) + 1) #p 
praw_cb.c2

zscore<-net.zscore(BTN2centrality_cb, bt.cb_c2)
zscore
net.metric.pvalue <- 2*pnorm(-abs(zscore))
net.metric.pvalue #

#
bet_nullmodel_w_n2_FP<-lapply(bet_nullmodel_w_n2, function(x) x[8,2]) #FP

bt.fp_c2<-as.data.frame(unlist(bet_nullmodel_w_n2_FP))
bt.fp_c2<-as.numeric(unlist(bt.fp_c2)) 

praw_fp.c2<-(sum(abs(bt.fp_c2) >= abs(BTN2centrality_fp)) + 1) / (length(bt.fp_c2) + 1) #p 
praw_fp.c2

zscore<-net.zscore(BTN2centrality_fp, bt.fp_c2)
zscore
net.metric.pvalue <- 2*pnorm(-abs(zscore))
net.metric.pvalue #


################################################################################################
#colony 2 / colony b 
#####node bet #weights and links null model 2
#1 = "WL", 2= "SG", 3= "AT", 4= "G", 5="BG", 6="BC", 7="CB", 8="FP")

Nperm=1000
null.model_l_n2 <- lapply(seq_len(Nperm), function(x){  
  randomnet <- rg_reshuffling_w(BTNC2totaligs_b_tnet, option="links") 
})

bet_links <- function(x){
  betweenness_w(x, alpha=0.5)
}

bet_nullmodel_l_n2<-lapply(null.model_l_n2, bet_links)

##############node bet #links null model 2

bet_nullmodel_l_n2_WL<-lapply(bet_nullmodel_l_n2, function(x) x[1,2]) #WL

bt.wl_c2<-as.data.frame(unlist(bet_nullmodel_l_n2_WL))
bt.wl_c2<-as.numeric(unlist(bt.wl_c2)) #

praw_wl.c2<-(sum(abs(bt.wl_c2) >= abs(BTN2centrality_wl)) + 1) / (length(bt.wl_c2) + 1) #p 
praw_wl.c2

zscore<-net.zscore(BTN2centrality_wl, bt.wl_c2)
zscore
net.metric.pvalue <- 2*pnorm(-abs(zscore))
net.metric.pvalue #

#
bet_nullmodel_l_n2_SG<-lapply(bet_nullmodel_l_n2, function(x) x[2,2]) #SG

bt.sg_c2<-as.data.frame(unlist(bet_nullmodel_l_n2_SG))
bt.sg_c2<-as.numeric(unlist(bt.sg_c2)) #

praw_sg.c2<-(sum(abs(bt.wl_c2) >= abs(BTN2centrality_sg)) + 1) / (length(bt.sg_c2) + 1) #p 
praw_sg.c2

zscore<-net.zscore(BTN2centrality_sg, bt.sg_c2)
zscore
net.metric.pvalue <- 2*pnorm(-abs(zscore))
net.metric.pvalue #
#

bet_nullmodel_l_n2_AT<-lapply(bet_nullmodel_l_n2, function(x) x[3,2]) #AT

bt.at_c2<-as.data.frame(unlist(bet_nullmodel_l_n2_AT))
bt.at_c2<-as.numeric(unlist(bt.at_c2)) #

praw_at.c2<-(sum(abs(bt.at_c2) >= abs(BTN2centrality_at)) + 1) / (length(bt.at_c2) + 1) #p 
praw_at.c2

zscore<-net.zscore(BTN2centrality_at, bt.at_c2)
zscore
net.metric.pvalue <- 2*pnorm(-abs(zscore))
net.metric.pvalue #
#

bet_nullmodel_l_n2_G<-lapply(bet_nullmodel_l_n2, function(x) x[4,2]) #G

bt.g_c2<-as.data.frame(unlist(bet_nullmodel_l_n2_G))
bt.g_c2<-as.numeric(unlist(bt.g_c2)) #

praw_g.c2<-(sum(abs(bt.g_c2) >= abs(BTN2centrality_g)) + 1) / (length(bt.g_c2) + 1) #p 
praw_g.c2

zscore<-net.zscore(BTN2centrality_g, bt.g_c2)
zscore
net.metric.pvalue <- 2*pnorm(-abs(zscore))
net.metric.pvalue # 0.95

#
bet_nullmodel_l_n2_BG<-lapply(bet_nullmodel_l_n2, function(x) x[5,2]) #BG

bt.bg_c2<-as.data.frame(unlist(bet_nullmodel_l_n2_BG))
bt.bg_c2<-as.numeric(unlist(bt.bg_c2)) #

#praw_bg.c2<-(sum(abs(bt.bg_c2) >= abs(BTN2centrality_bg)) + 1) / (length(bt.bg_c2) + 1) #p 
#praw_bg.c2

zscore<-net.zscore(BTN2centrality_bg, bt.bg_c2)
zscore
net.metric.pvalue <- 2*pnorm(-abs(zscore))
net.metric.pvalue # 0.88

#
bet_nullmodel_l_n2_BC<-lapply(bet_nullmodel_l_n2, function(x) x[6,2]) #BC

bt.bc_c2<-as.data.frame(unlist(bet_nullmodel_l_n2_BC))
bt.bc_c2<-as.numeric(unlist(bt.bc_c2)) #

#praw_bc.c2<-(sum(abs(bt.bc_c2) >= abs(BTN2centrality_bc)) + 1) / (length(bt.bc_c2) + 1) #p 
#praw_bc.c2

zscore<-net.zscore(BTN2centrality_bc, bt.bc_c2)
zscore
net.metric.pvalue <- 2*pnorm(-abs(zscore))
net.metric.pvalue # 

#
bet_nullmodel_l_n2_BC<-lapply(bet_nullmodel_l_n2, function(x) x[7,2]) #CB

bt.cb_c2<-as.data.frame(unlist(bet_nullmodel_l_n2_CB))
bt.cb_c2<-as.numeric(unlist(bt.cb_c2)) #

#praw_cb.c2<-(sum(abs(bt.cb_c2) >= abs(BTN2centrality_cb)) + 1) / (length(bt.cb_c2) + 1) #p 
#praw_cb.c2
#

zscore<-net.zscore(BTN2centrality_cb, bt.cb_c2)
zscore
net.metric.pvalue <- 2*pnorm(-abs(zscore))
net.metric.pvalue # 

bet_nullmodel_l_n2_FP<-lapply(bet_nullmodel_l_n2, function(x) x[8,2]) #FP

bt.fp_c2<-as.data.frame(unlist(bet_nullmodel_l_n2_FP))
bt.fp_c2<-as.numeric(unlist(bt.fp_c2)) #

#praw_fp.c2<-(sum(abs(bt.fp_c2) >= abs(BTN2centrality_fp)) + 1) / (length(bt.fp_c2) + 1) #p 
#praw_fp.c2

zscore<-net.zscore(BTN2centrality_fp, bt.fp_c2)
zscore
net.metric.pvalue <- 2*pnorm(-abs(zscore))
net.metric.pvalue # 

####################################################################################
#colony 3 / colony c 
#####node bet #weights null model 1
#1 = "WL", 2= "SG", 3= "AT", 4= "CF", 5="BC", 6="FP", 7="FD"

Nperm=1000
null.model_w_n3 <- lapply(seq_len(Nperm), function(x){  
  randomnet <- rg_reshuffling_w(BTNC3totaligs_b_tnet, option="weights") 
})

bet_weights <- function(x){
  betweenness_w(x, alpha=0.5)
}

bet_nullmodel_w_n3<-lapply(null.model_w_n3, bet_weights)

##############node bet #WEIGHTS null model 1

bet_nullmodel_w_n3_WL<-lapply(bet_nullmodel_w_n3, function(x) x[1,2]) #WL

bt.wl_c3<-as.data.frame(unlist(bet_nullmodel_w_n3_WL))
bt.wl_c3<-as.numeric(unlist(bt.wl_c3)) #

#praw_wl.c3<-(sum(abs(bt.wl_c3) >= abs(BTN3centrality_wl)) + 1) / (length(bt.wl_c3) + 1) #p 
#praw_wl.c3

zscore<-net.zscore(BTN3centrality_wl, bt.wl_c3)
zscore
net.metric.pvalue <- 2*pnorm(-abs(zscore))
net.metric.pvalue # 

#
bet_nullmodel_w_n3_SG<-lapply(bet_nullmodel_w_n3, function(x) x[2,2]) #SG

bt.sg_c3<-as.data.frame(unlist(bet_nullmodel_w_n3_SG))
bt.sg_c3<-as.numeric(unlist(bt.sg_c3)) #

#praw_sg.c3<-(sum(abs(bt.wl_c3) >= abs(BTN3centrality_sg)) + 1) / (length(bt.sg_c3) + 1) #p 
#praw_sg.c3

zscore<-net.zscore(BTN3centrality_sg, bt.sg_c3)
zscore
net.metric.pvalue <- 2*pnorm(-abs(zscore))
net.metric.pvalue # 

#
bet_nullmodel_w_n3_AT<-lapply(bet_nullmodel_w_n3, function(x) x[3,2]) #AT

bt.at_c3<-as.data.frame(unlist(bet_nullmodel_w_n3_AT))
bt.at_c3<-as.numeric(unlist(bt.at_c2)) #

#praw_at.c3<-(sum(abs(bt.at_c3) >= abs(BTN3centrality_at)) + 1) / (length(bt.at_c3) + 1) #p 
#praw_at.c3

zscore<-net.zscore(BTN3centrality_at, bt.at_c3)
zscore
net.metric.pvalue <- 2*pnorm(-abs(zscore))
net.metric.pvalue #

bet_nullmodel_w_n3_CF<-lapply(bet_nullmodel_w_n3, function(x) x[4,2]) #CF

bt.cf_c3<-as.data.frame(unlist(bet_nullmodel_w_n3_CF))
bt.cf_c3<-as.numeric(unlist(bt.cf_c3)) #

praw_cf.c3<-(sum(abs(bt.cf_c3) >= abs(BTN3centrality_cf)) + 1) / (length(bt.cf_c3) + 1) #p 
praw_cf.c3
#

bet_nullmodel_w_n3_BC<-lapply(bet_nullmodel_w_n3, function(x) x[5,2]) #BC

bt.bc_c3<-as.data.frame(unlist(bet_nullmodel_w_n3_CF))
bt.bc_c3<-as.numeric(unlist(bt.bc_c3)) #

praw_bc.c3<-(sum(abs(bt.bc_c3) >= abs(BTN3centrality_bc)) + 1) / (length(bt.bc_c3) + 1) #p 
praw_bc.c3
#

bet_nullmodel_w_n3_FP<-lapply(bet_nullmodel_w_n3, function(x) x[6,2]) #FP

bt.fp_c3<-as.data.frame(unlist(bet_nullmodel_w_n3_FP))
bt.fp_c3<-as.numeric(unlist(bt.fp_c3)) #

#praw_fp.c3<-(sum(abs(bt.fp_c3) >= abs(BTN3centrality_fp)) + 1) / (length(bt.fp_c3) + 1) #p 
#praw_fp.c3

zscore<-net.zscore(BTN3centrality_fp, bt.fp_c3)
zscore
net.metric.pvalue <- 2*pnorm(-abs(zscore))
net.metric.pvalue #

#
bet_nullmodel_w_n3_FD<-lapply(bet_nullmodel_w_n3, function(x) x[7,2]) #FD

bt.fd_c3<-as.data.frame(unlist(bet_nullmodel_w_n3_FD))
bt.fd_c3<-as.numeric(unlist(bt.fd_c3)) #

#praw_fd.c3<-(sum(abs(bt.fd_c3) >= abs(BTN3centrality_fd)) + 1) / (length(bt.fd_c3) + 1) #p 
#praw_fd.c3

zscore<-net.zscore(BTN3centrality_fd, bt.fd_c3)
zscore
net.metric.pvalue <- 2*pnorm(-abs(zscore))
net.metric.pvalue #
#

################################################################################################
#colony 3
#####node bet #weights and links null model 2
#1 = "WL", 2= "SG", 3= "AT", 4= "CF", 5="BC", 6="FP", 7="FD"

Nperm=1000
null.model_l_n3 <- lapply(seq_len(Nperm), function(x){  
  randomnet <- rg_reshuffling_w(BTNC3totaligs_b_tnet, option="links") 
})

bet_links <- function(x){
  betweenness_w(x, alpha=0.5)
}

bet_nullmodel_l_n3<-lapply(null.model_l_n3, bet_links)

##############node bet #links null model 2

bet_nullmodel_l_n3_WL<-lapply(bet_nullmodel_l_n3, function(x) x[1,2]) #WL

bt.wl_c3<-as.data.frame(unlist(bet_nullmodel_l_n3_WL))
bt.wl_c3<-as.numeric(unlist(bt.wl_c3)) #

#praw_wl.c3<-(sum(abs(bt.wl_c3) >= abs(BTN3centrality_wl)) + 1) / (length(bt.wl_c3) + 1) #p 
#praw_wl.c3

zscore<-net.zscore(BTN3centrality_wl, bt.wl_c3)
zscore
net.metric.pvalue <- 2*pnorm(-abs(zscore))
net.metric.pvalue #
#

bet_nullmodel_l_n3_SG<-lapply(bet_nullmodel_l_n3, function(x) x[2,2]) #SG

bt.sg_c3<-as.data.frame(unlist(bet_nullmodel_l_n3_SG))
bt.sg_c3<-as.numeric(unlist(bt.sg_c3)) #

zscore<-net.zscore(BTN3centrality_sg, bt.sg_c3)
zscore
net.metric.pvalue <- 2*pnorm(-abs(zscore))
net.metric.pvalue #
#
#

bet_nullmodel_l_n3_AT<-lapply(bet_nullmodel_l_n3, function(x) x[3,2]) #AT

bt.at_c3<-as.data.frame(unlist(bet_nullmodel_l_n3_AT))
bt.at_c3<-as.numeric(unlist(bt.at_c3)) #

zscore<-net.zscore(BTN3centrality_at, bt.at_c3)
zscore
net.metric.pvalue <- 2*pnorm(-abs(zscore))
net.metric.pvalue #
#

bet_nullmodel_l_n3_CF<-lapply(bet_nullmodel_l_n3, function(x) x[4,2]) #CF

bt.cf_c3<-as.data.frame(unlist(bet_nullmodel_l_n3_CF))
bt.cf_c3<-as.numeric(unlist(bt.cf_c3)) #

praw_cf.c3<-(sum(abs(bt.cf_c3) >= abs(BTN3centrality_cf)) + 1) / (length(bt.cf_c3) + 1) #p 
praw_cf.c3
#

bet_nullmodel_l_n3_BC<-lapply(bet_nullmodel_l_n3, function(x) x[5,2]) #BC

bt.bc_c3<-as.data.frame(unlist(bet_nullmodel_l_n3_BC))
bt.bc_c3<-as.numeric(unlist(bt.bc_c3)) #

praw_bc.c3<-(sum(abs(bt.bc_c3) >= abs(BTN3centrality_bc)) + 1) / (length(bt.bc_c3) + 1) #p 
praw_bc.c3
#

bet_nullmodel_l_n3_FP<-lapply(bet_nullmodel_l_n3, function(x) x[6,2]) #FP

bt.fp_c3<-as.data.frame(unlist(bet_nullmodel_l_n3_FP))
bt.fp_c3<-as.numeric(unlist(bt.fp_c3)) #

zscore<-net.zscore(BTN3centrality_fp, bt.fp_c3)
zscore
net.metric.pvalue <- 2*pnorm(-abs(zscore))
net.metric.pvalue #
#

bet_nullmodel_l_n3_FD<-lapply(bet_nullmodel_l_n3, function(x) x[7,2]) #FD

bt.fd_c3<-as.data.frame(unlist(bet_nullmodel_l_n3_FD))
bt.fd_c3<-as.numeric(unlist(bt.fd_c3)) #


zscore<-net.zscore(BTN3centrality_fd, bt.fd_c3)
zscore
net.metric.pvalue <- 2*pnorm(-abs(zscore))
net.metric.pvalue #
#


#
```
