## Appendix, as digital supplementary material to this article for "Worker-behavior and behavior-behavior interaction networks in the trap-jaw ant *Odontomachus chelifer* (Latreille, 1802) (Hymenoptera: Formicidae)": Script S10.R

```
library(RVAideMemoire)
library(corrplot)
library(BBmisc)
library(reshape2)
library(ggplot2)
#load data - BTNs including all the behaviors (even when a colony dont have one of them)
BTN_a<-read.table("Data S11.txt", header=TRUE)
BTN_b<-read.table("Data S13.txt", header=TRUE)
BTN_c<-read.table("Data S15.txt", header=TRUE)
#join them 
BTNall<-as.data.frame(BTN_a+BTN_b+BTN_c)

#extract, and isolate the diag
all_diagonal<-diag(as.matrix(BTNall))
no_diag<-colSums(BTNall, na.rm=TRUE)
diag(BTNall)=NA

behavioral_switch<-cbind(all_diagonal, no_diag)
behavioral_switch<-as.data.frame(behavioral_switch)
behavioral_switch_plot<-melt(behavioral_switch)

Font_My_Theme = theme(
  axis.title.x = element_text(size = 10),
  axis.title.y = element_text(size = 10),
  axis.text.x = element_text(size = 10),
  axis.text.y = element_text(size = 10))

## first part of the figure 6 
ggplot(behavioral_switch_plot,aes(x = reorder(Var1,-value), y = value, fill = Var2, ymax=0.40, ymin=0)) +
  geom_bar(stat="identity", position = "dodge", width = 0.7) +
  geom_bar(stat="identity", position = "dodge", colour = "black",
           width = 0.7) + theme_classic() + Font_My_Theme + scale_fill_grey()

#the non diagonal behaviors (task-switching) are significant higher than the diagonal ones (self-interaction)  
G.test(as.matrix(behavioral_switch+1)) 
#if we eliminate the behaviors with small frequencies the significance stands
G.test(as.matrix(behavioral_switch[-c(3, 8, 10), ]))

#stats for the switching frequency versus an uniform distribution of the data. 
G.theo.multcomp(na.omit(BTNall$wl+1), p.method = "BH")
G.theo.multcomp(na.omit(BTNall$sg+1), p.method = "BH")
G.theo.multcomp(na.omit(BTNall$at+1), p.method = "BH")
G.theo.multcomp(na.omit(BTNall$g+1), p.method = "BH")
G.theo.multcomp(na.omit(BTNall$bg+1), p.method = "BH")
G.theo.multcomp(na.omit(BTNall$bc+1), p.method = "BH")
G.theo.multcomp(na.omit(BTNall$cb+1), p.method = "BH")
G.theo.multcomp(na.omit(BTNall$cd+1), p.method = "BH")
G.theo.multcomp(na.omit(BTNall$fd+1), p.method = "BH")

BTNall_p_values<-read.table("Script S9.txt", header=TRUE) #p-values from the G-tests #Data  S22
# second part of the figure 6 
corrplot(as.matrix(BTNall), is.corr = FALSE, p.mat = as.matrix(BTNall_p_values), sig.level = .05, diag = F, col=gray.colors(100, rev = TRUE))
```
