## Appendix, as digital supplementary material to this article for "Worker-behavior and behavior-behavior interaction networks in the trap-jaw ant *Odontomachus chelifer* (Latreille, 1802) (Hymenoptera: Formicidae)": Script S13.R

```
#colony 3 #weights null model 1 out-degree
Nperm=1000
null.model_w_out_n3 <- lapply(seq_len(Nperm), function(x){  
  randomnet <- rg_reshuffling_w(BTNC3Itotaligs_b_tnet, option="weights") 
})

deg_out_weights <- function(x){
  degree_w(x,measure=c("output"), type="out", alpha=0.5)
}

deg_out_nullmodel_w_n3<-lapply(null.model_w_out_n3, deg_out_weights)

#colony 3
#####node degree out #weights null model 1

deg_out_nullmodel_w_n3_WL<-lapply(deg_out_nullmodel_w_n3, function(x) x[1,2]) #WL

deg_out.wl_c3<-as.data.frame(unlist(deg_out_nullmodel_w_n3_WL))
deg_out.wl_c3<-as.numeric(unlist(deg_out.wl_c3)) #

#praw_wl.c1<-(sum(abs(deg_out.wl_c1) >= abs(BTN1centrality_degreeout_wl)) + 1) / (length(deg_out.wl_c1) + 1) #p 1
#praw_wl.c1

zscore<-net.zscore(BTN3Icentrality_degreeout_wl, deg_out.wl_c3)
zscore
net.metric.pvalue <- 2*pnorm(-abs(zscore))
net.metric.pvalue # 
#

deg_out_nullmodel_w_n3_SG<-lapply(deg_out_nullmodel_w_n3, function(x) x[2,2]) #SG

deg_out.sg_c3<-as.data.frame(unlist(deg_out_nullmodel_w_n3_SG))
deg_out.sg_c3<-as.numeric(unlist(deg_out.sg_c3)) 

#praw_sg.c1<-(sum(abs(deg_out.sg_c1) >= abs(BTN1centrality_degreeout_sg)) + 1) / (length(deg_out.sg_c1) + 1) #p 0.99
#praw_sg.c1

zscore<-net.zscore(BTN3Icentrality_degreeout_sg, deg_out.sg_c3)
zscore
net.metric.pvalue <- 2*pnorm(-abs(zscore))
net.metric.pvalue # 

#
deg_out_nullmodel_w_n3_AT<-lapply(deg_out_nullmodel_w_n3, function(x) x[3,2]) #AT

deg_out.at_c3<-as.data.frame(unlist(deg_out_nullmodel_w_n3_AT))
deg_out.at_c3<-as.numeric(unlist(deg_out.at_c3)) #

#praw_at.c1<-(sum(abs(deg_out.at_c1) >= abs(BTN1centrality_degreeout_at)) + 1) / (length(deg_out.at_c1) + 1) #p  0.64
#praw_at.c1

zscore<-net.zscore(BTN3Icentrality_degreeout_at, deg_out.at_c3)
zscore
net.metric.pvalue <- 2*pnorm(-abs(zscore))
net.metric.pvalue # 
#

deg_out_nullmodel_w_n3_CF<-lapply(deg_out_nullmodel_w_n3, function(x) x[4,2]) #CF

deg_out.cf_c3<-as.data.frame(unlist(deg_out_nullmodel_w_n3_CF))
deg_out.cf_c3<-as.numeric(unlist(deg_out.cf_c3)) #

#praw_g.c1<-(sum(abs(deg_out.g_c1) >= abs(BTN1centrality_degreeout_g)) + 1) / (length(deg_out.g_c1) + 1) #p  0.46
#praw_g.c1

zscore<-net.zscore(BTN3Icentrality_degreeout_cf, deg_out.cf_c3)
zscore
net.metric.pvalue <- 2*pnorm(-abs(zscore))
net.metric.pvalue # 
#

deg_out_nullmodel_w_n3_BC<-lapply(deg_out_nullmodel_w_n3, function(x) x[5,2]) #BC

deg_out.bc_c3<-as.data.frame(unlist(deg_out_nullmodel_w_n3_BC))
deg_out.bc_c3<-as.numeric(unlist(deg_out.bc_c3)) #

zscore<-net.zscore(BTN3Icentrality_degreeout_bc, deg_out.bc_c3)
zscore
net.metric.pvalue <- 2*pnorm(-abs(zscore))
net.metric.pvalue # 
#

deg_out_nullmodel_w_n3_FP<-lapply(deg_out_nullmodel_w_n3, function(x) x[6,2]) #FP

deg_out.fp_c3<-as.data.frame(unlist(deg_out_nullmodel_w_n3_FP))
deg_out.fp_c3<-as.numeric(unlist(deg_out.fp_c3)) #

#praw_fp.c1<-(sum(abs(deg_out.fp_c1) >= abs(BTN1centrality_degreeout_fp)) + 1) / (length(deg_out.fp_c1) + 1) #p  0.058
#praw_fp.c1
#

zscore<-net.zscore(BTN3Icentrality_degreeout_fp, deg_out.fp_c3)
zscore
net.metric.pvalue <- 2*pnorm(-abs(zscore))
net.metric.pvalue # 
#

deg_out_nullmodel_w_n3_FD<-lapply(deg_out_nullmodel_w_n3, function(x) x[6,2]) #FD

deg_out.fd_c3<-as.data.frame(unlist(deg_out_nullmodel_w_n3_FD))
deg_out.fd_c3<-as.numeric(unlist(deg_out.fp_c3)) #

#praw_fp.c1<-(sum(abs(deg_out.fp_c1) >= abs(BTN1centrality_degreeout_fp)) + 1) / (length(deg_out.fp_c1) + 1) #p  0.058
#praw_fp.c1
#

zscore<-net.zscore(BTN3Icentrality_degreeout_fd, deg_out.fp_c3)
zscore
net.metric.pvalue <- 2*pnorm(-abs(zscore))
net.metric.pvalue # 
#
######################################################################################

#colony 3 #weights null model 1 in-degree
Nperm=1000
null.model_w_in_n3 <- lapply(seq_len(Nperm), function(x){  
  randomnet <- rg_reshuffling_w(BTNC3Itotaligs_b_tnet, option="weights") 
})

deg_in_weights <- function(x){
  degree_w(x,measure=c("output"), type="in", alpha=0.5)
}

deg_in_nullmodel_w_n3<-lapply(null.model_w_in_n3, deg_in_weights)

#####node degree in #weights null model 1

deg_in_nullmodel_w_n3_WL<-lapply(deg_in_nullmodel_w_n3, function(x) x[1,2]) #WL

deg_in.wl_c3<-as.data.frame(unlist(deg_in_nullmodel_w_n3_WL))
deg_in.wl_c3<-as.numeric(unlist(deg_in.wl_c3)) #

#praw_wl.c1<-(sum(abs(deg_in.wl_c1) >= abs(BTN1centrality_degreein_wl)) + 1) / (length(deg_in.wl_c1) + 1) #p 0.06
#praw_wl.c1

zscore<-net.zscore(BTN3Icentrality_degreein_wl, deg_in.wl_c3)
zscore
net.metric.pvalue <- 2*pnorm(-abs(zscore))
net.metric.pvalue # 

#
deg_in_nullmodel_w_n3_SG<-lapply(deg_in_nullmodel_w_n3, function(x) x[2,2]) #SG

deg_in.sg_c3<-as.data.frame(unlist(deg_in_nullmodel_w_n3_SG))
deg_in.sg_c3<-as.numeric(unlist(deg_in.sg_c3)) #

#praw_sg.c1<-(sum(abs(deg_in.sg_c1) >= abs(BTN1centrality_degreein_sg)) + 1) / (length(deg_in.sg_c1) + 1) #p 0.08
#praw_sg.c1

zscore<-net.zscore(BTN3Icentrality_degreein_sg, deg_in.sg_c3)
zscore
net.metric.pvalue <- 2*pnorm(-abs(zscore))
net.metric.pvalue # 

#
deg_in_nullmodel_w_n3_AT<-lapply(deg_in_nullmodel_w_n3, function(x) x[3,2]) #AT

deg_in.at_c3<-as.data.frame(unlist(deg_in_nullmodel_w_n3_AT))
deg_in.at_c3<-as.numeric(unlist(deg_in.at_c3)) #

#praw_at.c1<-(sum(abs(deg_in.at_c1) >= abs(BTN1centrality_degreein_at)) + 1) / (length(deg_in.at_c1) + 1) #p 0.96
#praw_at.c1

zscore<-net.zscore(BTN3Icentrality_degreein_at, deg_in.at_c3)
zscore
net.metric.pvalue <- 2*pnorm(-abs(zscore))
net.metric.pvalue # 

#
deg_in_nullmodel_w_n3_CF<-lapply(deg_in_nullmodel_w_n3, function(x) x[4,2]) #CF

deg_in.cf_c3<-as.data.frame(unlist(deg_in_nullmodel_w_n3_CF))
deg_in.cf_c3<-as.numeric(unlist(deg_in.cf_c3)) #

zscore<-net.zscore(BTN3Icentrality_degreein_cf, deg_in.cf_c3)
zscore
net.metric.pvalue <- 2*pnorm(-abs(zscore))
net.metric.pvalue # 

#
deg_in_nullmodel_w_n3_BC<-lapply(deg_in_nullmodel_w_n3, function(x) x[5,2]) #BC

deg_in.bc_c3<-as.data.frame(unlist(deg_in_nullmodel_w_n3_BC))
deg_in.bc_c3<-as.numeric(unlist(deg_in.bc_c3)) #


zscore<-net.zscore(BTN3Icentrality_degreein_bc, deg_in.bc_c3)
zscore
net.metric.pvalue <- 2*pnorm(-abs(zscore))
net.metric.pvalue # 

#
deg_in_nullmodel_w_n3_FP<-lapply(deg_in_nullmodel_w_n3, function(x) x[6,2]) #FP

deg_in.fp_c3<-as.data.frame(unlist(deg_in_nullmodel_w_n3_FP))
deg_in.fp_c3<-as.numeric(unlist(deg_in.fp_c3)) #

#praw_fp.c1<-(sum(abs(deg_in.fp_c1) >= abs(BTN1centrality_degreein_fp)) + 1) / (length(deg_in.fp_c1) + 1) #p 0.88
#praw_fp.c1

zscore<-net.zscore(BTN3Icentrality_degreein_fp, deg_in.fp_c3)
zscore
net.metric.pvalue <- 2*pnorm(-abs(zscore))
net.metric.pvalue #

#
deg_in_nullmodel_w_n3_FD<-lapply(deg_in_nullmodel_w_n3, function(x) x[7,2]) #FD

deg_in.fd_c3<-as.data.frame(unlist(deg_in_nullmodel_w_n3_FD))
deg_in.fd_c3<-as.numeric(unlist(deg_in.fd_c3)) #

#praw_fp.c1<-(sum(abs(deg_in.fp_c1) >= abs(BTN1centrality_degreein_fp)) + 1) / (length(deg_in.fp_c1) + 1) #p 0.88
#praw_fp.c1

zscore<-net.zscore(BTN3Icentrality_degreein_fd, deg_in.fd_c3)
zscore
net.metric.pvalue <- 2*pnorm(-abs(zscore))
net.metric.pvalue #

##############################################################################################


#colony 3 #weights null model 2 out-degree
Nperm=1000
null.model_l_out_n3 <- lapply(seq_len(Nperm), function(x){  
  randomnet <- rg_reshuffling_w(BTNC3Itotaligs_b_tnet, option="links") 
})

deg_out_links <- function(x){
  degree_w(x,measure=c("output"), type="out", alpha=0.5)
}

deg_out_nullmodel_l_n3<-lapply(null.model_l_out_n3, deg_out_links)

#colony 3
#####node degree out #weights null model 1

deg_out_nullmodel_l_n3_WL<-lapply(deg_out_nullmodel_l_n3, function(x) x[1,2]) #WL

deg_out.wl_c3<-as.data.frame(unlist(deg_out_nullmodel_l_n3_WL))
deg_out.wl_c3<-as.numeric(unlist(deg_out.wl_c3)) #

#praw_wl.c1<-(sum(abs(deg_out.wl_c1) >= abs(BTN1centrality_degreeout_wl)) + 1) / (length(deg_out.wl_c1) + 1) #p 1
#praw_wl.c1

zscore<-net.zscore(BTN3Icentrality_degreeout_wl, deg_out.wl_c3)
zscore
net.metric.pvalue <- 2*pnorm(-abs(zscore))
net.metric.pvalue # 
#

deg_out_nullmodel_l_n3_SG<-lapply(deg_out_nullmodel_l_n3, function(x) x[2,2]) #SG

deg_out.sg_c3<-as.data.frame(unlist(deg_out_nullmodel_l_n3_SG))
deg_out.sg_c3<-as.numeric(unlist(deg_out.sg_c3)) 

#praw_sg.c1<-(sum(abs(deg_out.sg_c1) >= abs(BTN1centrality_degreeout_sg)) + 1) / (length(deg_out.sg_c1) + 1) #p 0.99
#praw_sg.c1

zscore<-net.zscore(BTN3Icentrality_degreeout_sg, deg_out.sg_c3)
zscore
net.metric.pvalue <- 2*pnorm(-abs(zscore))
net.metric.pvalue # 

#
deg_out_nullmodel_l_n3_AT<-lapply(deg_out_nullmodel_l_n3, function(x) x[3,2]) #AT

deg_out.at_c3<-as.data.frame(unlist(deg_out_nullmodel_l_n3_AT))
deg_out.at_c3<-as.numeric(unlist(deg_out.at_c3)) #

#praw_at.c1<-(sum(abs(deg_out.at_c1) >= abs(BTN1centrality_degreeout_at)) + 1) / (length(deg_out.at_c1) + 1) #p  0.64
#praw_at.c1

zscore<-net.zscore(BTN2Icentrality_degreeout_at, deg_out.at_c3)
zscore
net.metric.pvalue <- 2*pnorm(-abs(zscore))
net.metric.pvalue # 
#

deg_out_nullmodel_l_n3_CF<-lapply(deg_out_nullmodel_l_n3, function(x) x[4,2]) #CF

deg_out.cf_c3<-as.data.frame(unlist(deg_out_nullmodel_l_n3_CF))
deg_out.cf_c3<-as.numeric(unlist(deg_out.cf_c3)) #

#praw_g.c1<-(sum(abs(deg_out.g_c1) >= abs(BTN1centrality_degreeout_g)) + 1) / (length(deg_out.g_c1) + 1) #p  0.46
#praw_g.c1

zscore<-net.zscore(BTN3Icentrality_degreeout_cf, deg_out.cf_c3)
zscore
net.metric.pvalue <- 2*pnorm(-abs(zscore))
net.metric.pvalue # 
#

deg_out_nullmodel_l_n3_BC<-lapply(deg_out_nullmodel_l_n3, function(x) x[5,2]) #BC

deg_out.bc_c3<-as.data.frame(unlist(deg_out_nullmodel_l_n3_BC))
deg_out.bc_c3<-as.numeric(unlist(deg_out.bc_c3)) #

#praw_bc.c1<-(sum(abs(deg_out.bc_c1) >= abs(BTN1centrality_degreeout_bc)) + 1) / (length(deg_out.bc_c1) + 1) #p  0.08
#praw_bc.c1

zscore<-net.zscore(BTN3Icentrality_degreeout_bc, deg_out.bc_c3)
zscore
net.metric.pvalue <- 2*pnorm(-abs(zscore))
net.metric.pvalue # 
#

deg_out_nullmodel_l_n3_FP<-lapply(deg_out_nullmodel_l_n3, function(x) x[6,2]) #FP

deg_out.fp_c3<-as.data.frame(unlist(deg_out_nullmodel_l_n3_FP))
deg_out.fp_c3<-as.numeric(unlist(deg_out.fp_c3)) #

#praw_fp.c1<-(sum(abs(deg_out.fp_c1) >= abs(BTN1centrality_degreeout_fp)) + 1) / (length(deg_out.fp_c1) + 1) #p  0.058
#praw_fp.c1
#

zscore<-net.zscore(BTN3Icentrality_degreeout_fp, deg_out.fp_c3)
zscore
net.metric.pvalue <- 2*pnorm(-abs(zscore))
net.metric.pvalue # 
#

deg_out_nullmodel_l_n3_FD<-lapply(deg_out_nullmodel_l_n3, function(x) x[6,2]) #FD

deg_out.fd_c3<-as.data.frame(unlist(deg_out_nullmodel_l_n3_FD))
deg_out.fd_c3<-as.numeric(unlist(deg_out.fp_c3)) #

#praw_fp.c1<-(sum(abs(deg_out.fp_c1) >= abs(BTN1centrality_degreeout_fp)) + 1) / (length(deg_out.fp_c1) + 1) #p  0.058
#praw_fp.c1
#

zscore<-net.zscore(BTN3Icentrality_degreeout_fd, deg_out.fp_c3)
zscore
net.metric.pvalue <- 2*pnorm(-abs(zscore))
net.metric.pvalue # 
#
######################################################################################

#colony 3 #weights null model 1 in-degree
Nperm=1000
null.model_l_in_n3 <- lapply(seq_len(Nperm), function(x){  
  randomnet <- rg_reshuffling_w(BTNC3Itotaligs_b_tnet, option="links") 
})

deg_in_links <- function(x){
  degree_w(x,measure=c("output"), type="in", alpha=0.5)
}

deg_in_nullmodel_l_n3<-lapply(null.model_l_in_n3, deg_in_weights)

#####node degree in #weights null model 1

deg_in_nullmodel_l_n3_WL<-lapply(deg_in_nullmodel_l_n3, function(x) x[1,2]) #WL

deg_in.wl_c3<-as.data.frame(unlist(deg_in_nullmodel_l_n3_WL))
deg_in.wl_c3<-as.numeric(unlist(deg_in.wl_c3)) #

#praw_wl.c1<-(sum(abs(deg_in.wl_c1) >= abs(BTN1centrality_degreein_wl)) + 1) / (length(deg_in.wl_c1) + 1) #p 0.06
#praw_wl.c1

zscore<-net.zscore(BTN3Icentrality_degreein_wl, deg_in.wl_c3)
zscore
net.metric.pvalue <- 2*pnorm(-abs(zscore))
net.metric.pvalue # 

#
deg_in_nullmodel_l_n3_SG<-lapply(deg_in_nullmodel_l_n3, function(x) x[2,2]) #SG

deg_in.sg_c3<-as.data.frame(unlist(deg_in_nullmodel_l_n3_SG))
deg_in.sg_c3<-as.numeric(unlist(deg_in.sg_c3)) #

#praw_sg.c1<-(sum(abs(deg_in.sg_c1) >= abs(BTN1centrality_degreein_sg)) + 1) / (length(deg_in.sg_c1) + 1) #p 0.08
#praw_sg.c1

zscore<-net.zscore(BTN3Icentrality_degreein_sg, deg_in.sg_c3)
zscore
net.metric.pvalue <- 2*pnorm(-abs(zscore))
net.metric.pvalue # 

#
deg_in_nullmodel_l_n3_AT<-lapply(deg_in_nullmodel_l_n3, function(x) x[3,2]) #AT

deg_in.at_c3<-as.data.frame(unlist(deg_in_nullmodel_l_n3_AT))
deg_in.at_c3<-as.numeric(unlist(deg_in.at_c3)) #

#praw_at.c1<-(sum(abs(deg_in.at_c1) >= abs(BTN1centrality_degreein_at)) + 1) / (length(deg_in.at_c1) + 1) #p 0.96
#praw_at.c1

zscore<-net.zscore(BTN3Icentrality_degreein_at, deg_in.at_c3)
zscore
net.metric.pvalue <- 2*pnorm(-abs(zscore))
net.metric.pvalue # 

#
deg_in_nullmodel_l_n3_CF<-lapply(deg_in_nullmodel_l_n3, function(x) x[4,2]) #CF

deg_in.cf_c3<-as.data.frame(unlist(deg_in_nullmodel_l_n3_CF))
deg_in.cf_c3<-as.numeric(unlist(deg_in.g_c3)) #

#praw_g.c1<-(sum(abs(deg_in.g_c1) >= abs(BTN1centrality_degreein_g)) + 1) / (length(deg_in.g_c1) + 1) #p 0.39
#praw_g.c1

zscore<-net.zscore(BTN3Icentrality_degreein_cf, deg_in.cf_c3)
zscore
net.metric.pvalue <- 2*pnorm(-abs(zscore))
net.metric.pvalue # 

#
deg_in_nullmodel_l_n3_BC<-lapply(deg_in_nullmodel_l_n3, function(x) x[5,2]) #BC

deg_in.bc_c3<-as.data.frame(unlist(deg_in_nullmodel_l_n3_BC))
deg_in.bc_c3<-as.numeric(unlist(deg_in.bc_c3)) #

#praw_bc.c1<-(sum(abs(deg_in.bc_c1) >= abs(BTN1centrality_degreein_bc)) + 1) / (length(deg_in.bc_c1) + 1) #p 0.38
#praw_bc.c1

zscore<-net.zscore(BTN3Icentrality_degreein_bc, deg_in.bc_c3)
zscore
net.metric.pvalue <- 2*pnorm(-abs(zscore))
net.metric.pvalue # 

#
deg_in_nullmodel_l_n3_FP<-lapply(deg_in_nullmodel_l_n3, function(x) x[6,2]) #FP

deg_in.fp_c3<-as.data.frame(unlist(deg_in_nullmodel_l_n3_FP))
deg_in.fp_c3<-as.numeric(unlist(deg_in.fp_c3)) #

#praw_fp.c1<-(sum(abs(deg_in.fp_c1) >= abs(BTN1centrality_degreein_fp)) + 1) / (length(deg_in.fp_c1) + 1) #p 0.88
#praw_fp.c1

zscore<-net.zscore(BTN3Icentrality_degreein_fp, deg_in.fp_c3)
zscore
net.metric.pvalue <- 2*pnorm(-abs(zscore))
net.metric.pvalue #

#
deg_in_nullmodel_l_n3_FD<-lapply(deg_in_nullmodel_l_n3, function(x) x[7,2]) #FD

deg_in.fd_c3<-as.data.frame(unlist(deg_in_nullmodel_l_n3_FD))
deg_in.fd_c3<-as.numeric(unlist(deg_in.fd_c3)) #

#praw_fp.c1<-(sum(abs(deg_in.fp_c1) >= abs(BTN1centrality_degreein_fp)) + 1) / (length(deg_in.fp_c1) + 1) #p 0.88
#praw_fp.c1

zscore<-net.zscore(BTN3Icentrality_degreein_fd, deg_in.fd_c3)
zscore
net.metric.pvalue <- 2*pnorm(-abs(zscore))
net.metric.pvalue #

##############################################################################################################################
```
