## Appendix, as digital supplementary material to this article for "Worker-behavior and behavior-behavior interaction networks in the trap-jaw ant *Odontomachus chelifer* (Latreille, 1802) (Hymenoptera: Formicidae)": Script S14.R

```
#TTNs_modularity
library(ggplot2)
TTNs_modularity<-read.table("Data S23.txt", header = TRUE) #Data S23

ggplot(TTNs_modularity, aes(x=colonies, y=Q_value, fill=Data)) + 
  geom_bar(stat="identity", color="black", 
           position=position_dodge()) +
  geom_errorbar(aes(ymin=Q_value-sd_data, ymax=Q_value+sd_data), width=.2,
                position=position_dodge(.9)) + theme_classic()

#####
TTNs_nested_Fc<-read.table("Data S24.txt", header = TRUE)

ggplot(TTNs_nested_Fc, aes(x=colonies, y=UNODF_value, fill=Data)) + 
  geom_bar(stat="identity", color="black", 
           position=position_dodge()) +
  geom_errorbar(aes(ymin=UNODF_value-sd_data, ymax=UNODF_value+sd_data), width=.2,
                position=position_dodge(.9)) + theme_classic()

#TTNs_nested_Fr

TTNs_nested_Fr<-read.table("Data S25.txt", header = TRUE)

ggplot(TTNs_nested_Fr, aes(x=colonies, y=UNODF_value, fill=Data)) + 
  geom_bar(stat="identity", color="black", 
           position=position_dodge()) +
  geom_errorbar(aes(ymin=UNODF_value-sd_data, ymax=UNODF_value+sd_data), width=.2,
                position=position_dodge(.9)) + theme_classic()

    
#TTNs_0.01_nested_Fc
TTNs_0.01_nested_Fc<-read.table("Data S26.txt", header = TRUE)

ggplot(TTNs_0.01_nested_Fc, aes(x=colonies, y=UNODF_value, fill=Data)) + 
  geom_bar(stat="identity", color="black", 
           position=position_dodge()) +
  geom_errorbar(aes(ymin=UNODF_value-sd_data, ymax=UNODF_value+sd_data), width=.2,
                position=position_dodge(.9)) + theme_classic()

#TTNs_0.01_nested_Fr
TTNs_0.01_nested_Fr<-read.table("Data S26.txt", header = TRUE)

ggplot(TTNs_0.01_nested_Fr, aes(x=colonies, y=UNODF_value, fill=Data)) + 
  geom_bar(stat="identity", color="black", 
           position=position_dodge()) +
  geom_errorbar(aes(ymin=UNODF_value-sd_data, ymax=UNODF_value+sd_data), width=.2,
                position=position_dodge(.9)) + theme_classic()
```
