## Appendix, as digital supplementary material to this article for "Worker-behavior and behavior-behavior interaction networks in the trap-jaw ant *Odontomachus chelifer* (Latreille, 1802) (Hymenoptera: Formicidae)": Script S15.R

```
#modularity nulls models and p-values
#load first the BTN 1; BTN2 and BTN3 files
directed_undirected <- function(x){
  as.undirected(x, mode = c("collapse"), edge.attr.comb = "sum")
}

modular <- function(x){
  return(cluster_louvain(x)$modularity)
}

###############################################################################
#total 1 modularity null model 1 weight (w)
null.model_w_n1_igraph_mod<-lapply(null.model_w_n1_igraph, directed_undirected)
null.model_w_n1_igraph_mod<-lapply(null.model_w_n1_igraph_mod, modular)

null.model_w_n1_igraph_mod<-as.data.frame(unlist(null.model_w_n1_igraph_mod))
null.model_w_n1_igraph_mod<-as.numeric(unlist(null.model_w_n1_igraph_mod))


net.zscore = function(obsval, nullval) {
  (obsval - mean(nullval))/sd(nullval)  
} 

zscore<-net.zscore(BTN1mod$modularity, null.model_w_n1_igraph_mod)
zscore
net.metric.pvalue <- 2*pnorm(-abs(zscore))
net.metric.pvalue # p 0.08
#


#total 1 modularity null model 2 links (l)
null.model_l_n1_igraph_mod<-lapply(null.model_l_n1_igraph, directed_undirected)
null.model_l_n1_igraph_mod<-lapply(null.model_l_n1_igraph_mod, modular)

null.model_l_n1_igraph_mod<-as.data.frame(unlist(null.model_l_n1_igraph_mod))
null.model_l_n1_igraph_mod<-as.numeric(unlist(null.model_l_n1_igraph_mod))

zscore<-net.zscore(BTN1mod$modularity, null.model_l_n1_igraph_mod)
zscore
net.metric.pvalue <- 2*pnorm(-abs(zscore))
net.metric.pvalue # p 0.24
#

##############################################################################
#total 2 modularity null model 1 weight (w)
null.model_w_n2_igraph_mod<-lapply(null.model_w_n2_igraph, directed_undirected)
null.model_w_n2_igraph_mod<-lapply(null.model_w_n2_igraph_mod, modular)

null.model_w_n2_igraph_mod<-as.data.frame(unlist(null.model_w_n2_igraph_mod))
null.model_w_n2_igraph_mod<-as.numeric(unlist(null.model_w_n2_igraph_mod))

zscore<-net.zscore(BTN2mod$modularity,null.model_w_n2_igraph_mod)
zscore
net.metric.pvalue <- 2*pnorm(-abs(zscore))
net.metric.pvalue # 0.07
#

#total 2 modularity null model 2 links (l)
null.model_l_n2_igraph_mod<-lapply(null.model_l_n2_igraph, directed_undirected)
null.model_l_n2_igraph_mod<-lapply(null.model_l_n2_igraph_mod, modular)

null.model_l_n2_igraph_mod<-as.data.frame(unlist(null.model_l_n2_igraph_mod))
null.model_l_n2_igraph_mod<-as.numeric(unlist(null.model_l_n2_igraph_mod))

zscore<-net.zscore(BTN2mod$modularity,null.model_l_n2_igraph_mod)
zscore
net.metric.pvalue <- 2*pnorm(-abs(zscore))
net.metric.pvalue # 0.001

###################################################################################
#total 3 modularity null model 1 weight (w)
null.model_w_n3_igraph_mod<-lapply(null.model_w_n3_igraph, directed_undirected)
null.model_w_n3_igraph_mod<-lapply(null.model_w_n3_igraph_mod, modular)

null.model_w_n3_igraph_mod<-as.data.frame(unlist(null.model_w_n3_igraph_mod))
null.model_w_n3_igraph_mod<-as.numeric(unlist(null.model_w_n3_igraph_mod))

zscore<-net.zscore(BTN3mod$modularity,null.model_w_n3_igraph_mod)
zscore
net.metric.pvalue <- 2*pnorm(-abs(zscore))
net.metric.pvalue # 0.38


#total 3 modularity null model 2 links (l)
null.model_l_n3_igraph_mod<-lapply(null.model_l_n3_igraph, directed_undirected)
null.model_l_n3_igraph_mod<-lapply(null.model_l_n3_igraph_mod, modular)

null.model_l_n3_igraph_mod<-as.data.frame(unlist(null.model_l_n3_igraph_mod))
null.model_l_n3_igraph_mod<-as.numeric(unlist(null.model_l_n3_igraph_mod))

zscore<-net.zscore(BTN3mod$modularity,null.model_l_n3_igraph_mod)
zscore
net.metric.pvalue <- 2*pnorm(-abs(zscore))
net.metric.pvalue # 0.07
```
